# Benchmarking resting state fMRI connectivity pipelines for classification: Robust accuracy despite processing variability in cross-site eye state prediction

**DOI:** 10.1101/2025.10.20.683049

**Authors:** Tatiana Medvedeva, Irina Knyazeva, Ruslan Masharipov, Alexander Korotkov, Denis Cherednichenko, Maxim Kireev

**Affiliations:** N. P. Bechtereva Institute of the Human Brain, Russian Academy of Sciences, St. Petersburg, Russia

**Keywords:** fMRI, Resting state, Functional connectivity, Denoising, Machine learning

## Abstract

The rapid evolution of machine learning (ML) methods has yielded promising results in human brain neuroscience. However, the reproducibility of ML applications in neuroimaging remains limited, challenging the generalizability of inferences to broader populations. In addition to the inherent variability of the brain activity (both in healthy and pathological states), poor reproducibility is further enhanced by inconsistencies in data preprocessing techniques and methods for calculating functional connectivity (FC), which are used as parameters for brain state classification. To systematically assess the impact of abovementioned factors on ML applications to fMRI data, we benchmarked a comprehensive set of FC analysis pipelines for the classification task between fMRI data recorded in two fundamentally different states: eyes open and eyes closed. In contrast to studies involving heterogeneous clinical populations or using complex cognitive tasks, our controlled experimental design – based on two independent datasets of healthy participants collected in different laboratories – minimizes variability related to a task design or pathological brain states. Classification accuracy and reproducibility were compared for 256 distinct FC analysis pipelines, covering common preprocessing approaches, brain parcellation schemes, and connectivity metrics. Notably, we employed two ways of validation: a direct cross-site validation strategy – when a model was trained on one site and tested on another, and few-shot domain adaptation – when a few samples of testing site were added to the train set. Despite the substantial variability in pipeline configurations, we observed consistently high classification accuracy (∼80%), confirming that FC-based models can robustly discriminate between well-defined brain states (eye conditions) across different acquisition sites. Best results both in terms of classification accuracy and stability were observed using Pearson correlation and tangent space parametrization as FC, Brainnetome as atlas, and confound regression strategies based on the CompCor method. These findings highlight the resilience of rs-fMRI FC-derived characteristics to methodological variation and support their utility in the discovery of biomarkers, particularly in settings that involve stable and reproducible brain states.

## Introduction

Despite recent advances in the application of machine learning (ML) to classify brain states, conditions, and patient groups using resting-state fMRI (rs-fMRI) functional connectivity (FC) parameters, reproducibility remains a major challenge. This limitation significantly prevents the development of reliable neuromarkers, largely due to methodological variability introduced by diverse data processing pipelines.

From rs-fMRI data preprocessing steps to choices in brain anatomical parcellation and functional connectivity metrics, decisions made during fMRI analysis can strongly influence the results obtained using ML models (Noble et al., 2019). This variability not only complicates the comparison of findings across studies, but also undermines confidence in the generalizability of reported results, limiting progress in understanding brain function using ML approaches, and developing reliable ML-based neuroimaging biomarkers for diagnostic purposes. Therefore, achieving high consistency and reproducibility in research utilizing rs-fMRI functional connectivity for ML-based classification is challenging due to a combination of physiological and technical factors. (Peterson et al., 2025) One of the most problematic factors is head movement during fMRI scanning, which can seriously corrupt the results of a study. (Friston et al., 1996) Additionally, physiological sources of variability include heart rate and respiration, state-dependent changes, which also introduce noise into the fMRI signal. (Murphy et al., 2013)

Recent efforts have highlighted the need for systematic evaluations of how methodological decisions influence the conclusions drawn from fMRI data. Methodological changes may be introduced in all steps of the analysis with limitless number of choices, starting with raw MRI data preprocessing and confound regression of BOLD signal (Ciric et al., 2017; Parkes et al., 2018; Satterthwaite et al., 2013), ending with the choice of functional connectivity measures (Z.-Q. Liu et al., 2025; Luppi et al., 2024) and machine learning models (Dadi et al., 2019; Khosla et al., 2018).

Another source of variation in such studies is the choice of an fMRI dataset, according to which fMRI processing pipelines are tested. Some of the most investigated resting state datasets are ABIDE (Di Martino et al., 2014), HCP (Van Essen et al., 2013), ADNI (Mueller et al., 2005), UK Biobank (Mansour et al., 2023). In most studies evaluating classification performance of ML methods using rs-fMRI data, experimental designs typically assume inter-subject comparisons.

They often involve contrasting patients to controls (Eslami et al., 2019; Riaz et al., 2020; Segato et al., 2020), or distinct clinical populations (Qin et al., 2022). However, such designs inherently lack ground truth and introduce considerable variability due to individual differences that are difficult to control. This uncontrolled inter-subject variability can significantly compromise the interpretability and reproducibility of classification results, obscuring the true impact of methodological choices.

To address this limitation, the present study, which focuses on evaluating the robustness of rs-fMRI-based classification pipelines, adopts a within-subject design. By comparing brain states within the same individuals, we effectively control for inter-subject variability, allowing for a more precise assessment of methodological influences. Specifically, we utilize a well-established experimental paradigm that contrasts the eyes open and fixated (EO-F) and eyes closed (EC) conditions during resting-state — two physiological states known to elicit distinct neural activity patterns, as first demonstrated in classical studies (Berger, 1929; Gusnard & Raichle, 2001; Marx et al., 2004) and consistently reported in subsequent literature (Agcaoglu et al., 2020; Costumero et al., 2020; DeRamus et al., 2021; Han et al., 2023; Patriat et al., 2013; Weng et al., 2020). This binary classification task provides a controlled framework for evaluating classification performance and methodological reproducibility.

Opening and closing the eyes triggers activity changes in many brain areas. (Agcaoglu et al., 2019; Costumero et al., 2020; Gusnard & Raichle, 2001; Marx et al., 2004; Patriat et al., 2013; Weng et al., 2020; Yan et al., 2009) For instance, (Agcaoglu et al., 2019) demonstrated a functional connectivity decrease in visual networks when subjects closed their eyes, presumably due to visual input removal. Conversely, opening the eyes restored this visual network activity, highlighting the brain’s quick adjustment to visual changes. It was also found that for auditory and sensorimotor brain networks, there were more functionally connected brain regions during EC than during EO. Furthermore, the default mode network (DMN), a network of brain regions that are active during resting-state, is reported in numerous studies on differences between EO/EC states (Gusnard & Raichle, 2001; Marx et al., 2004; Yan et al., 2009). The medial prefrontal cortex (mPFC) and the posterior cingulate cortex (PCC) are core regions of the default mode network: these areas are associated with mind wandering and self-referential thinking, cognitive states often occurring when an individual is not focused on tasks dealing with external stimuli. (Mason et al., 2007) By using seed-based between group analysis it was demonstrated that in the EC group primary visual cortex had higher connectivity with the DMN and sensorimotor network, whilst in the EO group higher connectivity was observed with the salience network. (Costumero et al., 2020) However, (Yan et al., 2009) showed that the DMN had stronger connectivity for EO than for EC. Moreover, (Patriat et al., 2013) reported very small differences in functional connectivity maps between EO, EO-F and EC conditions. In (Weng et al., 2020) review the main findings of the EC/EO comparisons were summarized as differences in functional connectivity in the resting state observed in the visual cortex, the sensory cortex and supplementary motor area, the thalamus, frontal eye fields and the visual, sensorimotor, default mode, attention networks as well. Therefore, the EO and EC conditions represent suitable candidates for evaluating the effectiveness of classification methods in distinguishing between different brain states using FC fMRI data.

Alongside physiological variability, technical sources of variation further complicate the analysis. Functional connectome-based classification refers to the process of leveraging patterns in functional connectivity data to differentiate among multiple subject states or conditions. (Khosla et al., 2018) Despite advantages of this approach, different analysis workflows may lead to drastically different results. (Dadi et al., 2019) In Figure 1, a common pipeline of creating functional connectome-based classification models is shown. Basically, every step of this pipeline has a great number of parameters to vary:

1. Data acquisition: acquisition parameters variation, effects of scanner and scanned population (e.g., multi-site datasets). (Bari et al., 2019)
2. Functional and structural data minimal preprocessing: variations in raw data processing steps (such as segmentation, motion correction and registration) have a significant impact on results, as shown by (Li et al., 2024).
3. BOLD signal denoising strategies: choosing a non-neural signals removal technique among other methods (Ciric et al., 2017; Parkes et al., 2018; Satterthwaite et al., 2013)
4. Brain parcellation: variability of data-driven algorithms (clustering, ICA, etc.) or predefined atlases (structural or functional, with bigger or smaller number of ROIs). (Arslan et al., 2018)
5. Functional connectivity estimation: diversity of algorithms that measure similarity between regions’ activation. (Z.-Q. Liu et al., 2025; Peterson et al., 2025)
6. Diversity of Machine learning (ML) models to solve a classification task. (Dadi et al., 2019; Khosla et al., 2018)

**Figure 1.**
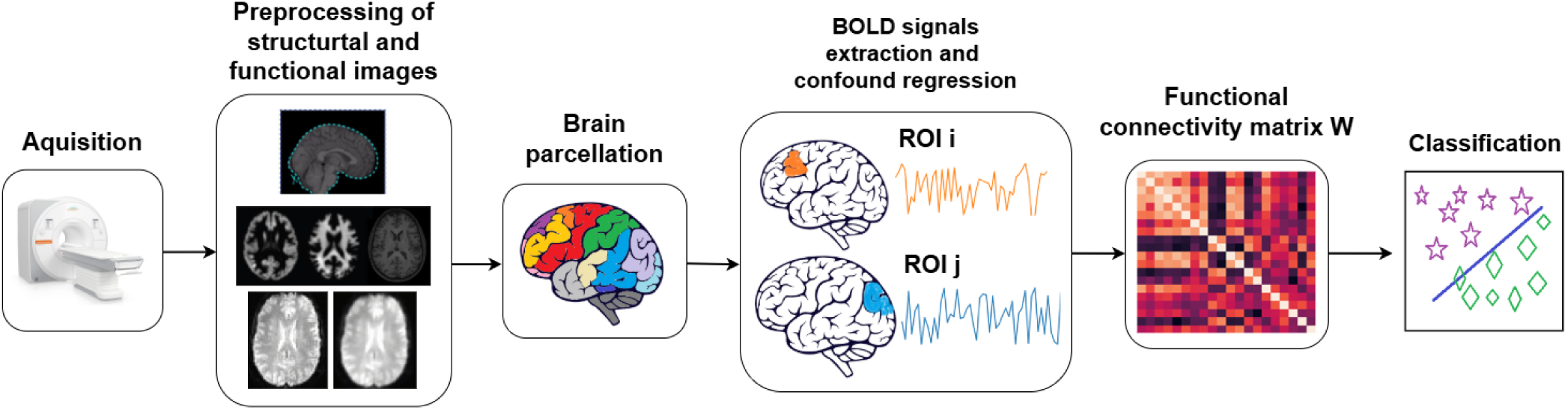
Pipeline of creating a functional connectome based predictive model *1)* Data collection stage. 2) Preprocessing of functional and structural images, which includes image time correction, white and gray matter segmentation, motion correction, comparison of structural and functional images, image smoothing. 3) Brain parcellation is the grouping of voxels into larger regions of interest. 4) Signal extraction and confound regression – noise associated with head movement, breathing, blood flow, etc. is removed from the BOLD signal of the regions of interest. 5) Functional connectivity matrix – each element of the matrix corresponds to a measure of functional connectivity between the signals of the two regions of interest. 6) The functional connectivity matrix is used to solve a classification problem.

Considering all variability sources can be overwhelming due to the numerous complex factors involved at all stages of fMRI data collection and analysis. In this paper, we fixated the last step of the pipeline and used the simplest yet effective classification model – Logistic Regression (Popov et al., 2024), while focusing on benchmarking four critical data processing factors, including: 1) Data acquisition variability – by using 2 datasets acquired in different circumstances; 2) BOLD signal denoising variability – by comparing 16 denoising strategies; 3) Brain parcellation variability – by using 4 atlases; 4) Connectivity estimation variability – by comparing 4 functional connectivity measures.

In total, we evaluated 256 distinct FC pipelines, systematically varying preprocessing strategies, anatomical brain parcellation schemes, and functional connectivity measures. We employed two complementary validation strategies. First, we conducted cross-site validation by training a ML model on data from one research site and testing it on another. Second, we used a few-shot domain adaptation setting, in which we augmented the training set with a small, controlled number of target-site samples to improve generalization to the remaining target-site participants. Together, these strategies help ensure that our findings reflect true methodological robustness rather than site-specific artifacts or overfitting. Although large sample sizes typically improve ML generalizability, fMRI datasets are often limited due to the complexity and cost of data collection. By leveraging two datasets (one smaller and one larger) we can assess whether predictive models generalize across different data regimes, thereby maximizing the utility of limited but valuable fMRI data.

Furthermore, recent discussions have highlighted the potential benefits of within-participant study designs for enhancing statistical power when neuroimaging data are used. (Marek et al., 2022; Spisak et al., 2023; Tervo-Clemmens et al., 2023) While our primary focus is on the pipeline robustness for state classification, we also investigate whether the structure of the training data – specifically, leveraging repeated measures from fewer subjects versus single measures from more subjects – impacts the performance of our classification models in distinguishing eye states.

Our results demonstrated that despite substantial differences in pipeline configurations, FC-based classification models sustainably achieve high accuracy (about 80% for best pipelines). This finding suggests that functional connectivity features are resilient to many forms of methodological variation when used to capture stable brain state differences. Nevertheless, we identified the most successful pipeline option that allowed us to achieve the highest classification accuracy. Interestingly, our analysis did not reveal a significant difference in model performance between these two experimental designs (within-subject versus between-subject). This suggests that in the terms of state classification (e.g., eyes-open vs. eyes-closed) both within- and between-subject setups can yield comparable predictive performance under appropriate modeling conditions. These results have important implications for the development of reproducible neuroimaging biomarkers, particularly in settings where brain states are expected to be consistent and clearly defined.

## Methods

### Data acquisition variability

Data acquisition (including scanner variability, acquisition parameters and sample bias) is a major source of variation that can significantly impact estimates of functional connectivity. For instance, variations in MR-scanner field strength can lead to differences in signal-to-noise ratio and spatial resolution, affecting the detection of functional connections. Even with identical imaging sequences and parameters, site-specific differences can still occur due to various physical factors. These include field inhomogeneity, configurations of transmit and receive coils, system stability, maintenance routines, scanner drift over time, and other potential variables (Bari et al., 2019). Moreover, we cannot deny the effect of the sample group. In particular, if we take clinical data as an example, disease factors may become confounding along with scanner-specific confounds in situations, when a site is focused on sampling specific types of disorder (Yamashita et al., 2019).

Benchmarking data acquisition variability allows us to assess how changes in acquisition protocols influence the robustness of predictive models. This is particularly crucial for multi-site studies, where harmonizing data across different scanners is an ongoing challenge. Data used in the present study was taken from 2 sources: 1) China, Beijing: Eyes Open Eyes Closed Study dataset (D. Liu et al., 2013); 2) Research data from the Institute of the Human Brain of the Russian Academy of Sciences (IHB RAS).

### BOLD signal denoising variability

Benchmarking denoising strategies is critical for understanding their impact on functional connectivity and avoiding potential confounds in the data. Denoising strategy might either under-correct or over-correct the data, leading to either residual noise contaminating the connectivity estimates or meaningful signals being mistakenly discarded. Estimating variability across different denoising strategies helps researchers find the right balance between preserving functional signals and eliminating noise, ensuring that the connectivity matrices accurately reflect brain network dynamics.

Denoising can significantly alter the connectivity structure, and consequently affect the performance of predictive models. Some techniques are commonly used among researchers (such as regression of motion parameters), others are controversial. For instance, global signal regression (GSR), which is regression of average signal across all voxels in the brain, is among those that are often discussed. The issue is that GSR may introduce poorly explained negative correlations in FC, leading to bias in connectivity estimates (Murphy & Fox, 2017). In the current study, we checked if there were notable differences in strategies that regress global signal out and those that did not.

We also employed CompCor (principal component-based noise correction) in our analysis (Behzadi et al., 2007). Two variants are available: aCompCor, or anatomical CompCor, obtains confound regressors from the principal components of the BOLD signal in white matter (WM) and cerebrospinal fluid (CSF), and tCompCor, or temporal CompCor, takes voxels with high temporal standard deviation.

In some sense, using both CompCor and GSR is excessive, since global signal takes into account time series of WM and CSF (e.g., in the CONN Toolbox they suggest using either CompCor or GSR (Whitfield-Gabrieli & Nieto-Castanon, 2012)). In fact, studies that aimed to compare confound regression strategies usually did not apply both GSR and CompCor simultaneously (e.g., (Ciric et al., 2017)). In the present study, we also compared both strategies with simultaneous CompCor and GSR, and only CompCor application to explore their influence on ML classification between analyzed brain states.

We also benchmarked ICA-AROMA strategies in this work. ICA-AROMA (Pruim et al., 2015) is an automated method designed to remove motion-related artifacts from fMRI data, without requiring manual component selection. It first decomposes the data into spatial independent components, then classifies each component as “motion-related” noise using heuristics such as spatial distribution (brain-edge and CSF coverage), high-frequency temporal characteristics, and correlation with volume-wise motion regressors; finally it regresses the associated temporal signals back into the data to produce a less motion-contaminated time series. We utilized fmripost-aroma^1^ implementation of ICA-AROMA. Two denoising outputs were generated: an “aggressive” and a “non-aggressive” AROMA variant. In the aggressive strategy, the full time series of all components classified as motion-related were regressed out from the BOLD signal, aiming to maximally reduce motion-induced variance but at the risk of over-removal of neural signal. In the non-aggressive version, only the portion of each noise component strictly attributable to motion artifacts was removed, while shared variance (potentially reflecting neural-related fluctuations) was preserved, making it a more conservative option. Since ICA-AROMA discards motion-related components only, for both variants we also added aCompCor regressors with 10 components to handle confounds associated with WM and CSF.

In total, we evaluated 16 denoising strategies (Table 1). Being an essential part of fMRI data processing, head motion parameters (HMP) were included in all strategies. The 24 HMP strategy included 6 motion parameters, 6 temporal derivatives, 6 quadratic terms, and 6 quadratic expansions of the derivatives of motion estimates (Friston et al., 1996). Here we also used the 12 HMP strategy, which included 6 motion estimates and their temporal derivatives. We used aCompCor with different number of components: by default we took 10 principal components calculated using a WM and CSF combined anatomical mask, resulting in fixed number of regressors for every subject; and aCompCor50, which included principal components that explain 50% of variance, calculated using a WM and CSF combined anatomical mask (number of regressors varied among subjects). For tCompCor, the number of regressors also differed from subject to subject. Finally, we added both aggressive and non-aggressive ICA-AROMA with aCompCor. Strategies 9-16 included 4 global signal regressors: the global signal, its derivative, its square, and the derivative of its square.

**Table 1.**
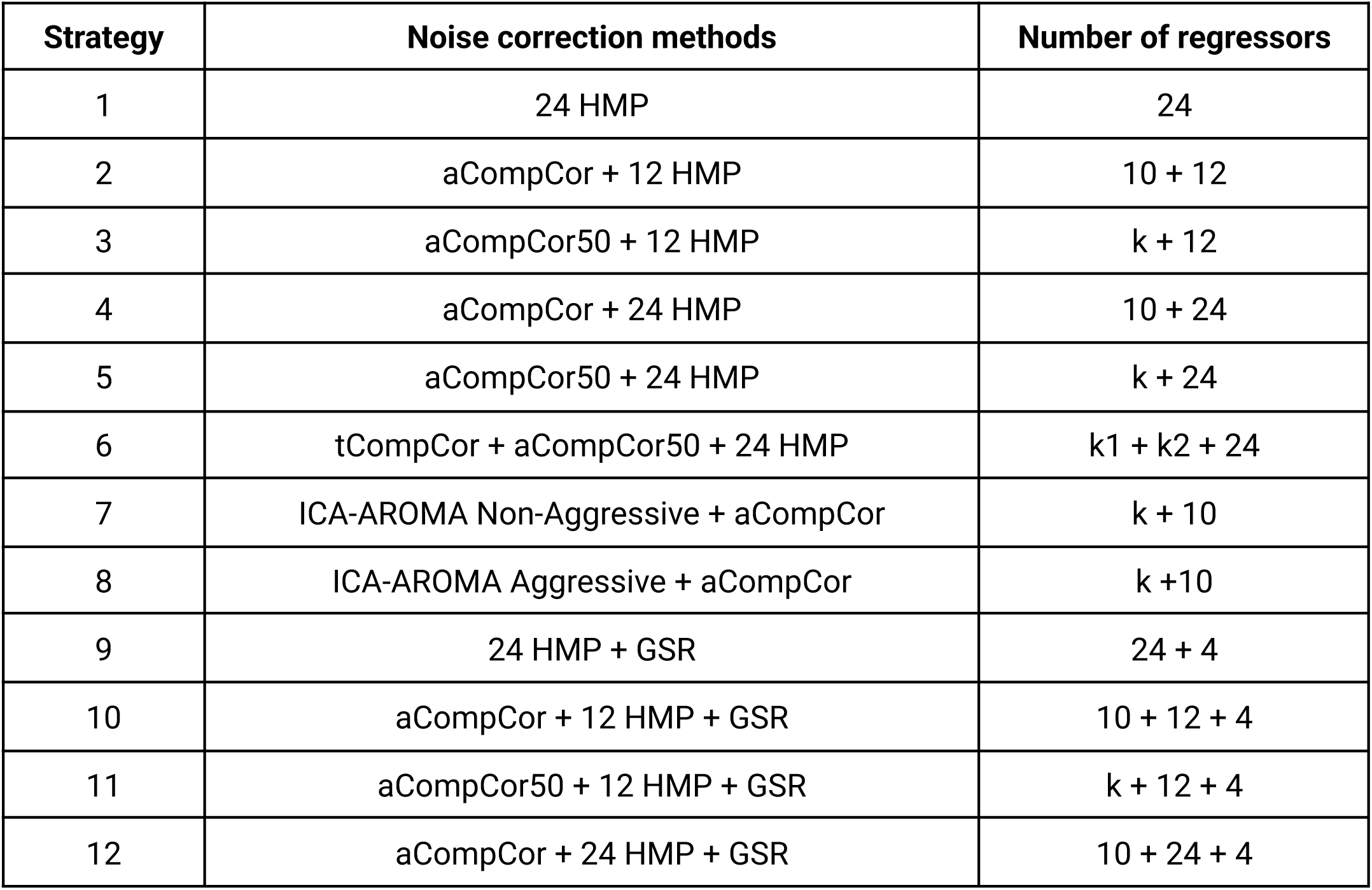
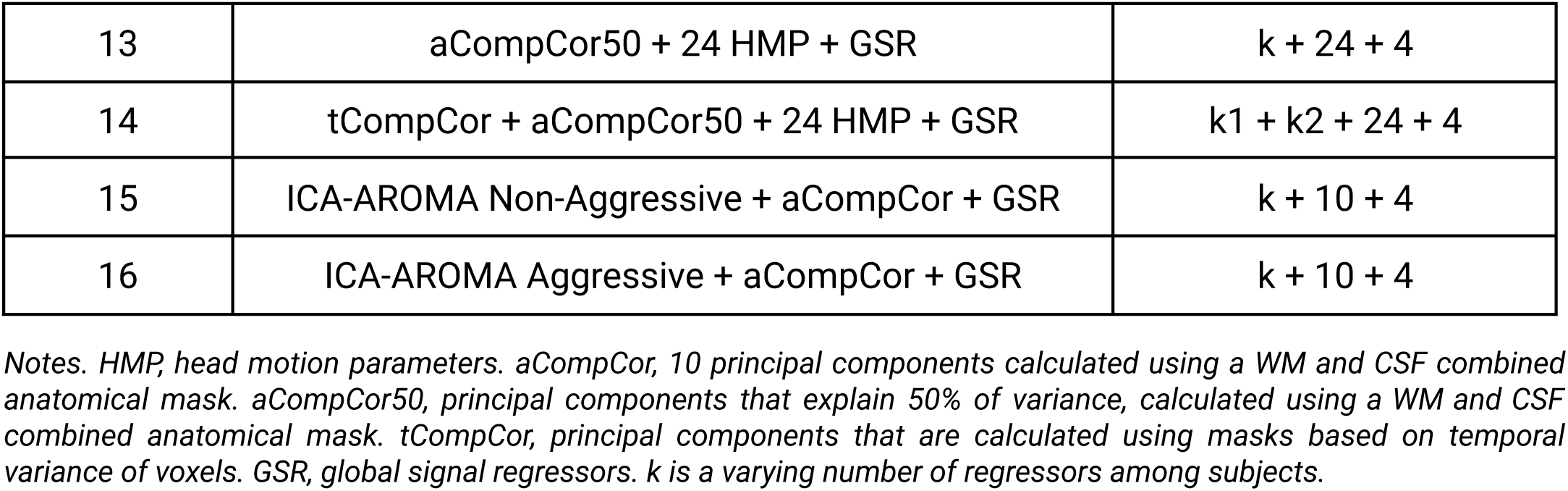
Characteristics of used pipelines.

| Strategy | Noise correction methods | Number of regressors |
| --- | --- | --- |
| 1 | 24 HMP | 24 |
| 2 | aCompCor + 12 HMP | 10 + 12 |
| 3 | aCompCor50 + 12 HMP | k + 12 |
| 4 | aCompCor + 24 HMP | 10 + 24 |
| 5 | aCompCor50 + 24 HMP | k + 24 |
| 6 | tCompCor + aCompCor50 + 24 HMP | k<sub>1</sub> + k<sub>2</sub> + 24 |
| 7 | ICA-AROMA Non-Aggressive + aCompCor | k + 10 |
| 8 | ICA-AROMA Aggressive + aCompCor | k + 10 |
| 9 | 24 HMP + GSR | 24 + 4 |
| 10 | aCompCor + 12 HMP + GSR | 10 + 12 + 4 |
| 11 | aCompCor50 + 12 HMP + GSR | k + 12 + 4 |
| 12 | aCompCor + 24 HMP + GSR | 10 + 24 + 4 |
| 13 | aCompCor50 + 24 HMP + GSR | $k + 24 + 4$ |
| 14 | tCompCor + aCompCor50 + 24 HMP + GSR | $k_1 + k_2 + 24 + 4$ |
| 15 | ICA-AROMA Non-Aggressive + aCompCor + GSR | $k + 10 + 4$ |
| 16 | ICA-AROMA Aggressive + aCompCor + GSR | $k + 10 + 4$ |

*Notes. HMP, head motion parameters. aCompCor, 10 principal components calculated using a WM and CSF combined anatomical mask. aCompCor50, principal components that explain 50% of variance, calculated using a WM and CSF combined anatomical mask. tCompCor, principal components that are calculated using masks based on temporal variance of voxels. GSR, global signal regressors. $k$ is a varying number of regressors among subjects.*

### Brain parcellation variability

Parcellation refers to dividing the brain into distinct anatomical regions of interest (ROIs), each representing a node in the brain network. It is a common practice to perform dimensionality reduction, since the dimensionality of fMRI data is quite high. The choice of parcellation scheme (based on brain atlases or data-driven) can lead to significant variability in the resulting connectivity matrices. Parcellation schemes affect the spatial resolution of the brain regions, which can drastically change the connectivity estimates. For instance, using a coarse atlas with larger regions may obscure fine-grained, functionally relevant connections, while overly fine atlases can introduce noise, leading to spurious connections and higher computation costs (Khan & Shang, 2025). Understanding how different atlases influence connectivity estimation helps researchers identify the optimal parcellation for their research questions.

Overall, all methods can be divided into 2 groups: 1) Pre-defined: provide coordinates in normalized brain space and are consistent across studies; 2) Data-driven: calculated (usually using clustering and linear decomposition algorithms) for specific dataset and cannot be used on other data.

The choice of parcellation approach also impacts the reproducibility of findings across studies. Usage of data-driven parcellations potentially complicates direct comparison across studies utilizing different datasets for parcellation (Badea et al., 2017). Since we analyze two different datasets and in order to make our research more reproducible, we utilized only predefined brain atlases. The four selected brain atlases encompass a range of parcellation strategies, spatial resolutions, and historical usage within the literature.

1. **Automated Anatomical Labeling (AAL)** (Tzourio-Mazoyer et al., 2002) consists of 116 regions encompassing both cortical and subcortical brain structures (we used the SPM12 version available in nilearn). AAL is a widely recognized anatomical brain atlas and serves as a historical standard in neuroimaging research. Its inclusion allows for direct comparison with a large body of existing literature that has utilized this foundational parcellation scheme, based primarily on macroscopic anatomical landmarks.
2. **Schaefer Atlas** (Schaefer et al., 2018) provides a parcellation based on functional connectivity derived from resting-state fMRI data. We selected the 200-region version, which focuses on cortical areas. Including this functional atlas complements the anatomical AAL parcellation by defining regions based on their functional properties and network organization, offering a different perspective on brain division based on distributed connectivity patterns.
3. **Brainnetome Atlas** (Fan et al., 2016) offers a parcellation that attempts to integrate both anatomical and functional brain connectivity information, resulting in 246 regions (210 cortical and 36 subcortical). Its inclusion allows us to examine results using a parcellation scheme that incorporates multiple levels of information and provides a medium-level resolution covering both cortical and subcortical brain areas, balancing the purely anatomical and purely functional approaches of AAL and Schaefer, respectively.
4. **HCPex Atlas** (Huang et al., 2022) is a high-resolution volumetric parcellation derived from the surface-based Human Connectome Project multimodal atlas (HCP-MMP v1.0), extended with detailed subcortical regions, totaling 426 parcells. This brain atlas represents a more recent and detailed approach, leveraging high-quality multimodal HCP data to define regions at a finer granularity. Its inclusion allows for the exploration of effects at a higher spatial resolution and provides a parcellation based on advanced mapping techniques.

### Connectivity estimation

Connectivity measures are used to quantify the functional relationships between different regions of the human brain. Variability in connectivity estimation arises from the choice of methods, such as Pearson correlation, tangent space parametrization, partial correlation. The selection of the appropriate connectivity measure can dramatically impact the outputs of FC-based machine learning models, as these measures capture different aspects of the brain’s functional organization (Z.-Q. Liu et al., 2025).

#### Pearson Correlation

This widely-used measure captures linear relationships between time series of different brain regions, it is simple and computationally effective. However, Pearson correlation is limited in its ability to account for indirect interactions between regions and nonlinear dependencies, which can result in oversimplification of the complex brain networks and overestimation of connectivity (Peterson et al., 2025).

#### Tangent space parametrization

Tangent space parametrization extends Pearson correlation by projecting the functional connectivity matrix onto a tangent space, making it more sensitive to individual variability, while preserving commonalities across subjects. This method is especially useful when we need to estimate the differences between subjects more precisely. Moreover, it is robust to noise and inter-subject variability. In predictive modeling, tangent space parametrization has been shown to improve classification accuracy and sensitivity to individual differences in brain connectivity profiles, providing a more nuanced representation of brain function (Abbas et al., 2023; Dadi et al., 2019).

#### Partial Correlation

Partial correlation accounts for the direct interaction between two brain regions, while controlling for the influence of other regions. This method is essential in connectivity estimation as it reveals the unique relationships between brain regions that are not mediated by other areas, which is crucial for identifying specific pathways that may contribute to behavior or clinical outcomes. Ignoring such direct connections might obscure important predictive information (Peterson et al., 2025).

**Regularized Partial Correlation**, or Graphical Lasso, or glasso. Partial correlation estimation may be extremely overfitted to the data, consequently it is less reliable and stable to retest tests. To lower overfitting, some researchers suggest using regularized partial correlation estimation (Peterson et al., 2025). Graphical lasso is a partial correlation estimation with *L1*-regularization.

In this work, we used nilearn’s implementation of Pearson correlation, partial correlation and tangent space parametrization. Code for regularized partial correlation was adapted from (Peterson et al., 2025).

### Data

IHB RAS data consists of 2 resting state sessions with eyes closed (EС), eyes open with fixation (EO-F) for 84 healthy subjects. Acquisition of MR images was performed with 3T Philips Achieva scanner. EPI scan parameters: 32 axial slices, 120 volumes, repetition time (TR) = 2500 ms, echo time (TE) = 35 ms, flip angle= 90°. Structural images acquisition parameters: 130 axial slices, slice thickness = 0.94 mm, TR = 2500 ms, TE = 3.1 ms, flip angle = 30°.

The China dataset^2^ included 48 healthy controls and each subject has 3 rs-fMRI scans. During the first scan, participants were instructed to rest with their eyes closed, whereas the 2nd and 3rd sessions were either with closed or open eyes. Sessions were recorded sequentially. The MR images were acquired using a SIEMENS TRIO 3-Tesla scanner. EPI scan parameters: 33 axial slices, 240 volumes, TR = 2000 ms, TE = 30 ms, flip angle= 90°. T1-weighted MPRAGE parameters: 128 sagittal slices, slice thickness = 1.33 mm, TR = 2530 ms, TE = 3.39 ms, flip angle = 7° (D. Liu et al., 2013).

Raw fMRI data preprocessing for both datasets was performed using fMRIPrep 23.2.0. (Esteban et al., 2019). Anatomical processing included skull stripping (for IHB RAS), spatial normalization according to MNI standard template (2 spaces used: MNI152NLin6Asym for ICA-AROMA, MNI152NLin2009cAsym for the rest strategies) , and tissue segmentation. For functional data, slice timing correction, motion correction, normalization and co-registration to the corresponding anatomical image were performed. Lastly, data was smoothed using a Gaussian filter with 4 mm kernel. Additionally, confounding time-series (e.g., motion parameters and physiological noise estimates) were extracted for subsequent analyses.

Functional scans tend to have areas with extremely uneven signal that is contaminated by signals of non-neural origin (e.g., breathing in the nasal sinuses area) or caused by MRI scanner characteristics. The elimination of such areas is performed with several steps. Firstly, for every subject a new brain mask is formed from voxels, the signal level in which is not lower than some part of the global signal (we used 0.6). Then the mean mask for all subjects is formed from the new masks. To binarize the mean mask, we used a threshold to accept voxels that presented in new masks of at least 50% of all subjects.

Further, we used the mean brain mask for all subjects to get the coverage of parcels for each brain atlas. For this, we calculated the number of voxels that fall into each parcel. We rejected parcels that had less than 10% of voxels, according to the mean brain mask. Resulting number of ROI for each atlas: AAL – 106 ROIs of 116, Schaefer – 182 ROIs of 200, Brainnetome – 220 ROIs of 246, HCPex – 379 ROIs of 426.

Next, to extract the BOLD time-series, we used the nilearn^3^ library for Python. For each atlas, we extracted time-series with denoising strategies described in Table 1. Confounding time-series used for denoising were calculated in fMRIPrep: framewise displacement (FD), mean relative RMS (root-mean-square) displacement, global signal, head motion parameters, aCompCor and tCompCor estimates. We also added discrete cosines transformation basis regressors to handle low-frequency signal drifts in all strategies.

We did not discard any subjects due to motion artifacts. The mean framewise displacement for the IHB RAS dataset was 0.154 mm for the EO condition and 0.172 mm for the EC condition. For the China dataset, the corresponding values were 0.094 mm (EO) and 0.106 mm (EC).

### Functional connectivity quality metrics

To evaluate and compare all the varieties of pipelines discussed, we used a number of metrics that assess data quality after preprocessing, residual motion artifacts, test-retest reliability and the effectiveness of the classification model.

#### QC-FC correlations

The Quality Control-Functional Connectivity (QC-FC) metric is a key benchmark used to assess the effectiveness of denoising strategies in resting-state fMRI studies. It provides a direct measure of the relationship between motion-related artifacts and functional connectivity, unlike other quality metrics (e.g., framewise displacement, FD) that focus on general signal quality without directly addressing functional connectivity measures (Ciric et al., 2017; Parkes et al., 2018). QC-FC is calculated by examining the correlation between motion metrics and the functional connectivity strength for each edge in the connectivity matrix across subjects (Power et al., 2015). In the present study, we used the mean relative RMS displacement as a motion metric, which refers to the average frame-to-frame motion of the brain (Jenkinson et al., 2002). High mean relative RMS displacement suggests significant motion artifacts that can corrupt results. Consequently, high QC-FC correlations indicate that motion artifacts are still persistent in the data and influence FC estimates, suggesting a need for further refinement of the denoising process.

#### Similarity between denoising strategies

To assess similarity between denoising strategies, we calculate Pearson correlation between functional connectivity matrices (vectorized upper triangle) derived using different denoising strategies. In particular, for each person, we took connectivity matrices in two states and in 16 versions (a total of 32 matrices for one person). Then, within one subject, the correlation between flattened FC matrices is calculated. As a result, we get a correlation matrix, where each element represents the similarity between functional connectivity weights for two states and 16 strategies. Then we get a mean matrix across all subjects. This procedure is repeated for all means of variability separately. Using this metric, we can assess the divergence of the resulting FC weights among selected pipelines.

#### Intraclass Correlation Coefficient

The Intraclass Correlation Coefficient (ICC) is an established reliability measure in functional connectivity research, primarily assessing if the variability in a dataset was caused by stable between-subject differences or random and within-subject fluctuations (Shrout & Fleiss, 1979). The ICC score is especially valuable in connectivity studies as it can evaluate consistency at an individual level, which differentiates it from other metrics that often aggregate measures across subjects (Fiecas et al., 2013; Mahadevan et al., 2021). ICC values range from 0 to 1, with higher values indicating greater stability of measurements across repeated assessments.

Depending on the sources of error incorporated in the design, different forms of ICC are available. In this study, we focused on three commonly used ICC formulations that differ in their underlying assumptions about raters. Our primary reliability metric was ICC(3,1), corresponding to a two-way mixed-effects model, consistency, and single rater. In this formulation, subjects are treated as random effects, whereas the “rater” (i.e. pipeline) is treated as a fixed effect, and the ICC reflects the consistency of subject rankings across measurements rather than their absolute equality. ICC(3,1) is therefore appropriate when the specific measurement conditions used in the study are of primary interest and not intended to generalize to other, unobserved raters or conditions.

We also report ICC(2,1), defined as a two-way random-effects, absolute-agreement, single-measure model. Here, both subjects and raters (pipelines in our case) are modeled as random effects, and the ICC quantifies the degree to which repeated measurements on the same subject are in absolute agreement, allowing generalization to a broader population of similar raters. Finally, we include ICC(1,1), a one-way random-effects, absolute-agreement, single-measure formulation, which models variability across subjects with a single random effect and assumes that each subject is rated by a randomly selected rater. ICC(1,1) provides a more basic estimate of reliability when only a single source of random variance is explicitly modeled.

One limitation of ICC in FC analyses is its sensitivity to sparse connectivity methods, which often produce numerous edges with near-zero weights for all subjects. Since BMS and WMS for these “null” edges both tend to zero, the ICC values for these edges can be biased downward, clustering around zero. Consequently, using ICC across all edges in a sparse connectivity model may underrepresent reliability. To mitigate this, researchers typically exclude edges that consistently show zero weights and instead focus on those with systematic, non-zero values across subjects. (Peterson et al., 2025) This approach avoids the penalty of sparse models, enabling a more accurate assessment of edge-level reliability, where between-subject variability is present.

Given these considerations, in our study, we applied a masking strategy to focus on edges with meaningful between-subject variability, as suggested by (Peterson et al., 2025). Specifically, we excluded edges with near-zero correlation values, as these typically reflect noise rather than true connectivity. To ensure the ICC reliably captured relevant variability, we retained only edges in the 95th percentile of the group-averaged connectivity matrix for each functional connectivity measure. This approach minimized the influence of null edges, providing a more accurate representation of reliability across meaningful connections in our FC analysis.

#### Assessment of classification performance

We used logistic regression (LR) to assess classification performance of chosen pipelines. LR is a statistical method commonly used for binary classification tasks, where the goal is to predict one of two possible outcomes. LR estimates the probability that a given input belongs to a particular category by fitting data to a logistic function, producing outputs between 0 and 1.

Unlike linear regression, which predicts continuous values, logistic regression uses the logistic (or sigmoid) function to map any real-valued number into a probability. This model is widely used in fields such as medicine and social sciences for problems like disease prediction and risk assessment, because it is powerful yet quite simple and explainable, which is important in high-risk fields.

Because FC matrices are symmetric, we vectorized each matrix by extracting the upper triangular part while excluding the diagonal, and used the resulting feature vector as input to the LR. To mitigate the high dimensionality and multicollinearity of FC features, we applied principal component analysis (PCA) as a preprocessing step before logistic regression. Importantly, PCA was fit only on the training data within each split to avoid information leakage; the learned projection was then applied to transform the held-out data. The number of principal components was selected to explain 95% of the variance in the training set, and logistic regression was trained on the resulting low-dimensional representations. Furthermore, we used logistic regression with L2 regularization (inverse regularization strength C = 1.0, it penalizes large coefficient magnitudes, helping to prevent overfitting) and the LBFGS solver (ma× 1000 iterations) to assess classification performance of chosen pipelines.

In this study, we suppose that the classification performance of machine learning models trained on a given preprocessing pipeline can serve as an indicator of whether the data have been sufficiently preprocessed to enable the model to capture meaningful data patterns. Classification metrics provide quantitative measures for evaluating ML model performance. Among these, accuracy, which is defined as the proportion of correct predictions out of the total number of predictions, is commonly used, particularly when class distributions are balanced. We also report the area under the receiver operating characteristic curve (ROC-AUC), which summarizes discrimination performance across all possible decision thresholds by plotting the true positive rate against the false positive rate. Unlike accuracy, ROC-AUC is less sensitive to class imbalance and provides a threshold-independent measure of how well the model separates the two classes.

With two relatively small datasets prone to overfitting, our evaluation relied on two ways of ML model validation:

1. In the direct cross-site validation protocol, we trained the model using data from one site (source domain; e.g., the full China dataset) and evaluated it on data from a different, entirely unseen site (target domain; e.g., the full IHB dataset). No target-site participants were used at any stage of model development, including preprocessing decisions tied to training, feature selection, hyperparameter tuning, or threshold calibration. This setting provides a stringent test of out-of-site generalization and quantifies the impact of site-specific distribution shifts.
2. In the few-shot domain adaptation protocol, we assessed whether limited access to target-site data could improve transfer to the target domain. Specifically, we augmented the source-site training set (China) with a small, controlled number of target-site participants (e.g., n=20 from IHB) and trained the model on this combined dataset. Performance was then evaluated on the remaining, non-overlapping target-site participants, which were held out from all training and tuning steps. This setting simulates realistic scenarios in which a small calibration sample from a new site is available and tests how efficiently models adapt under constrained target-site labeling.

To avoid information leakage during model training, we performed subject-level splitting such that all scans from the same participant were always assigned to train or to test. Specifically, each subject’s EO and EC runs were kept together in either the training set or the test set, and a participant never contributed one scan to training and another to testing. This design prevents identity leakage, in which a model can learn subject-specific traits rather than patterns genuinely related to the experimental condition. Random seeds were fixed and kept identical across all atlases and preprocessing pipelines to ensure a consistent, directly comparable evaluation setup. To prevent data leakage caused by FC computing, the connectivity-measure estimator was fitted using only the training data in a given split, and the fitted estimator was then applied to the held-out test data.

#### Analysis of stable edges in EO/EC prediction

To identify robust functional-connectivity biomarkers discriminating EO from EC, we quantified the stability of logistic-regression coefficients across repeated random subsamples of a pooled multi-site dataset (IHB + China; N=132 total, 84 IHB and 48 China). The analysis was designed to reduce dependence on any single split and to prioritize edges that were consistently informative across many perturbations. We used the Schaefer200 atlas with Yeo 7-network labels to support network-level interpretability of the resulting stable edges. All preprocessing and nuisance regression used the top-performing denoising strategy from our broader model selection (aCompCor(50%) + 12 HMP).

We performed 1000 iterations of stratified subsampling, selecting 80% of subjects without replacement per iteration while maintaining the IHB/China site proportion (site-stratified subsampling). When a subject was selected into a subsample, both of that subject’s EO and EC scans were included in the training set, preserving the within-subject pairing structure of the EO/EC contrast. In this analysis we used precomputed FC matrices that were estimated once on the full pooled dataset (IHB + China). This choice was made for computational feasibility, because these analyses involve extensive repeated subsampling and recomputing FC (especially the tangent space FC) within each repetition would be prohibitively expensive. Importantly, these analyses did not involve an explicit train/test generalization evaluation.

For each iteration, we trained a logistic regression classifier on PCA-reduced edge features, retaining enough principal components to explain 95% of the variance. Further, to ensure neurophysiologically valid interpretation of the classification features, we applied the Haufe transformation (Haufe et al., 2014) to convert classifier extraction filters (backward model weights) into forward model activation patterns. While backward model weights are optimized to maximize class separability and may not reflect underlying neural sources, forward model activation patterns *α* = Σ*_x_w* (where Σ*x* — data covariance matrix, *w* — weight vector) indicate which functional connections genuinely covary with the EO/EC state. Since the classifier operates in PCA-reduced space, the activation pattern was computed efficiently via the spectral decomposition: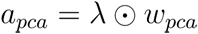, where λ — the PCA eigenvalues, *w_pca_* — the classifier coefficients.

We defined an edge as “stable” if it satisfied two criteria across the 1000 subsamples: 1) sign consistency of at least 80% (the edge’s coefficient direction was preserved in ≥80% of iterations), 2) high importance, operationalized as being ranked within the top 500 edges by absolute coefficient magnitude (approximately the top ∼3% of all connections). This yields a conservative set of biomarkers that are both directionally reliable and repeat among the most discriminative features.

## Results

### QC-FC correlations

In the present study, we calculated QC-FC as the Pearson correlation (absolute value) between subjects’ edge weights (edge in the connectivity matrix) and mean relative RMS displacement, as it was described by (Ciric et al., 2017; Peterson et al., 2025). Then we took the mean edge value across subjects to get the overall view on the influence of motion artifacts. The lower the QC-FC score, the less residual movement is left in the data. The results of the comparison between the processing strategies analyzed with rs-fMRI data and the FC calculation strategies including both GSR and its absence are presented in Figure 2 for the EC state.

**Figure 2.**
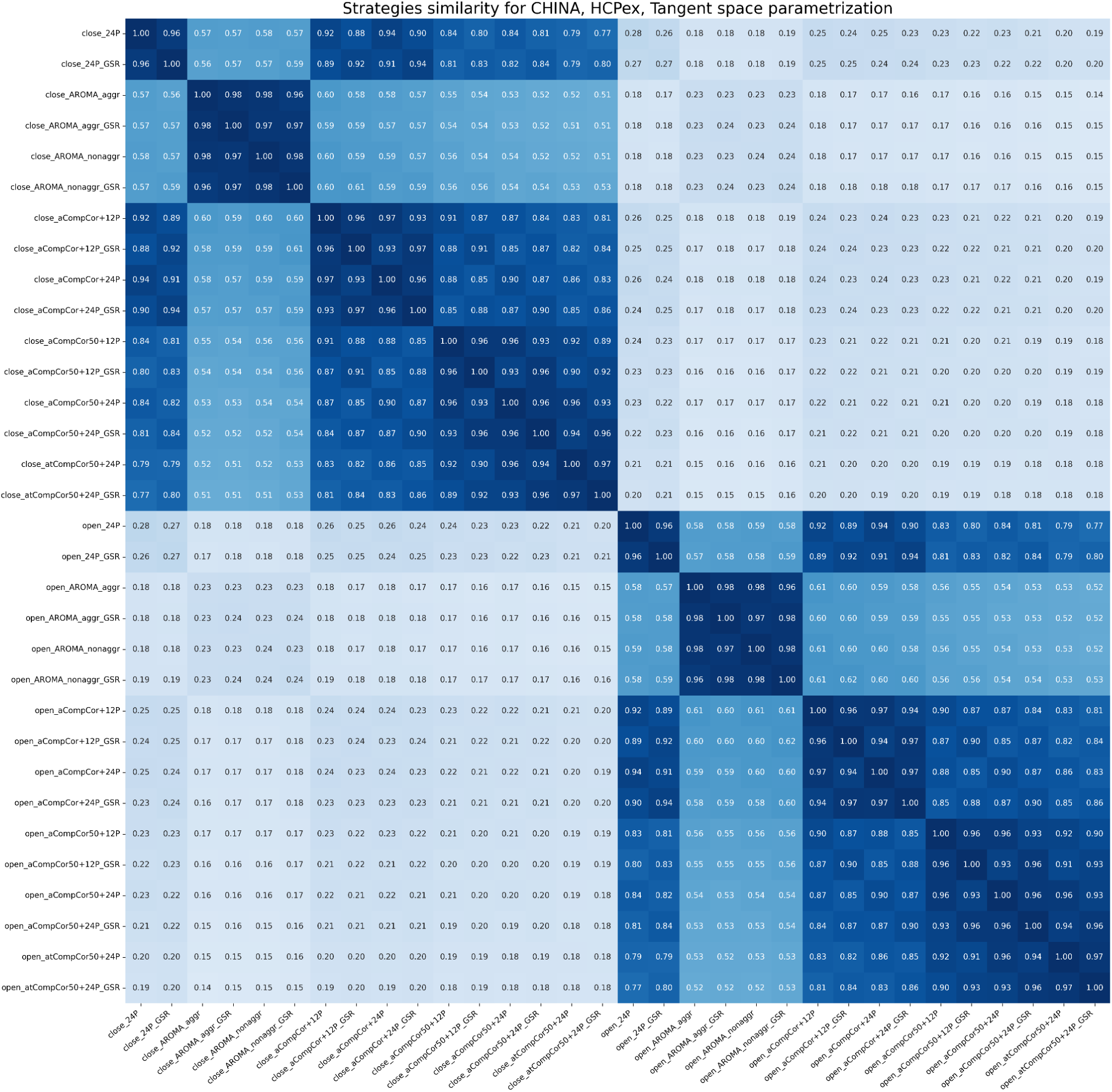
Similarity between signals of both EO and EC states denoised using different strategies for the China dataset, the HCPex atlas, tangent space FC. The heatmap shows similarity for one FC measure. The color range is from white (0, low similarity) to dark blue (1, high similarity). Mean values across subjects are presented.

For QC-FC, the ANOVA factor importance model indicated substantial effects of FC type (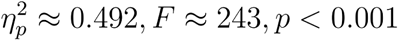) and site (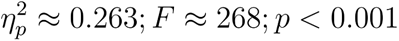). Surprisingly, smaller but statistically reliable effects were observed for denoising strategy (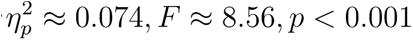) and eyes state (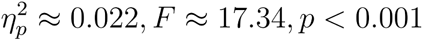). GSR showed a modest but significant association with QC-FC (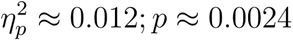), while atlas choice contributed negligibly (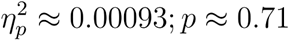). The lowest QC-FC scores were dominated by precision-regularized estimators (e.g., graphical lasso and partial correlation) and CompCor-based denoising without GSR. Top-5 results according to QC-FC score are shown in Table 2 (ranking was based on lowest absolute correlation with motion, top-20 for both states can be found in Supplementary Materials).

**Table 2.**
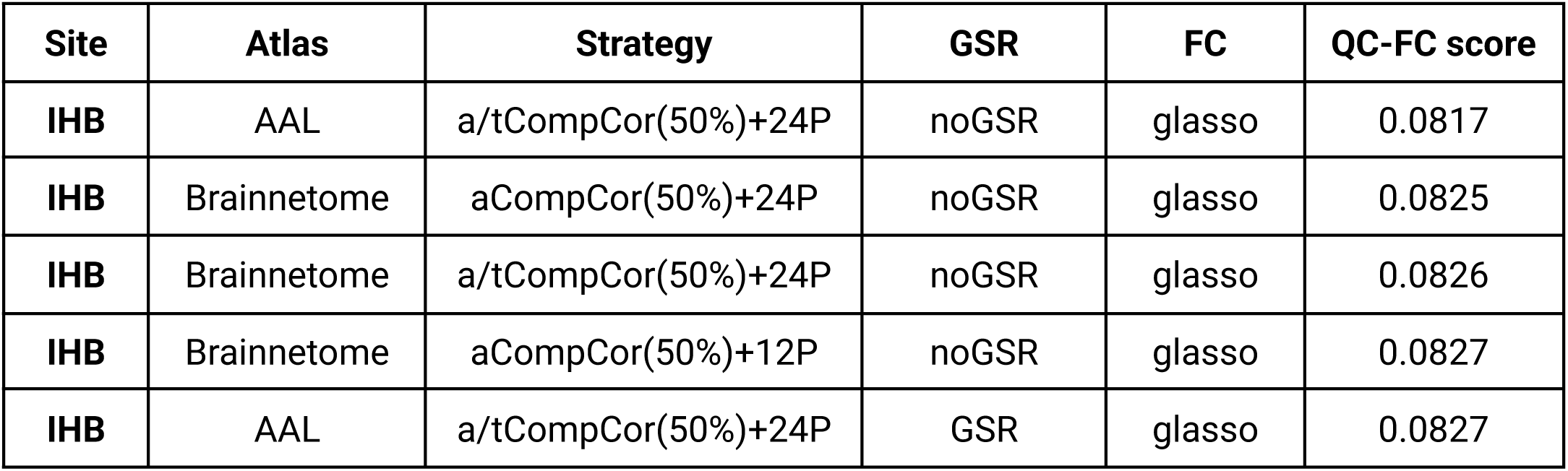
Top-5 pipelines for EC according to QC-FC score.

### Similarity between denoising strategies

As mentioned earlier, the similarity between denoising strategies was measured as the correlation between FC vectors of different denoising strategies and brain states. As a result, we got a matrix, where each entry is a mean similarity value across subjects. In Figures 3 and 4, an example for the China dataset is presented for the HCPex atlas, for partial correlation and tangent space FC. Other pipelines can be found in Supplementary Materials.

**Figure 3.**
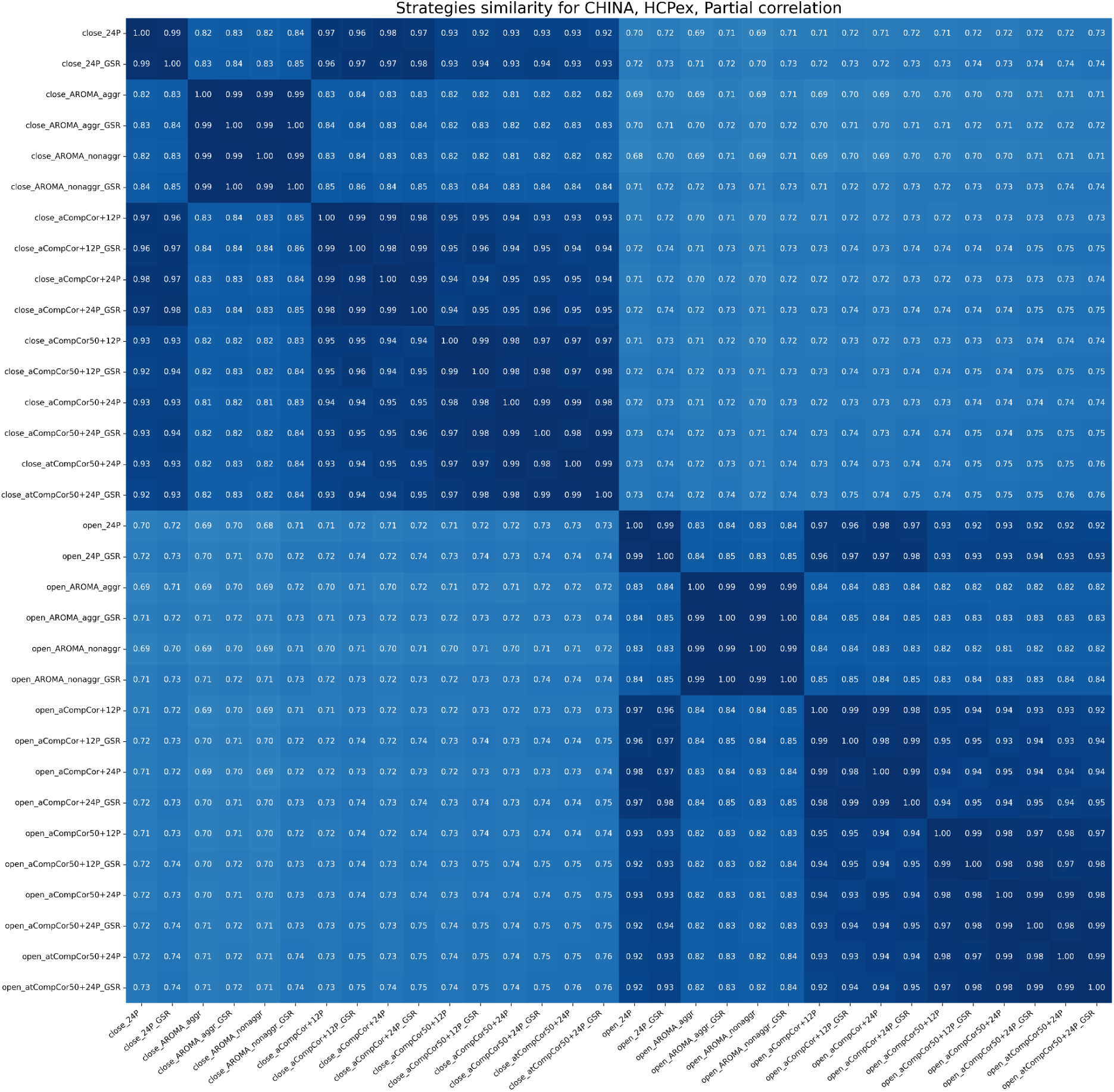
Similarity between signals of both EO and EC states denoised using different strategies for the China dataset, the HCPex atlas, partial correlation FC. The heatmap shows similarity for one FC measure. The color range is from white (0, low similarity) to dark blue (1, high similarity). Mean values across subjects are presented.

**Figure 4.**
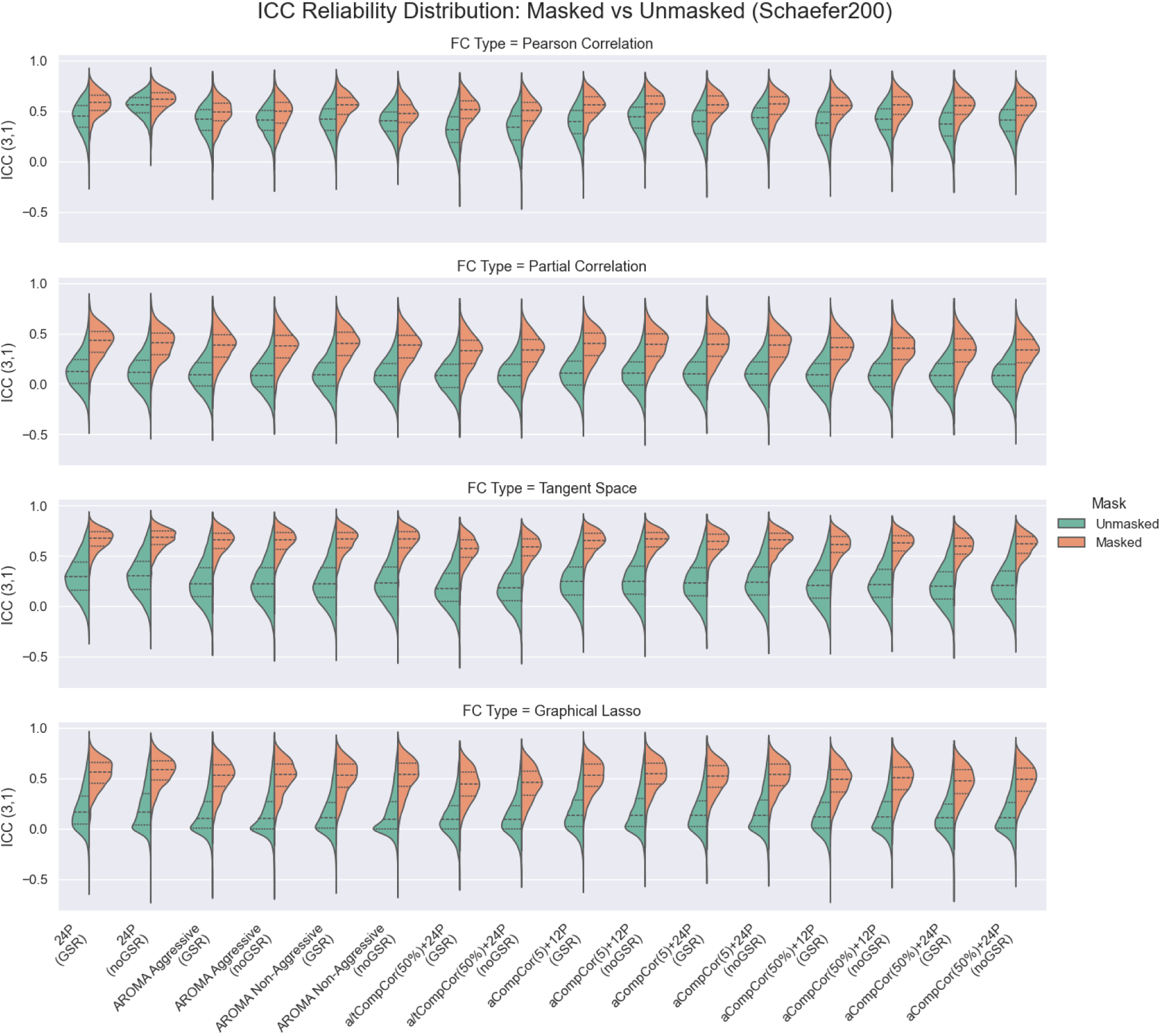
Comparison of ICC scores with (orange) and without masking (green) of near-zero FC weights. Schaefer200 atlas taken for visualization. Every violin plot shows the distribution of ICC scores for every FC measure (rows) and strategies (columns). The central line within each violin plot represents the median. The upper and lower lines indicate the 25th percentile and the 75th percentile. There is no practical need in applying masking onto Pearson correlation, but we give it for comparison.

The results indicate that the magnitude of the EO–EC difference depends strongly on the chosen FC measure. In Figures 3–4, the separation between EO and EC is more pronounced for the tangent representation than for partial correlation, suggesting that these measures capture state-related effects with different sensitivity. The same pattern is reflected in the classification results: when the two classes are more similar in feature space, a linear classifier has less separable structure to exploit, which leads to lower discrimination performance. Moreover, the similarity within “strategy families” is clearly evident: pipelines using ICA-AROMA tend to be more similar to each other than to CompCor-based pipelines, and the same holds in the opposite direction for CompCor strategies.

### Intraclass Correlation Coefficient

Since ICC should be calculated using a dataset that has both multiple scanning sessions and repeated conditions (scanning with EC occurred twice), we calculated it only for the China dataset. The China data had 3 sequentially recorded sessions: first session was only with closed eyes, second and third were alternately closed or open. Therefore, we calculated ICC between first session EC and second session EC, first session EC and third session EC.

Given that (Peterson et al., 2025) hypothesized that the average ICC scores for sparse FC measures tend to zero, because most values of such FC matrices are close to zero and vary from subject to subject only due to noise, we checked if there were differences in ICC scores for pipelines with masking (thresholding) of near-zero entries and without it. The comparison of ICC scores between the masked and unmasked FC matrices is shown in Figure 4.

For masked ICC (ICC(3,1), ICC(2,1), ICC(1,1)), N-way ANOVA and OLS models explained most of the observed variability across pipelines (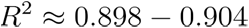 across ICC variants). ANOVA factor importance consistently ranked FC estimator as the dominant driver of ICC (e.g., 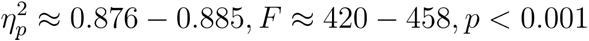), followed by denoising strategy (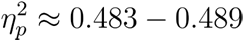) and atlas (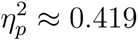), whereas GSR had a negligible and non-significant contribution (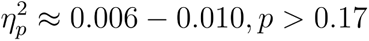).

In Table 3, the top-performing configurations concentrated around the Schaefer200 atlas combined with tangent-space FC (Table 3 is ranked by masked ICC(3,1), top-5 pipelines are listed in Table 3, top-20 list can be found in Supplementary Materials).

**Table 3.**
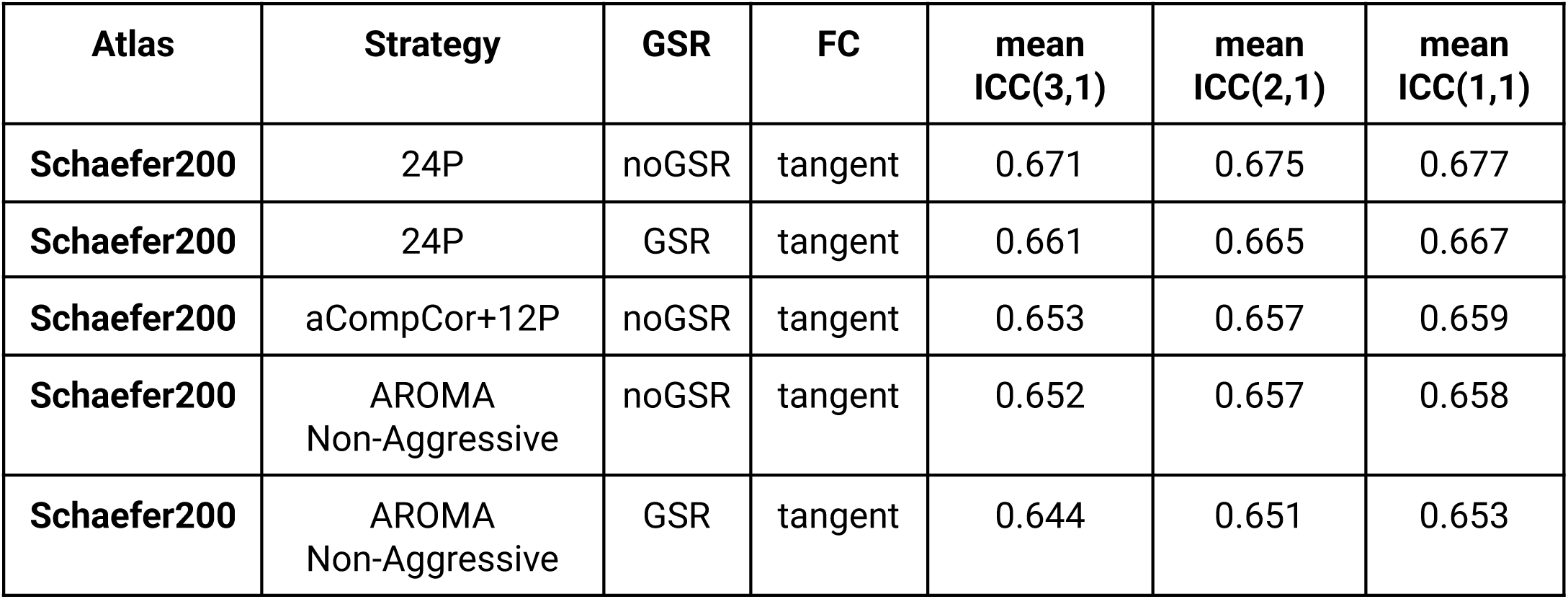
Top-5 pipelines according to masked ICC(3,1) score.

| Atlas | Strategy | GSR | FC | mean ICC(3,1) | mean ICC(2,1) | mean ICC(1,1) |
| --- | --- | --- | --- | --- | --- | --- |
| Schaefer200 | 24P | noGSR | tangent | 0.671 | 0.675 | 0.677 |
| Schaefer200 | 24P | GSR | tangent | 0.661 | 0.665 | 0.667 |
| Schaefer200 | aCompCor+12P | noGSR | tangent | 0.653 | 0.657 | 0.659 |
| Schaefer200 | AROMA<br>Non-Aggressive | noGSR | tangent | 0.652 | 0.657 | 0.658 |
| Schaefer200 | AROMA<br>Non-Aggressive | GSR | tangent | 0.644 | 0.651 | 0.653 |

### Selecting top-performing pipelines for brain state classification

As previously noted, we used a multi-site dataset to train a logistic regression model for a binary classification task – EC versus EO with balanced class distributions using two approaches: few-shot domain adaptation and direct cross-site validation. Classification metrics (accuracy) are presented in Figure 5 for a few-shot strategy – train on China (96 samples) + IHB (40 samples), test on the rest of IHB (128 samples). Top-5 pipelines according to average cross-site accuracy and ROC-AUC are presented in Table 4 and Table 5 respectively (top-20 pipelines according to both ROC-AUC and accuracy can be found in Supplementary Materials).

**Figure 5.**
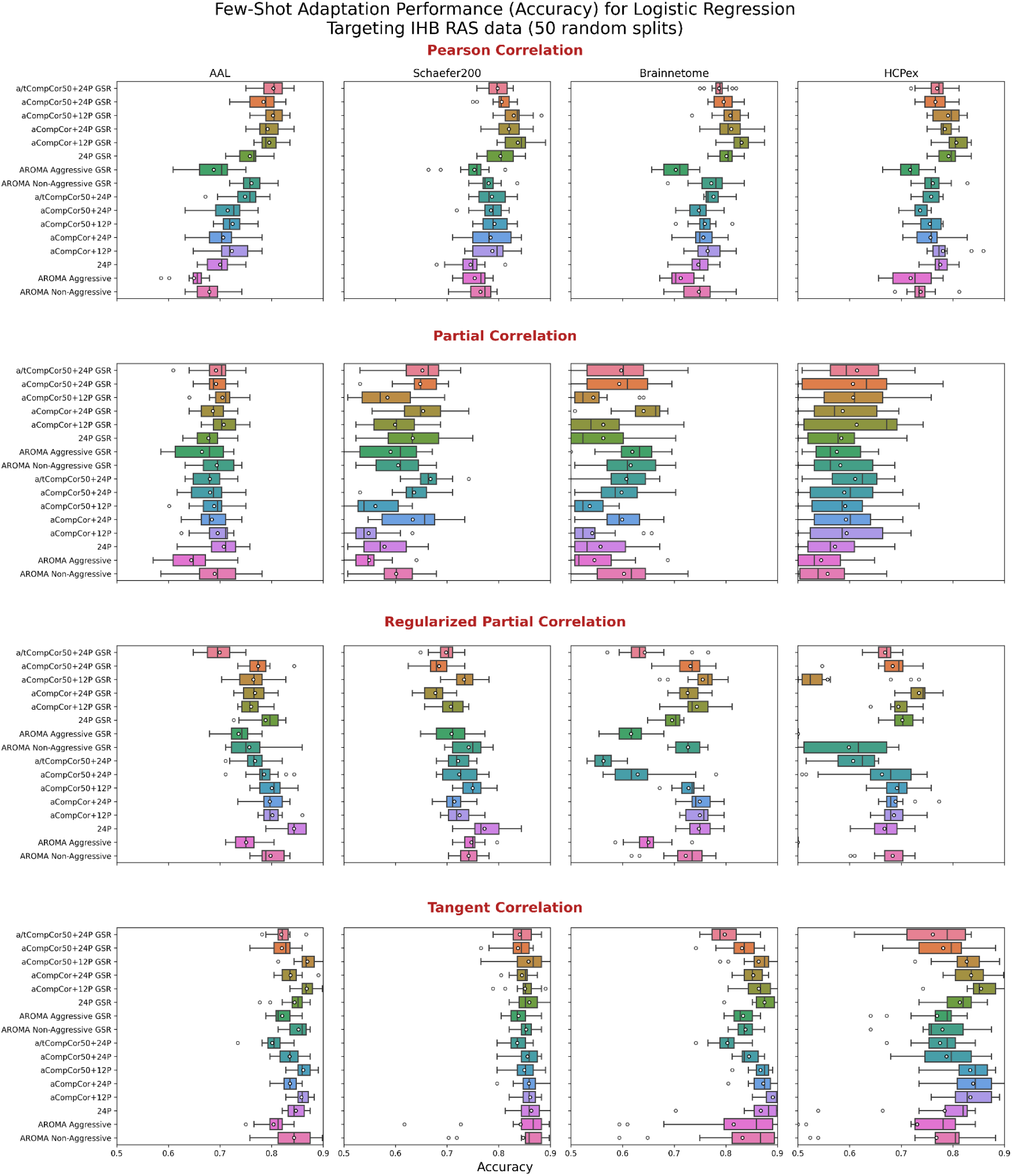
Classification accuracy for logistic regression models using different connectivity estimation methods and parcellations. The models were trained using full China data and a subset of IHB data and tested on the rest of IHB RAS data. Each subplot shows accuracy distributions across 15 iterations for combinations of denoising strategies, parcellation schemes, and connectivity estimation methods. Boxplots illustrate the median, interquartile range, and outliers. Overall, performance varies depending on the choice of preprocessing, parcellation, and connectivity metric.

**Table 4.** Top-5 pipelines according to average cross-site ROC-AUC.

| Atlas | Strategy | GSR | FC | China to IHB ROC-AUC | IHB to China ROC-AUC | Average cross-site ROC-AUC |
| --- | --- | --- | --- | --- | --- | --- |
| **Brainnetome** | 24P | GSR | tangent | 0.908 | 0.936 | 0.922 |
| **Schaefer200** | AROMA Non-Aggressive | GSR | tangent | 0.873 | 0.955 | 0.914 |
| **Schaefer200** | aCompCor+12P | noGSR | tangent | 0.884 | 0.939 | 0.912 |
| **HCPex** | aCompCor+12P | noGSR | tangent | 0.886 | 0.936 | 0.911 |
| **HCPex** | 24P | GSR | tangent | 0.88 | 0.940 | 0.910 |

**Table 5.** Top-5 pipelines according to average cross-site accuracy.

| Atlas | Strategy | GSR | FC | China to IHB accuracy | IHB to China accuracy | Average cross-site accuracy |
| --- | --- | --- | --- | --- | --- | --- |
| **Brainnetome** | aCompCor+12P | noGSR | tangent | 0.809 | 0.864 | 0.837 |
| **Brainnetome** | aCompCor+24P | noGSR | tangent | 0.762 | 0.875 | 0.818 |
| **AAL** | aCompCo+12P | noGSR | tangent | 0.738 | 0.895 | 0.816 |
| **AAL** | aCompCor(50%)+<br>24P | noGSR | tangent | 0.714 | 0.906 | 0.810 |
| **AAL** | aCompCor+24P | noGSR | tangent | 0.702 | 0.906 | 0.804 |

To evaluate the relative importance of pipeline factors on classification performance, we employed a linear mixed-effects model (LMM) with the Brier score as the dependent variable, pipeline factors (FC type, atlas, denoising strategy, and GSR) as fixed effects, and a test subject as a random intercept, thereby accounting for the repeated-measures structure arising from the same subjects being evaluated under all pipeline configurations. The Brier score (the squared difference between predicted probability and true label) was chosen because it is a strictly proper scoring rule that decomposes additively across samples, enabling per-subject aggregation required for the mixed model. In contrast, classification accuracy at the sample level is binary and unsuitable for linear modeling, while ROC-AUC is a population-level rank statistic that cannot be decomposed per subject. Across all pipeline configurations, the Brier score was strongly anti-correlated with both accuracy (Pearson r = −0.91) and AUC (r = −0.82), confirming that factor rankings derived from Brier loss are consistent with those based on accuracy or ROC-AUC.

The results are presented in Table 6. Factor importance was assessed via Type III Wald tests of fixed effects. The LMM confirmed that FC type is by far the most important factor across both directions, followed by Atlas and Strategy.

**Table 6.** The LMM results on factor importance.

|  | **China → IHB** |  | **IHB → China** |  | **Combined** |  |
| --- | --- | --- | --- | --- | --- | --- |
|  | *Wald $\chi^2$* | *p-value* | *Wald $\chi^2$* | *p-value* | *Wald $\chi^2$* | *p-value* |
| **FC** | 1085 | $< 10^{-15}$ | 991.17 | $< 10^{-15}$ | 1508.13 | $< 10^{-15}$ |
| **Atlas** | 270.33 | $< 10^{-15}$ | 8.39 | 0.039 | 231.59 | $< 10^{-15}$ |
| **Strategy** | 56.06 | $< 10^{-10}$ | 167.02 | $< 10^{-15}$ | 119.78 | $< 10^{-15}$ |
| **GSR** | 0.07 | 0.794 | 22.06 | $2.6 \times 10^{-6}$ | 4.40 | 0.036 |

For pairwise factor-level comparisons (e.g., GSR vs. noGSR, tangent vs. Pearson correlation), we employed nonparametric sign-flip randomization tests that inherently account for the paired/repeated structure. These tests:

1. Identify matched pipeline pairs that differ only on the factor of interest,
2. Compute per-sample losses for each pipeline in the pair,
3. Aggregate losses to the subject level (averaging over all matched pairs per subject),
4. Perform a sign-flip test on the subject-level mean differences (5000 permutations).

This approach makes no distributional assumptions and directly respects the within-subject structure by treating the subject as the unit of randomization. P-values were corrected for multiple comparisons using the Benjamini-Hochberg FDR method within each factor.

A total of 82 pairwise comparisons were tested (all factor-level pairs × 2 cross-site directions), of which 48 (58.5%) were significant after FDR correction at α = 0.05. The key results are summarized below.

#### FC type (10/12 significant)

Tangent space was significantly better than all other FC types in both directions (i.e. China→IHB and IHB→China). The largest effects were tangent space vs. partial correlation (Δ = −0.131, p_FDR < 0.001 in China→IHB; Δ = −0.111, p_FDR < 0.001 in IHB→China). Pearson correlation was significantly better than partial correlation in both directions. The two non-significant comparisons were Pearson correlation vs. tangent space in IHB→China (Δ = −0.022, p_FDR = 0.073) and glasso vs. partial correlation in IHB→China (Δ = +0.009, p_FDR = 0.199).

#### Atlas (5/12 significant)

Atlas effects were significant only in the China→IHB direction, where AAL was the best-performing atlas (significantly better than all others) and HCPex was the worst (significantly worse than Brainnetome and Schaefer200). No atlas comparisons reached significance in IHB→China, suggesting atlas choice is less critical when generalizing to a larger test set.

#### Denoising strategy (32/56 significant)

ICA-AROMA aggressive was consistently the worst strategy, significantly worse than all standard CompCor-based strategies in both directions (all p_FDR < 0.05). Among CompCor strategies, strategy 4 (aCompCor+24P) was generally among the best. The non-aggressive AROMA variant performed better than aggressive but still significantly worse than most standard strategies.

#### GSR (1/2 significant)

GSR was significantly better than noGSR only in the IHB→China direction (Δ = +0.015, p_FDR < 0.001). In the China→IHB direction, the difference was negligible and non-significant (Δ = −0.001, p_FDR = 0.608), consistent with the LMM finding that GSR is the least important factor overall.

To sum up, FC type was by far the strongest determinant of performance, followed by atlas and denoising strategy, while GSR had only a minor and direction-dependent effect. In particular, tangent-space connectivity consistently yielded the lowest Brier loss, whereas ICA-AROMA aggressive denoising was reliably associated with worse performance than CompCor-based strategies.

#### Analysis of stable edges in EO/EC prediction

Across repeated subsampling, both Pearson correlation and tangent space projection FC yielded consistent, interpretable network-level signatures distinguishing EO from EC. Pearson correlation produced 447 stable edges (214 EO, 233 EC), whereas tangent-space connectivity identified 299 stable edges (153 EO, 146 EC).

At the large-scale network level, the Haufe transformation patterns revealed coherent and interpretable organization across both FC representations. For the EO state, the most prominent increases in connectivity were observed between the Control and Default Mode networks (40 stable edges in Pearson correlation FC, 22 in tangent space FC), as well as between Control–Somatomotor and Default–Somatomotor systems. This pattern suggests that EO is characterized by strengthened coupling between frontoparietal control regions, default-mode hubs, and sensorimotor areas.

In contrast, the EC state was associated with heightened coupling between Visual and Somatomotor networks, as well as between Visual and Dorsal Attention networks (23 and 21 stable correlation edges for SomMot–Vis and DorsAttn–Vis, respectively, and 12-13 corresponding edges in tangent FC). Additional EC-dominant increases were detected between Default and Dorsal Attention networks and within the Dorsal Attention network itself. Together, these findings indicate that withdrawal of visual input is accompanied by tighter synchronization between visual cortices, attention systems, and somatomotor regions.

Across both correlation and tangent FC, the Haufe transformation patterns converged on a split organization of the visual system. General visual cortex connectivity with Somatomotor and Dorsal Attention networks preferentially supported the EC state, whereas specific frontoparietal–default pathways, particularly Control–Default coupling, preferentially supported the EO state. These complementary signatures delineate how EO and EC states reorganize large-scale connectivity between control, default, visual, somatomotor, and attention networks, in agreement with prior EO/EC literature (Han et al., 2023; Patriat et al., 2013; Wei et al., 2018; Yan et al., 2009).

## Discussions

In the present study, we systematically benchmarked 256 different rs-fMRI processing pipelines on an eyes-open versus eyes-closed classification task. Our primary finding is the remarkable consistency of high classification accuracy across a wide array of pipeline configurations and between two independently acquired datasets. This suggests that to differentiate physiologically distinct human brain states associated with eye condition, the core information embedded in the rs-fMRI FC patterns is largely robust to considerable methodological variability.

The choice of an EO/EC paradigm was strategic, as it represents a well-defined physiological manipulation known to induce large-scale changes in brain activity and functional connectivity (Agcaoglu et al., 2019; Han et al., 2023; Weng et al., 2020; Yan et al., 2009). The pronounced nature of these state differences likely contributes to the observed robustness, as the strong signal-to-noise ratio of the underlying biological effect may overshadow more subtle variations introduced by different processing approaches. This contrasts with studies involving more nuanced cognitive tasks or heterogeneous clinical populations, where pipeline choices might exert a more critical influence on outcomes.

Despite the overall high accuracy, our results revealed performance differences attributable to specific methodological choices. Among the compared functional connectivity measures, the most superior and stable classification performance was revealed for the tangent space parametrization and Pearson correlation. Tangent space parametrization, in particular, demonstrated high accuracy, aligning with previous reports of its effectiveness in brain-state prediction (Dadi et al., 2019; Ng et al., 2017) and fingerprinting (Abbas et al., 2023). However, this method is reported less frequently than the more popular partial correlation or Pearson correlations (about 9000 papers in the last 5 years that mention Pearson correlation as their method versus about 1100 mentioning tangent space parametrization, based on a search on Semantic Scholar).

In the present study, partial correlation and regularized partial correlation consistently yielded lower metrics. This poorer performance could be attributed to several factors, including: estimation instability and “curse of dimensionality”, especially with a higher number of ROIs compared to samples; sensitivity to noise, which might be amplified when conditioning out other regions; and the regularization step potentially being too aggressive or difficult to optimize. On the other hand, both partial and regularized partial correlation showed better reliability, compared to Pearson correlation, which does not contradict conclusions by (Peterson et al., 2025). Nevertheless, the computational complexity associated with these partial correlation methods, in addition to their suboptimal performance in this specific classification task, suggests that simpler correlation measures might be more preferable for similar high-contrast state classifications.

The atlas choice showed moderate effect on classification performance. AAL and Brainnetome had both higher classification metrics and lower QC-FC scores, whereas Schaefer200 showed significant impact on reliability and ICC scores. High-resolution HCPex atlas could potentially introduce more noise or computational burden for the logistic regression model, especially with connectivity measures sensitive to dimensionality (i.e. regularized partial correlation). These results align with previous findings, where the authors discussed the impact of brain atlas size on the results of the study (Khan & Shang, 2025; Wu et al., 2025). The authors agreed that more detailed brain atlases provided better connectivity features, but required more computational resources. In this sense, AAL, Schaefer200 and Brainnetome are an optimal solution for this particular task.

Denoising strategies based on the inclusion of GSR had no significant impact on classification accuracy or reliability. Some authors report that GSR has little or no effect on classification performance (Shi et al., 2021; Sotero et al., 2023).

Cross-site validation demonstrated generalizability between the datasets used. Models trained on data from one site successfully generalized to unseen data from the other site. Our stable-edge results are consistent with, and do not contradict, prior EO/EC resting-state findings showing that eye state induces widespread, network-level reconfiguration (Agcaoglu et al., 2019; Costumero et al., 2020; Han et al., 2023; Weng et al., 2020). In particular, the prominent role of sensory and attention systems in our biomarkers (Visual-Somatomotor and Visual-Dorsal Attention edges contributing to EC vs EO discrimination) aligns with earlier reports that EO/EC modulates connectivity across visual and sensorimotor networks and their coupling with other systems. Likewise, the stable involvement of default-mode and control/attention circuitry in our EO-related edges (e.g., Control–Default coupling and within-DMN connectivity) matches the established observation that DMN and attentional/control networks show condition-dependent connectivity differences between EO and EC.

### Limitations

The primary limitation of the present study is the high contrast nature of the EO/EC states. The robustness observed here may not be extended to experimental tasks with subtler neural signatures or studies aiming to detect small effect sizes in clinical cohorts. Secondly, we employed a single classification algorithm (Logistic Regression); while chosen for its simplicity, other more complex models might interact differently with pipeline choices. Furthermore, due to the enormous number of possible pipeline combinations, we tested only a small part of the possible options, not including other popular denoising methods, (e.g. scrubbing) (Ciric et al., 2017), parcellation methods (other brain atlases and data-driven parcellations) (Wu et al., 2025), and FC measures (Z.-Q. Liu et al., 2025). Finally, our study focused on healthy participants and pipeline effects in patient populations could differ substantially from healthy cohort.

### Implications and Future Directions

Our findings bolster confidence in the use of FC-based classification for brain states that are physiologically well distinguished and reproducible. The resilience to pipeline variability and cross-site differences is encouraging for multi-site studies and biomarker development, suggesting that meaningful signals can be extracted even without perfect pipeline harmonization, especially for strong biological effects. Future research should extend this benchmarking approach to more complex cognitive paradigms, clinical data, and investigate the interaction between pipeline choices and more sophisticated machine learning architectures. Further exploration into the asymmetrical cross-site generalization performance could also yield insights into optimizing multi-site study designs.

## Conclusion

This study systematically benchmarked the impact of 256 distinct rs-fMRI processing pipelines on the classification of resting states with open eyes versus closed eyes, utilizing a rigorous cross-site validation strategy. Our central finding is the remarkable robustness of FC-based predictive models: despite substantial variability in BOLD signal denoising strategies, brain parcellation schemes, and connectivity estimation metrics, high classification accuracy was consistently achieved across both acquisition sites. Our results demonstrate successful cross-site generalization, highlighting the potential of functional connectivity measures as robust markers even when data are collected in different settings.

In summary, our study shows that FC-based classification reliably differentiates fundamental brain states in resting conditions with open versus closed eyes in diverse processing pipelines and acquisition settings. Despite the robustness of the results regarding the pipelines used, we found that FC choice is the most important factor determining high metrics and reliability. Moreover, Pearson correlation is deservedly considered the most popular FC measure, but its effectiveness strongly depends on the quality of the data denoising. In this sense, tangent space parametrization can be a good compromise, combining stability to noise and relatively low computational complexity, compared to partial and regularized partial correlation.

## Research funding

This study was performed within the state assignment of the Ministry of Education and Science of Russian Federation (IHB RAS budget project № 125030703331-8)

## Data availability statement

The analysis code and all scripts for reproducing the results, including denoising pipelines, atlas processing, connectivity computation, ICC and QC-FC analysis, and ML benchmarking, are publicly available at https://github.com/IHB-IBR-department/BenchmarkingEOEC. The repository includes detailed installation instructions, core workflows, and Markdown documentation for methods and data formats.

All data necessary to reproduce the results of this study (preprocessed timeseries, coverage masks) are available at: https://disk.yandex.ru/d/fNz33QPwJQj7HQ.

Original raw data for China dataset is available at: https://fcon_1000.projects.nitrc.org/indi/retro/BeijingEOEC.html. Financial support for this dataset was provided by a grant from the National Natural Science Foundation of China: 30770594 and a grant from the National High Technology Program of China (863): 2008AA02Z405.

## Footnotes

1 https://fmripost-aroma.readthedocs.io/latest/installation.html

2 Financial support for the data used in this project was provided by a grant from the National Natural Science Foundation of China: 30770594 and a grant from the National High Technology Program of China (863): 2008AA02Z405

3 https://nilearn.github.io/stable/index.html

