## Supplementary Materials for "Benchmarking resting state fMRI connectivity pipelines for classification: Robust accuracy despite processing variability in cross-site eye state prediction"

Supplementary materials present extended results including full-sized tables and figures for all datasets analyzed across sites, denoising strategies, atlases, and connectivity metrics.

### QC-FC Results

Top-20 results according to QC-FC score are shown in Table S1 for EO and Table S2 for EC (ranking was based on lowest absolute correlation with motion).

Table S1. Top-20 pipelines for EO according to QC-FC score

| Site | Atlas | Strategy | GSR | FC | QC_FC |
|---|---|---|---|---|---|
| ihb | Schaefer200 | a/tCompCor(50%)+24P | noGSR | glasso | 0.0784 |
| ihb | AAL | a/tCompCor(50%)+24P | noGSR | glasso | 0.0786 |
| ihb | Brainnetome | a/tCompCor(50%)+24P | noGSR | glasso | 0.0793 |
| ihb | Schaefer200 | aCompCor(50%)+12P | noGSR | glasso | 0.0795 |
| ihb | Schaefer200 | aCompCor(50%)+24P | noGSR | glasso | 0.0795 |
| ihb | Brainnetome | aCompCor(50%)+12P | noGSR | glasso | 0.0798 |
| ihb | Schaefer200 | aCompCor+12P | noGSR | glasso | 0.0803 |
| ihb | Schaefer200 | aCompCor+24P | noGSR | glasso | 0.0804 |
| ihb | Schaefer200 | AROMA Non-Aggressive | noGSR | glasso | 0.0804 |
| ihb | Brainnetome | aCompCor+12P | noGSR | glasso | 0.0806 |
| ihb | Brainnetome | aCompCor(50%)+24P | noGSR | glasso | 0.0807 |
| ihb | Brainnetome | 24P | noGSR | glasso | 0.0807 |
| ihb | Schaefer200 | 24P | noGSR | glasso | 0.081 |
| ihb | Brainnetome | AROMA Non-Aggressive | noGSR | glasso | 0.0812 |
| ihb | Brainnetome | aCompCor+24P | noGSR | glasso | 0.0814 |
| ihb | Brainnetome | aCompCor(50%)+12P | GSR | glasso | 0.0816 |
| ihb | AAL | aCompCor(50%)+24P | noGSR | glasso | 0.0819 |
| ihb | Schaefer200 | aCompCor(50%)+12P | GSR | glasso | 0.082 |
| ihb | Schaefer200 | AROMA Non-Aggressive | GSR | glasso | 0.0821 |
| ihb | Brainnetome | 24P | GSR | glasso | 0.0822 |

Table S2. Top-20 pipelines for EC according to QC-FC score

| Site | Atlas | Strategy | GSR | FC | QC_FC |
|---|---|---|---|---|---|
| ihb | AAL | a/tCompCor(50%)+24P | noGSR | glasso | 0.0817 |
| ihb | Brainnetome | aCompCor(50%)+24P | noGSR | glasso | 0.0825 |
| ihb | Brainnetome | a/tCompCor(50%)+24P | noGSR | glasso | 0.0826 |
| ihb | Brainnetome | aCompCor(50%)+12P | noGSR | glasso | 0.0827 |
| ihb | AAL | a/tCompCor(50%)+24P | GSR | glasso | 0.0827 |
| ihb | Brainnetome | AROMA Non-Aggressive | noGSR | glasso | 0.083 |
| ihb | Schaefer200 | a/tCompCor(50%)+24P | noGSR | glasso | 0.0831 |
| ihb | Schaefer200 | AROMA Non-Aggressive | noGSR | glasso | 0.0832 |
| ihb | AAL | a/tCompCor(50%)+24P | noGSR | partial | 0.0833 |
| ihb | Brainnetome | AROMA Non-Aggressive | GSR | glasso | 0.0833 |
| ihb | Brainnetome | aCompCor+24P | noGSR | glasso | 0.0834 |
| ihb | Schaefer200 | AROMA Non-Aggressive | GSR | glasso | 0.0835 |
| ihb | Schaefer200 | a/tCompCor(50%)+24P | GSR | glasso | 0.0837 |
| ihb | Brainnetome | 24P | noGSR | glasso | 0.0837 |
| ihb | Brainnetome | AROMA Aggressive | GSR | glasso | 0.0837 |
| ihb | AAL | aCompCor(50%)+24P | noGSR | glasso | 0.0838 |
| ihb | Brainnetome | aCompCor+12P | noGSR | glasso | 0.0838 |
| ihb | Brainnetome | a/tCompCor(50%)+24P | GSR | glasso | 0.0838 |
| ihb | Schaefer200 | aCompCor(50%)+24P | noGSR | glasso | 0.084 |
| ihb | Brainnetome | aCompCor(50%)+12P | GSR | glasso | 0.084 |

### ICC results

In Table S3 present top-20 pipelines, the table is ranked by masked ICC(3,1) score.

Table S3. Top-20 pipelines according to masked ICC(3,1)

| Atlas | Strategy | GSR | FC | Mean ICC(3,1) | Mean ICC(2,1) | Mean ICC(1,1) |
|---|---|---|---|---|---|---|
| Schaefer200 | 24P | noGSR | tang | 0.6712 | 0.6757 | 0.6769 |
| Schaefer200 | 24P | GSR | tang | 0.6613 | 0.6658 | 0.6672 |
| Schaefer200 | aCompCor+12P | noGSR | tang | 0.6531 | 0.6577 | 0.6591 |
| Schaefer200 | AROMA Non-Aggressive | noGSR | tang | 0.6525 | 0.6571 | 0.6585 |
| Schaefer200 | AROMA Non-Aggressive | GSR | tang | 0.6465 | 0.6511 | 0.6526 |
| Schaefer200 | aCompCor+24P | noGSR | tang | 0.644 | 0.6486 | 0.6501 |
| Schaefer200 | AROMA Aggressive | noGSR | tang | 0.642 | 0.6466 | 0.6481 |
| Schaefer200 | aCompCor+12P | GSR | tang | 0.6408 | 0.6455 | 0.647 |
| Schaefer200 | AROMA Aggressive | GSR | tang | 0.6405 | 0.6451 | 0.6466 |
| Schaefer200 | aCompCor+24P | GSR | tang | 0.6337 | 0.6384 | 0.64 |
| Schaefer200 | aCompCor(50%)+12P | noGSR | tang | 0.6151 | 0.6199 | 0.6216 |
| AAL | 24P | noGSR | tang | 0.6103 | 0.6151 | 0.6169 |
| Brainnetome | 24P | noGSR | tang | 0.6088 | 0.6135 | 0.6153 |
| Schaefer200 | 24P | noGSR | corr | 0.6046 | 0.5919 | 0.5873 |
| Schaefer200 | aCompCor(50%)+24P | noGSR | tang | 0.6038 | 0.6087 | 0.6105 |
| Brainnetome | 24P | noGSR | corr | 0.6005 | 0.5915 | 0.5883 |
| Schaefer200 | aCompCor(50%)+12P | GSR | tang | 0.5994 | 0.6042 | 0.6061 |
| Brainnetome | AROMA Non-Aggressive | noGSR | tang | 0.5983 | 0.603 | 0.6049 |
| Brainnetome | AROMA Non-Aggressive | GSR | tang | 0.5964 | 0.6011 | 0.603 |
| AAL | 24P | GSR | tang | 0.5942 | 0.599 | 0.6009 |

#### Differences between masked and unmasked ICC

The difference in ICC scores between the masked and unmasked FC matrices is shown in Figures S1-S4 for every atlas.

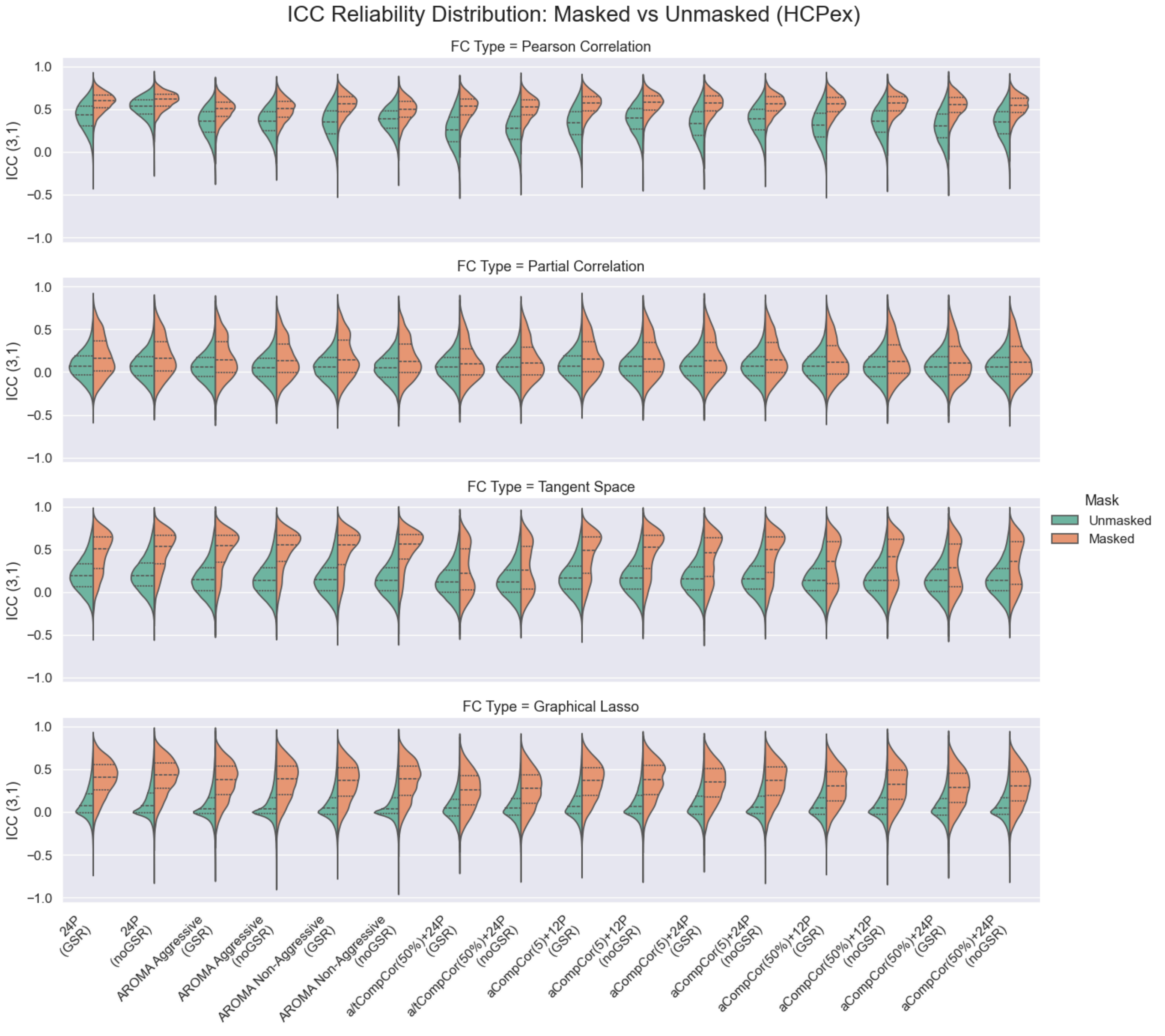


*Figure S1*

*Comparison of ICC scores with (orange) and without masking (green) of near-zero FC weights. HCPex atlas. Every violin plot shows the distribution of ICC scores for every FC measure (rows) and strategies (columns). The central line within each violin plot represents the median. The upper and lower lines indicate the 25th percentile and the 75th percentile. There is no practical need in applying masking onto Pearson correlation, but we give it for comparison.*

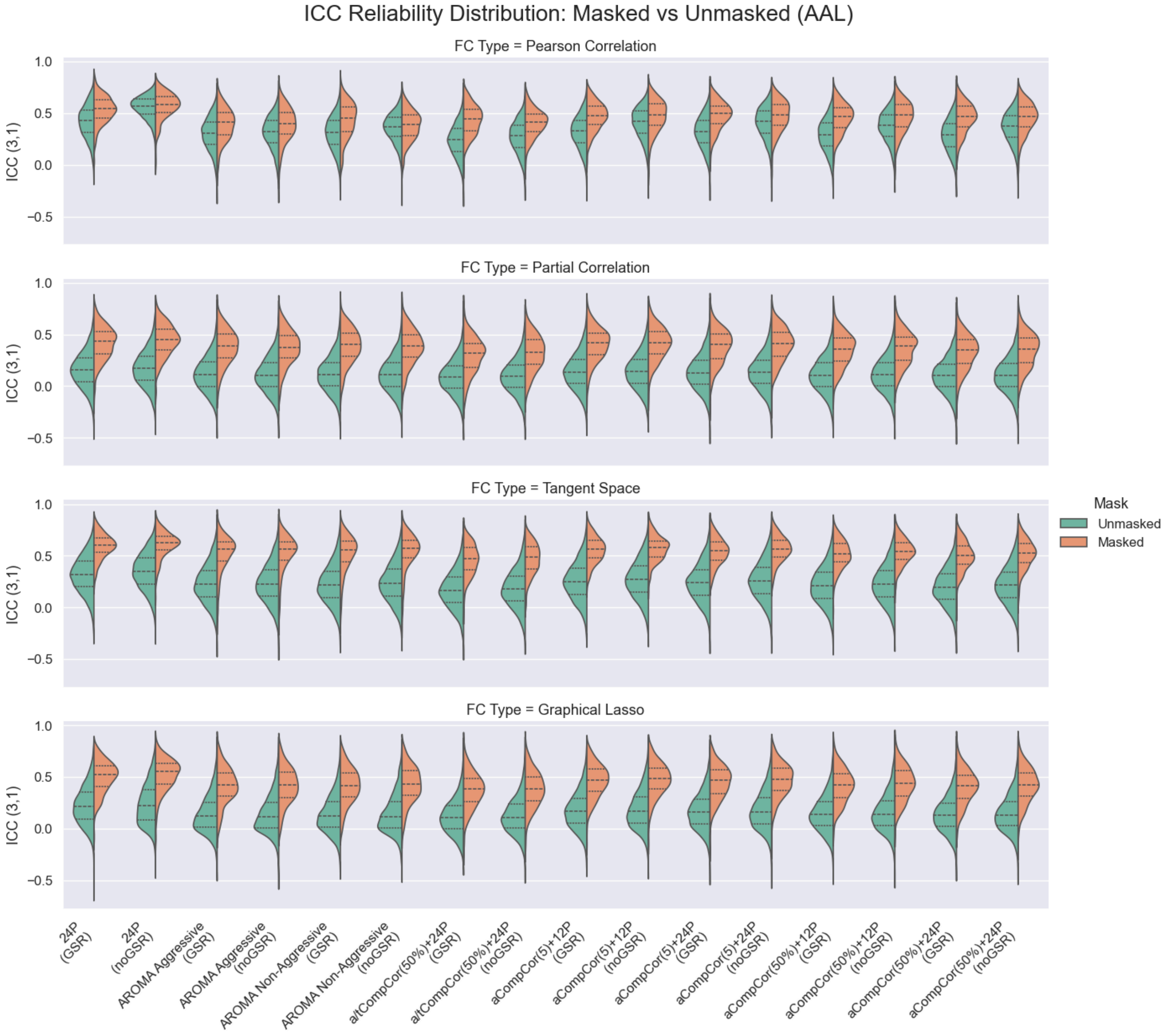


*Figure S2*

*Comparison of ICC scores with (orange) and without masking (green) of near-zero FC weights. AAL atlas. Every violin plot shows the distribution of ICC scores for every FC measure (rows) and strategies (columns). The central line within each violin plot represents the median. The upper and lower lines indicate the 25th percentile and the 75th percentile. There is no practical need in applying masking onto Pearson correlation, but we give it for comparison.*

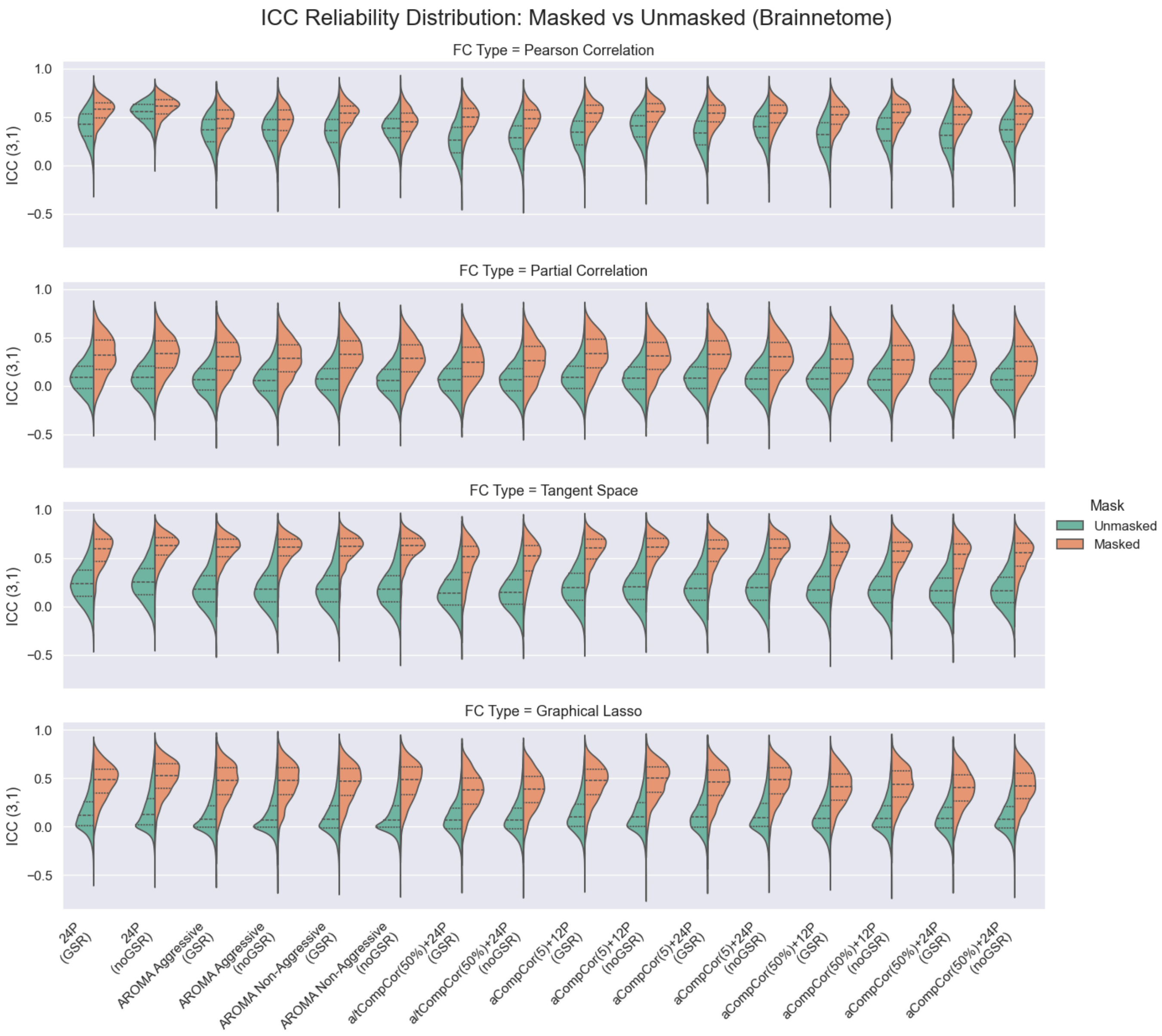


*Figure S3*

*Comparison of ICC scores with (orange) and without masking (green) of near-zero FC weights. Brainnetome atlas. Every violin plot shows the distribution of ICC scores for every FC measure (rows) and strategies (columns). The central line within each violin plot represents the median. The upper and lower lines indicate the 25th percentile and the 75th percentile. There is no practical need in applying masking onto Pearson correlation, but we give it for comparison.*

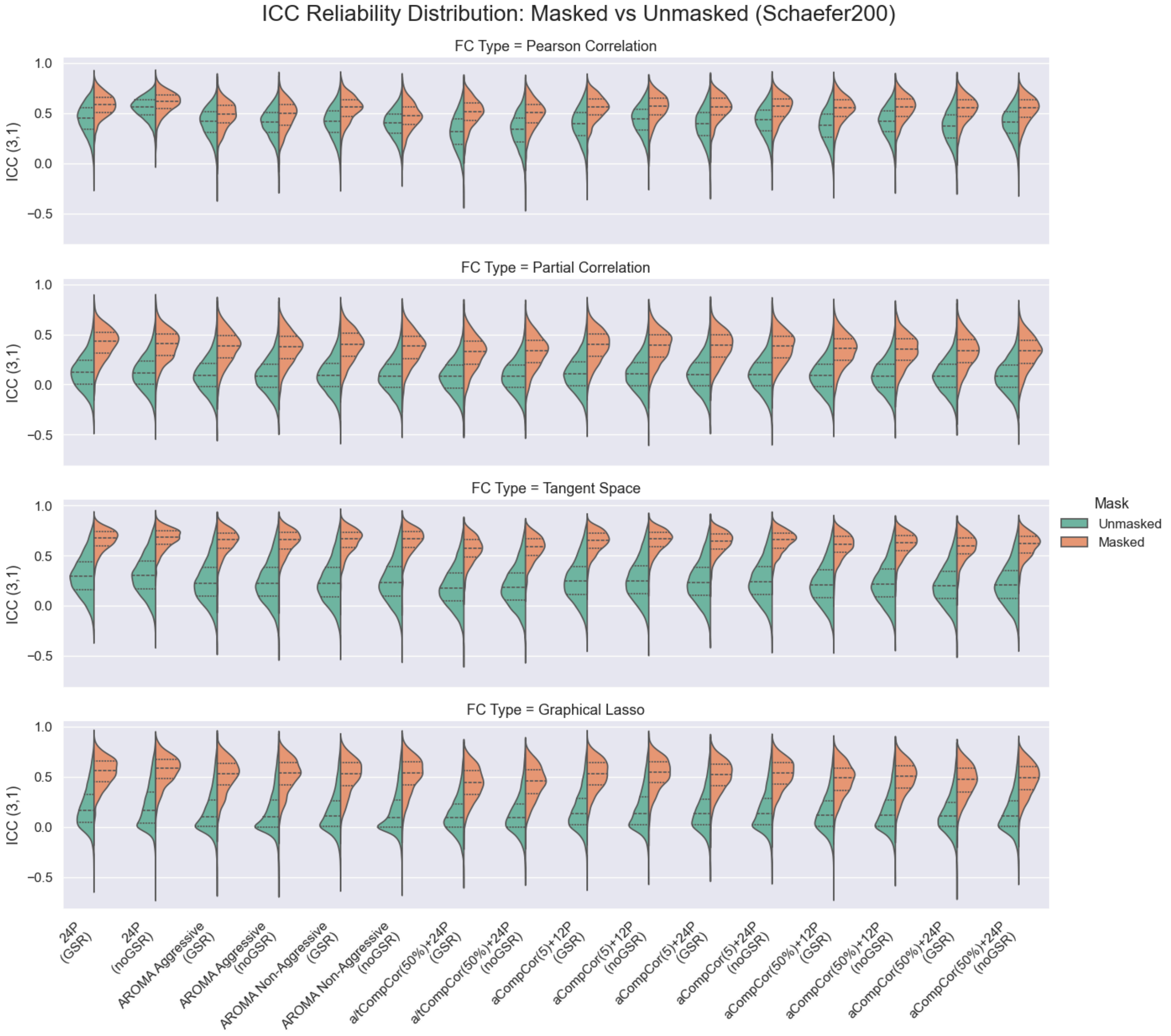


*Figure S4*

*Comparison of ICC scores with (orange) and without masking (green) of near-zero FC weights. Schaefer200 atlas. Every violin plot shows the distribution of ICC scores for every FC measure (rows) and strategies (columns). The central line within each violin plot represents the median. The upper and lower lines indicate the 25th percentile and the 75th percentile. There is no practical need in applying masking onto Pearson correlation, but we give it for comparison.*

### Classification results

Table S4. Top 20 Pipelines (Mean Cross-Site Accuracy)

| Atlas | Strategy | GSR | FC | China to IHB accuracy | IHB to China accuracy | Average cross-site accuracy |
|---|---|---|---|---|---|---|
| Brainnetome | aCompCor+12P | noGSR | tangent | 0.8095 | 0.8646 | 0.8371 |
| Brainnetome | aCompCor+24P | noGSR | tangent | 0.7619 | 0.875 | 0.8185 |
| AAL | aCompCor+12P | noGSR | tangent | 0.7381 | 0.8958 | 0.817 |
| AAL | aCompCor(50%)+24P | noGSR | tangent | 0.7143 | 0.9062 | 0.8103 |
| AAL | aCompCor+24P | noGSR | tangent | 0.7024 | 0.9062 | 0.8043 |
| Brainnetome | AROMA Aggressive | GSR | tangent | 0.7738 | 0.8333 | 0.8036 |
| AAL | aCompCor(50%)+12P | noGSR | tangent | 0.7143 | 0.8854 | 0.7999 |
| Schaefer200 | aCompCor(50%)+12P | noGSR | tangent | 0.7143 | 0.8854 | 0.7999 |
| Schaefer200 | aCompCor+12P | noGSR | tangent | 0.7321 | 0.8646 | 0.7984 |
| Schaefer200 | aCompCor+24P | noGSR | tangent | 0.7202 | 0.875 | 0.7976 |
| Brainnetome | aCompCor+12P | GSR | corr | 0.7083 | 0.8854 | 0.7969 |
| Brainnetome | aCompCor(50%)+24P | noGSR | tangent | 0.7321 | 0.8542 | 0.7932 |
| Schaefer200 | aCompCor(50%)+24P | noGSR | tangent | 0.7262 | 0.8542 | 0.7902 |
| Brainnetome | AROMA Aggressive | noGSR | tangent | 0.744 | 0.8333 | 0.7887 |
| AAL | aCompCor+12P | GSR | tangent | 0.7202 | 0.8542 | 0.7872 |
| AAL | AROMA Non-Aggressive | GSR | tangent | 0.6667 | 0.9062 | 0.7865 |
| AAL | aCompCor(50%)+12P | GSR | tangent | 0.6845 | 0.8854 | 0.785 |
| AAL | aCompCor+24P | GSR | tangent | 0.7024 | 0.8646 | 0.7835 |
| HCPex | aCompCor+12P | noGSR | corr | 0.6905 | 0.875 | 0.7827 |
| HCPex | aCompCor(50%)+12P | noGSR | corr | 0.6786 | 0.8854 | 0.782 |

Table S5. Top 20 Pipelines (Mean Cross-Site ROC-AUC)

| Atlas | Strategy | GSR | FC | China to IHB ROC-AUC | IHB to China ROC-AUC | Average cross-site ROC-AUC |
|---|---|---|---|---|---|---|
| Brainnetome | 24P | GSR | tangent | 0.9082 | 0.9362 | 0.9222 |
| Schaefer200 | AROMA Non-Aggressive | GSR | tangent | 0.8733 | 0.9553 | 0.9143 |
| Schaefer200 | aCompCor+12P | noGSR | tangent | 0.8839 | 0.9392 | 0.9116 |
| HCPex | aCompCor+12P | noGSR | tangent | 0.8865 | 0.9362 | 0.9113 |
| HCPex | 24P | GSR | tangent | 0.8802 | 0.9401 | 0.9102 |
| Brainnetome | 24P | noGSR | tangent | 0.8921 | 0.9266 | 0.9094 |
| HCPex | aCompCor+24P | noGSR | tangent | 0.8831 | 0.9332 | 0.9081 |
| Schaefer200 | aCompCor(50%)+12P | noGSR | tangent | 0.8678 | 0.9457 | 0.9068 |
| Schaefer200 | aCompCor+24P | noGSR | tangent | 0.8764 | 0.9371 | 0.9067 |
| Brainnetome | AROMA Non-Aggressive | GSR | tangent | 0.87 | 0.9423 | 0.9062 |
| Brainnetome | aCompCor+12P | noGSR | tangent | 0.8829 | 0.9275 | 0.9052 |
| Brainnetome | aCompCor+12P | GSR | tangent | 0.8754 | 0.9349 | 0.9052 |
| Schaefer200 | AROMA Non-Aggressive | noGSR | tangent | 0.87 | 0.9401 | 0.9051 |
| Schaefer200 | aCompCor(50%)+12P | GSR | tangent | 0.873 | 0.9366 | 0.9048 |
| Schaefer200 | aCompCor(50%)+24P | noGSR | tangent | 0.8669 | 0.9427 | 0.9048 |
| Brainnetome | aCompCor+24P | noGSR | tangent | 0.8729 | 0.9362 | 0.9045 |
| Brainnetome | aCompCor+24P | GSR | tangent | 0.8712 | 0.9375 | 0.9043 |
| HCPex | 24P | noGSR | tangent | 0.8581 | 0.9457 | 0.9019 |
| Brainnetome | aCompCor(50%)+24P | GSR | tangent | 0.8668 | 0.9358 | 0.9013 |

| | | | | | | |
|---|---|---|---|---|---|---|
| HCPex | aCompCor+12P | GSR | tangent | 0.8601 | 0.9405 | 0.9003 |

### Similarity between denoising strategies

#### China dataset

##### AAL

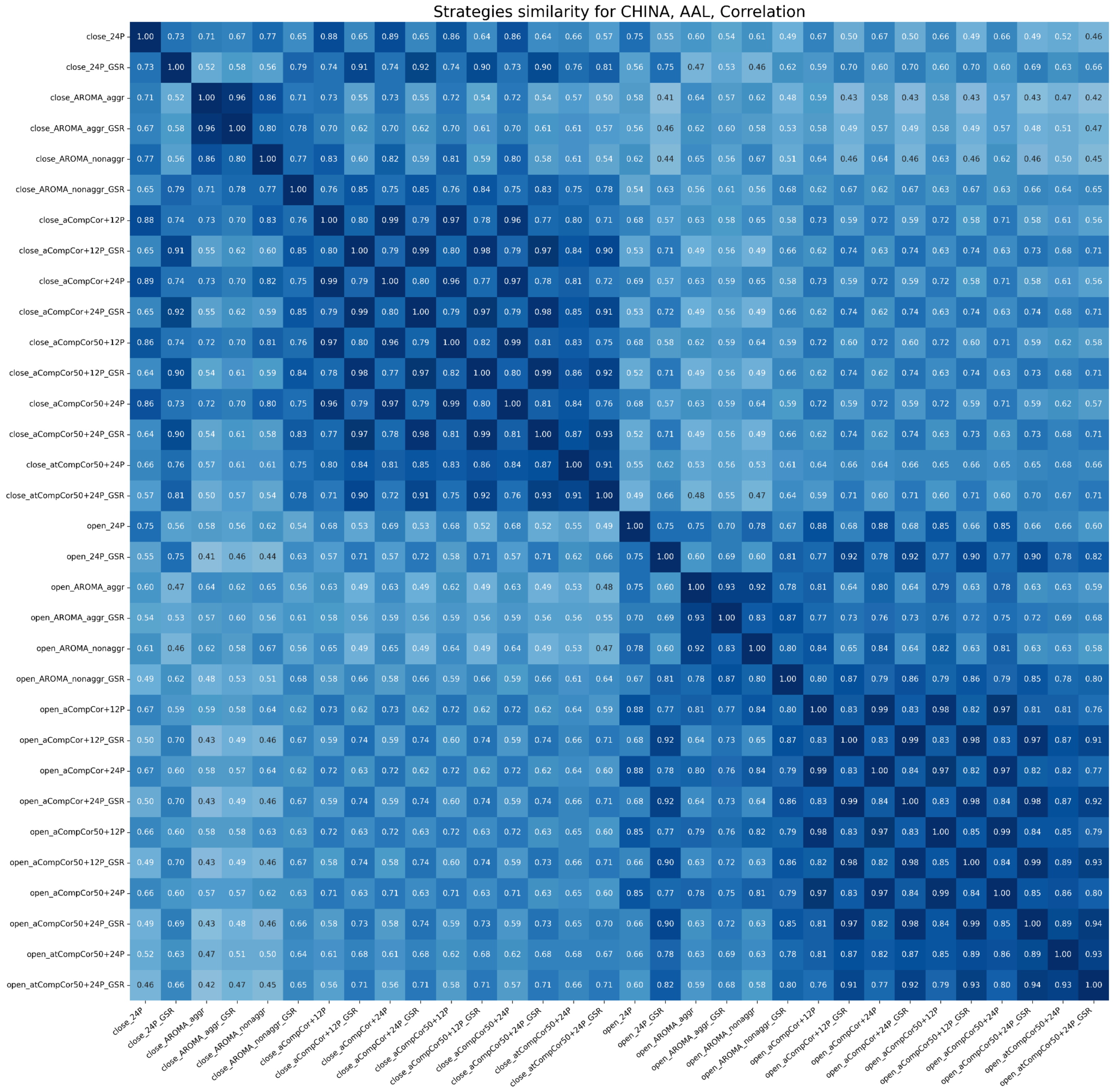


*Figure S5.*

*Similarity between signals of both EO and EC states denoised using different strategies for the China dataset, the AAL atlas, correlation FC. The heatmap shows similarity for one FC measure. The color range is from white (0, low similarity) to dark blue (1, high similarity). Mean values across subjects are presented.*

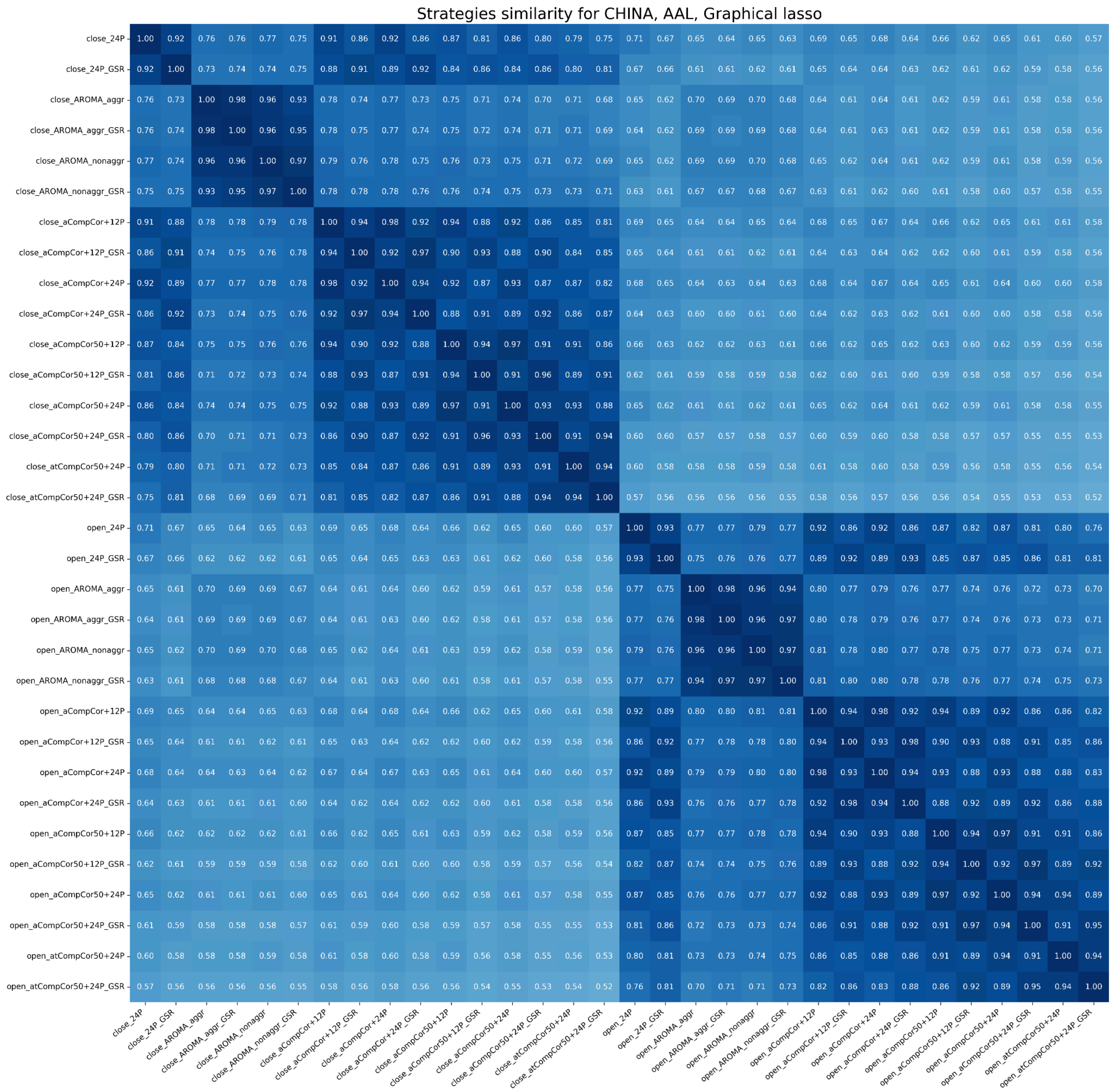


*Figure S6.*

*Similarity between signals of both EO and EC states denoised using different strategies for the China dataset, the AAL atlas, glasso FC. The heatmap shows similarity for one FC measure. The color range is from white (0, low similarity) to dark blue (1, high similarity). Mean values across subjects are presented.*

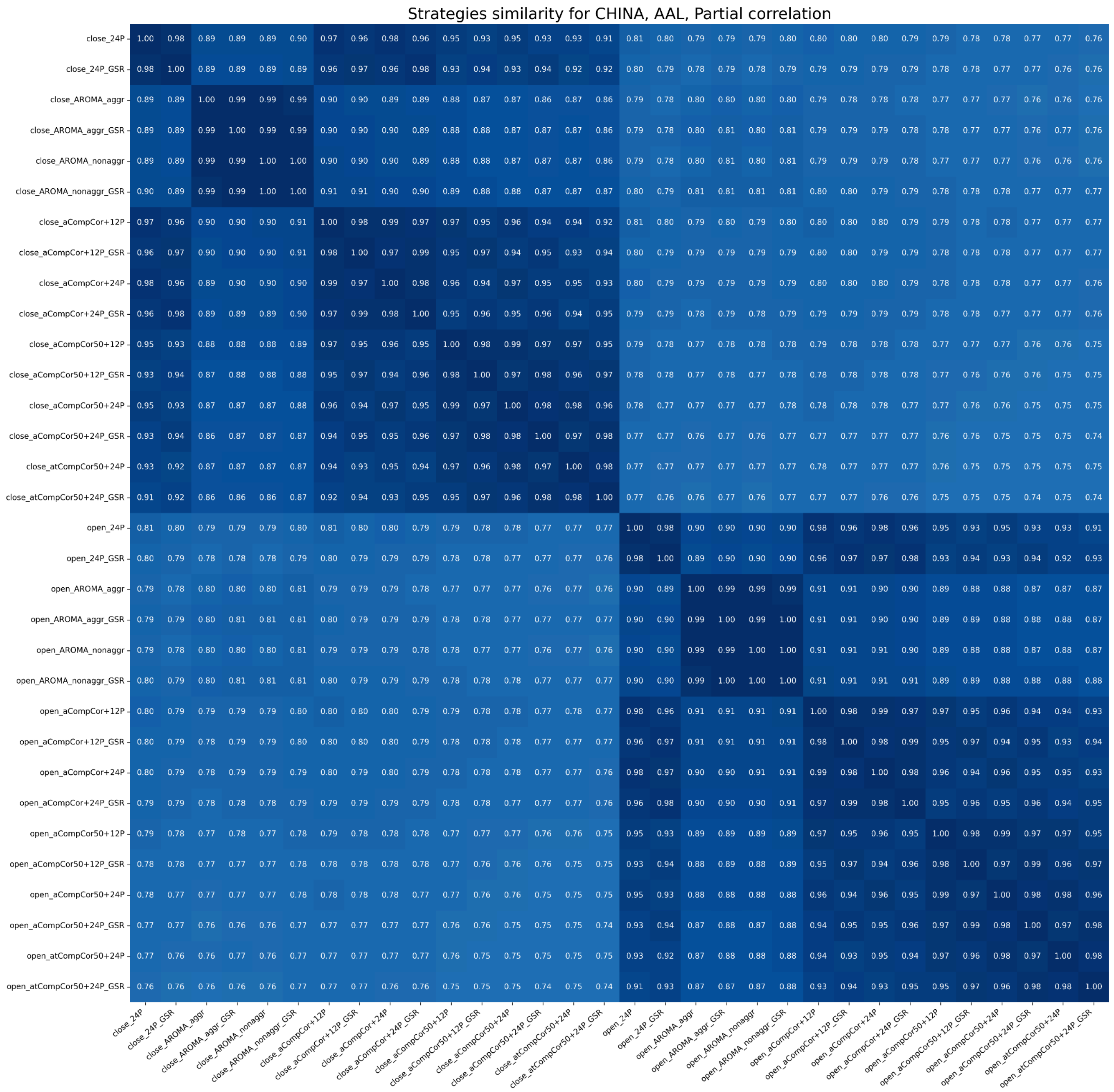


*Figure S7.*

*Similarity between signals of both EO and EC states denoised using different strategies for the China dataset, the AAL atlas, partial correlation FC. The heatmap shows similarity for one FC measure. The color range is from white (0, low similarity) to dark blue (1, high similarity). Mean values across subjects are presented.*

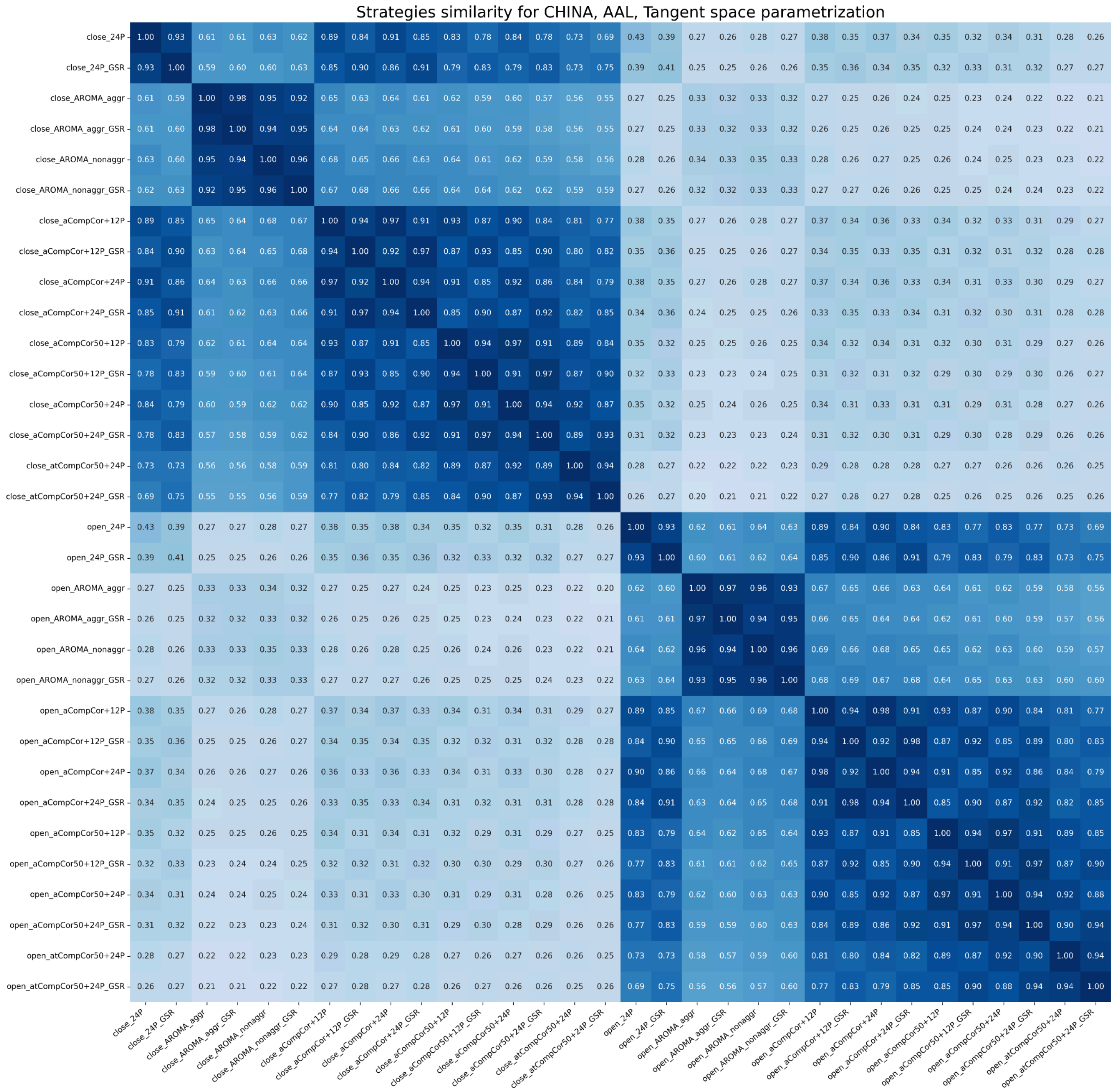


*Figure S8.*

*Similarity between signals of both EO and EC states denoised using different strategies for the China dataset, the AAL atlas, tangent space FC. The heatmap shows similarity for one FC measure. The color range is from white (0, low similarity) to dark blue (1, high similarity). Mean values across subjects are presented.*

### Brainnetome

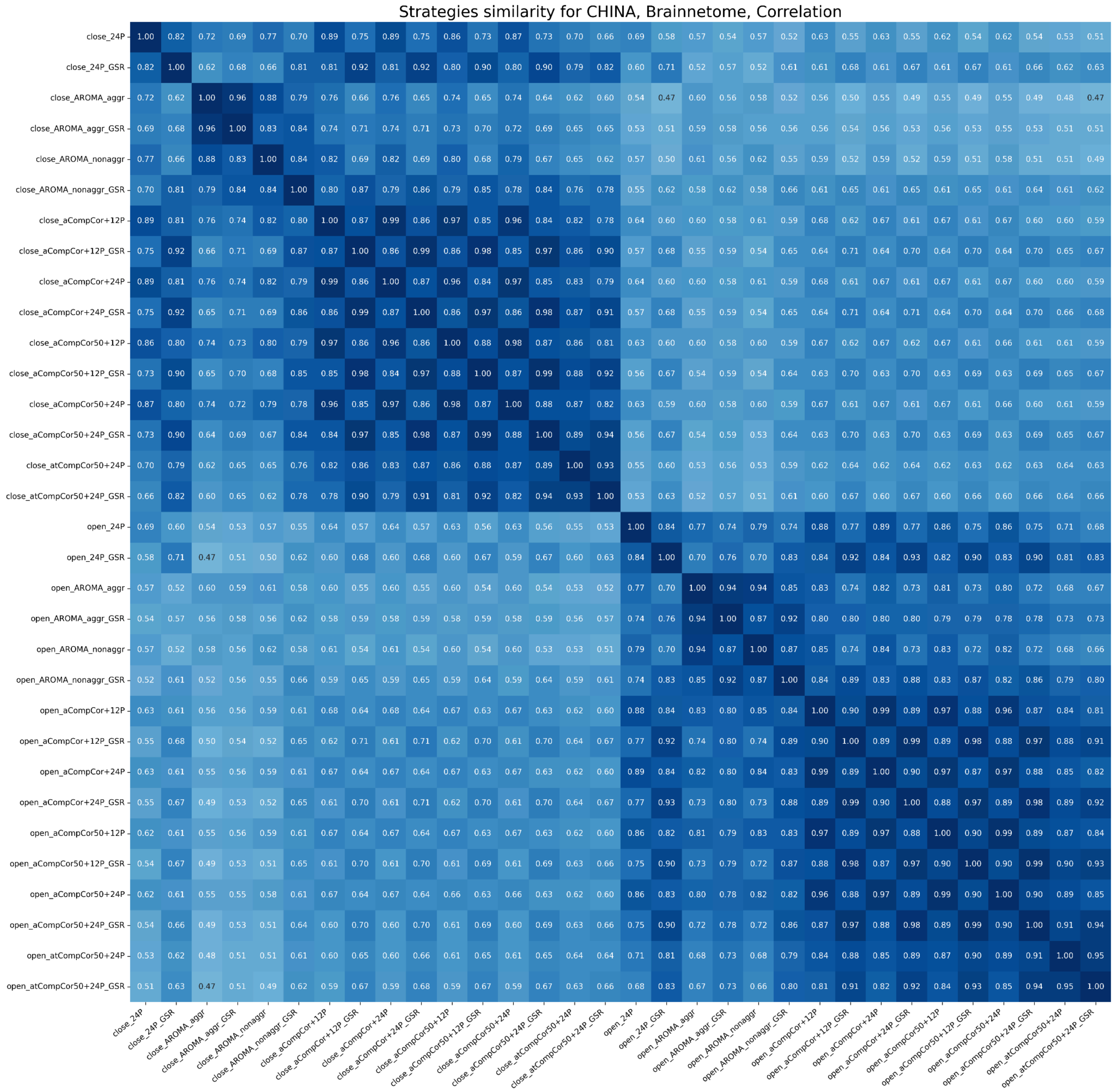


*Figure S9.*

*Similarity between signals of both EO and EC states denoised using different strategies for the China dataset, the Brainnetome atlas, correlation FC. The heatmap shows similarity for one FC measure. The color range is from white (0, low similarity) to dark blue (1, high similarity). Mean values across subjects are presented.*

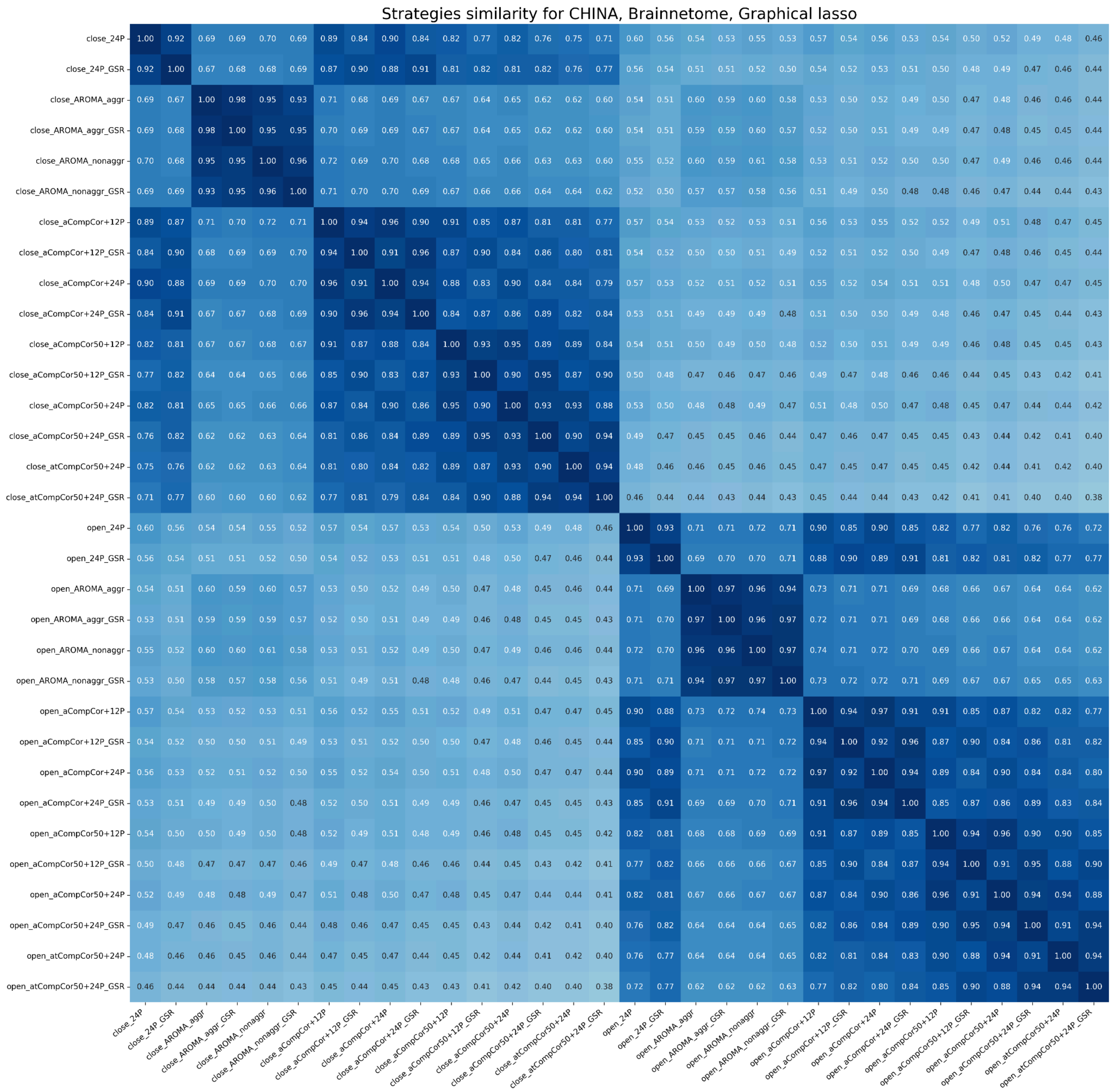


*Figure S10.*

*Similarity between signals of both EO and EC states denoised using different strategies for the China dataset, the Brainnetome atlas, glasso FC. The heatmap shows similarity for one FC measure. The color range is from white (0, low similarity) to dark blue (1, high similarity). Mean values across subjects are presented.*

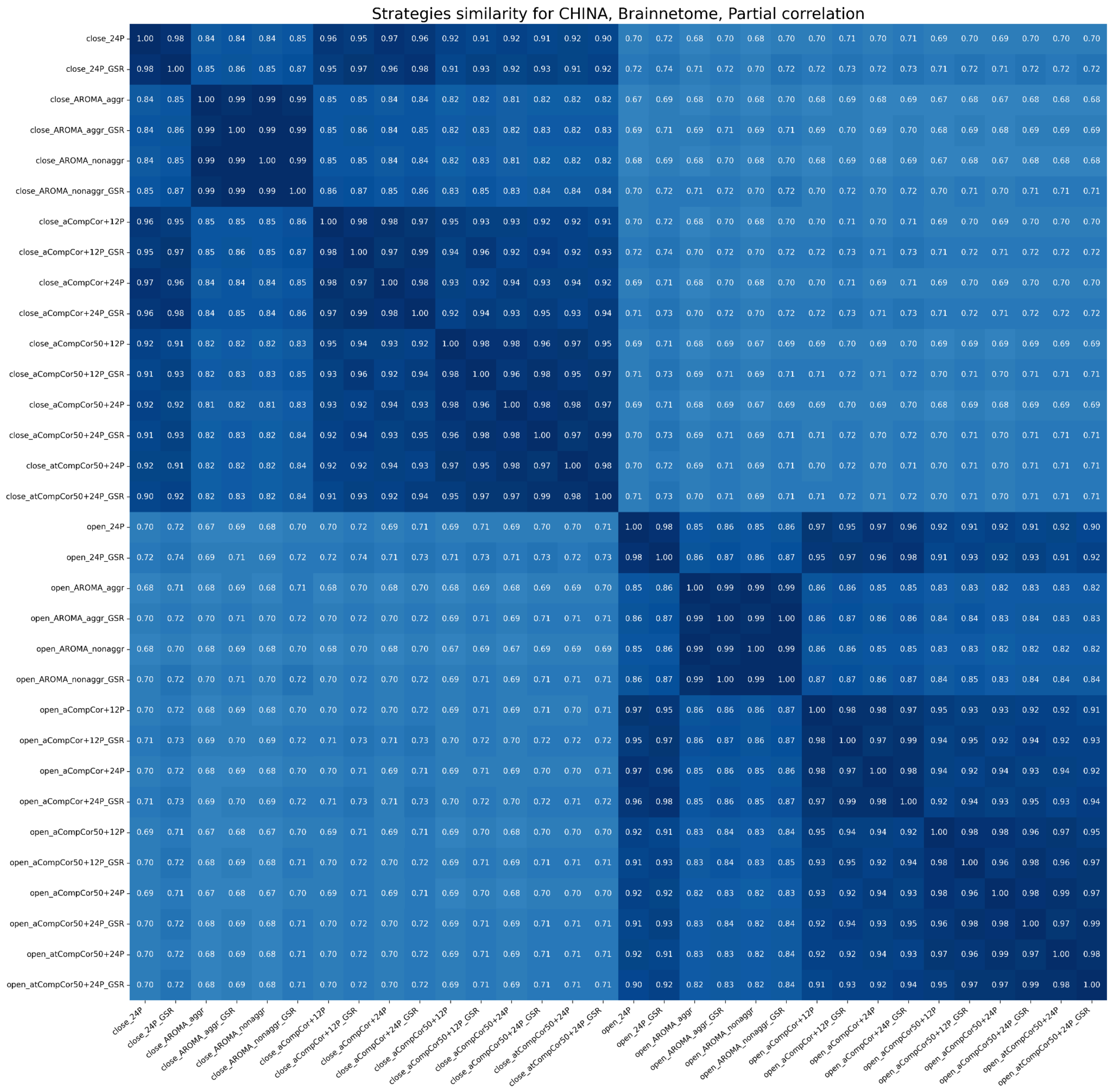


*Figure S11.*

*Similarity between signals of both EO and EC states denoised using different strategies for the China dataset, the Brainnetome atlas, partial correlation FC. The heatmap shows similarity for one FC measure. The color range is from white (0, low similarity) to dark blue (1, high similarity). Mean values across subjects are presented.*

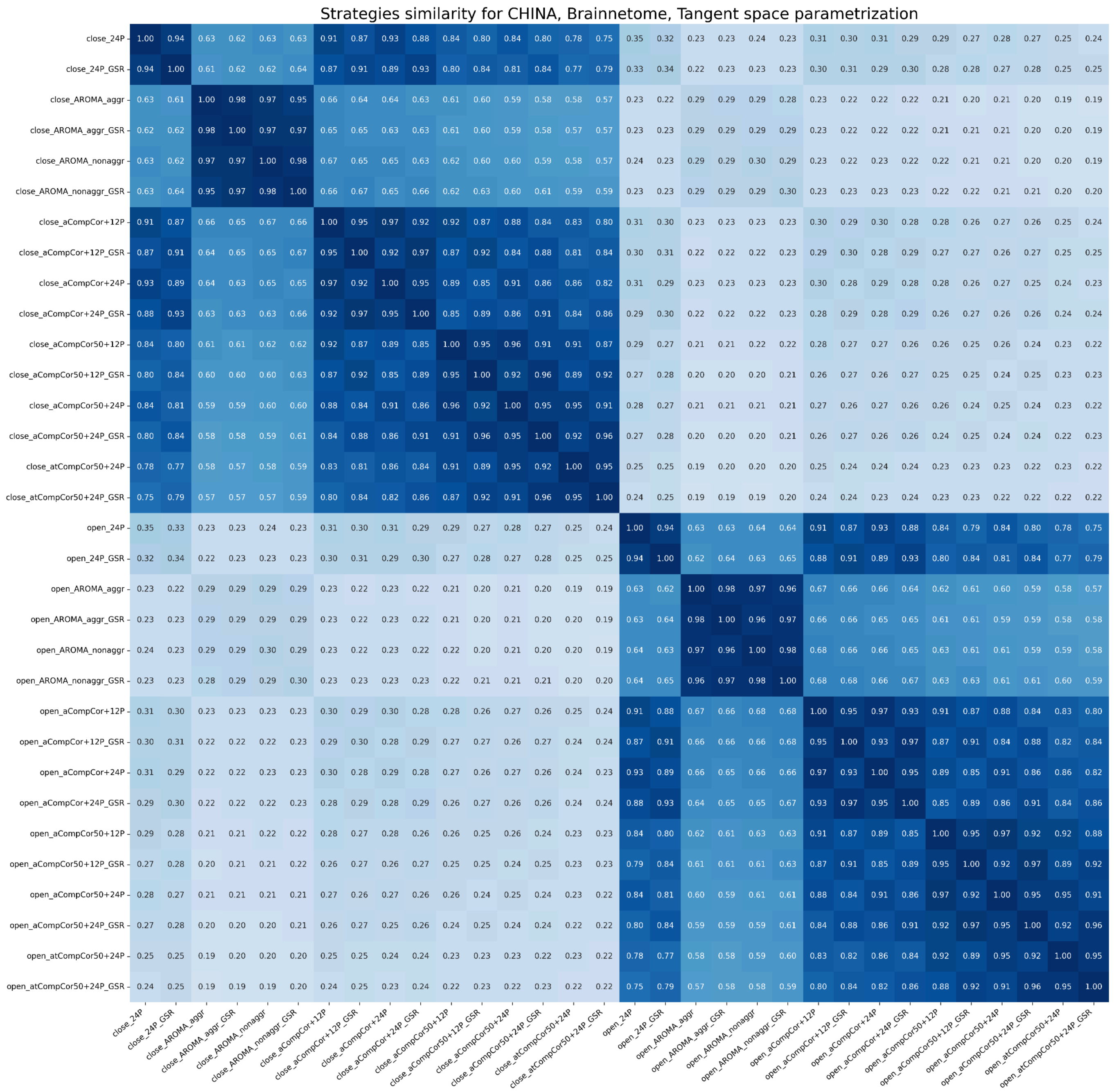


*Figure S12.*

*Similarity between signals of both EO and EC states denoised using different strategies for the China dataset, the Brainnetome atlas, tangent space FC. The heatmap shows similarity for one FC measure. The color range is from white (0, low similarity) to dark blue (1, high similarity). Mean values across subjects are presented.*

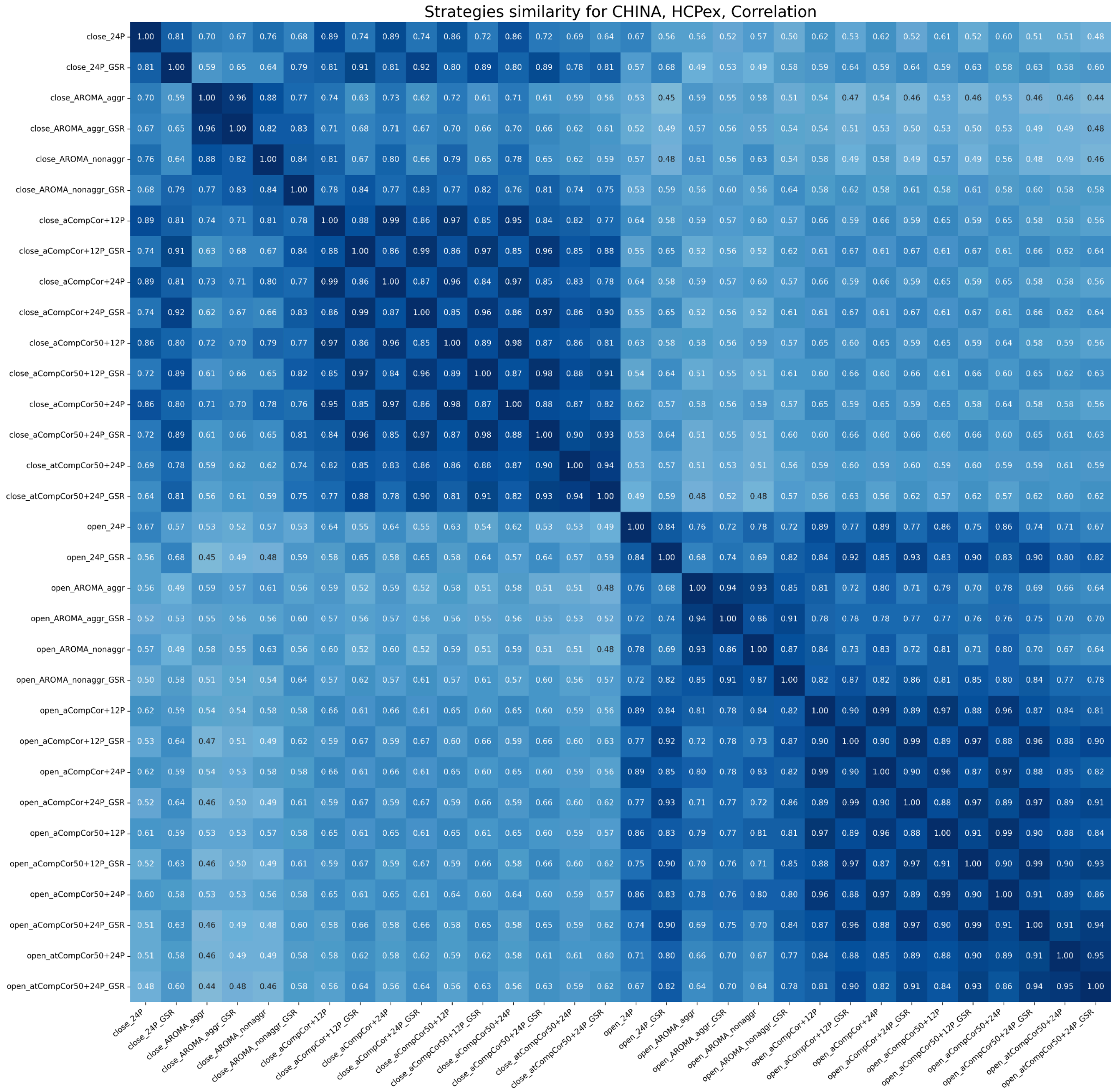


*Figure S13.*

*Similarity between signals of both EO and EC states denoised using different strategies for the China dataset, the HCPex atlas, correlation FC. The heatmap shows similarity for one FC measure. The color range is from white (0, low similarity) to dark blue (1, high similarity). Mean values across subjects are presented.*

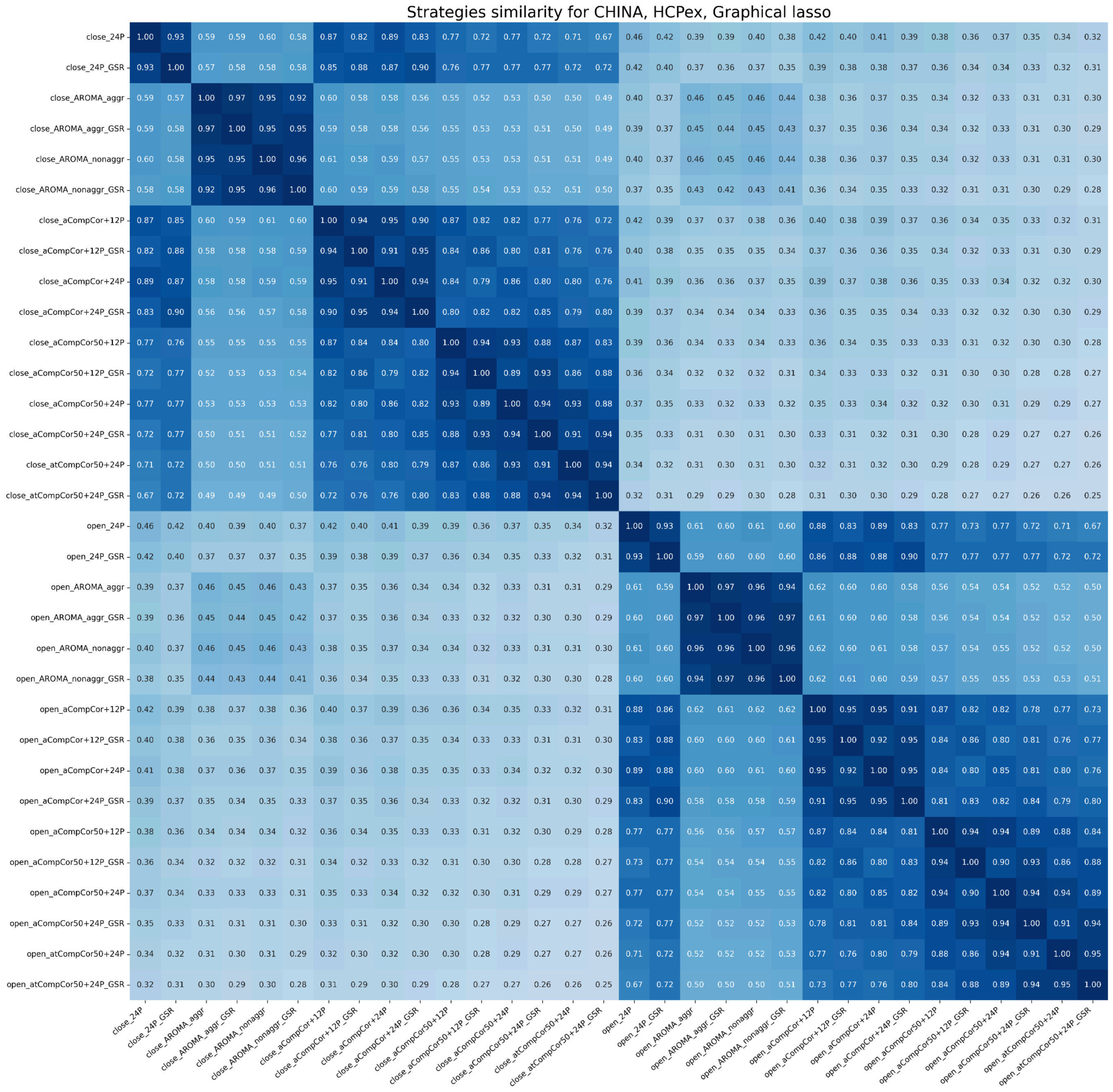


*Figure S14.*

*Similarity between signals of both EO and EC states denoised using different strategies for the China dataset, the HCPex atlas, correlation FC. The heatmap shows similarity for one FC measure. The color range is from white (0, low similarity) to dark blue (1, high similarity). Mean values across subjects are presented.*

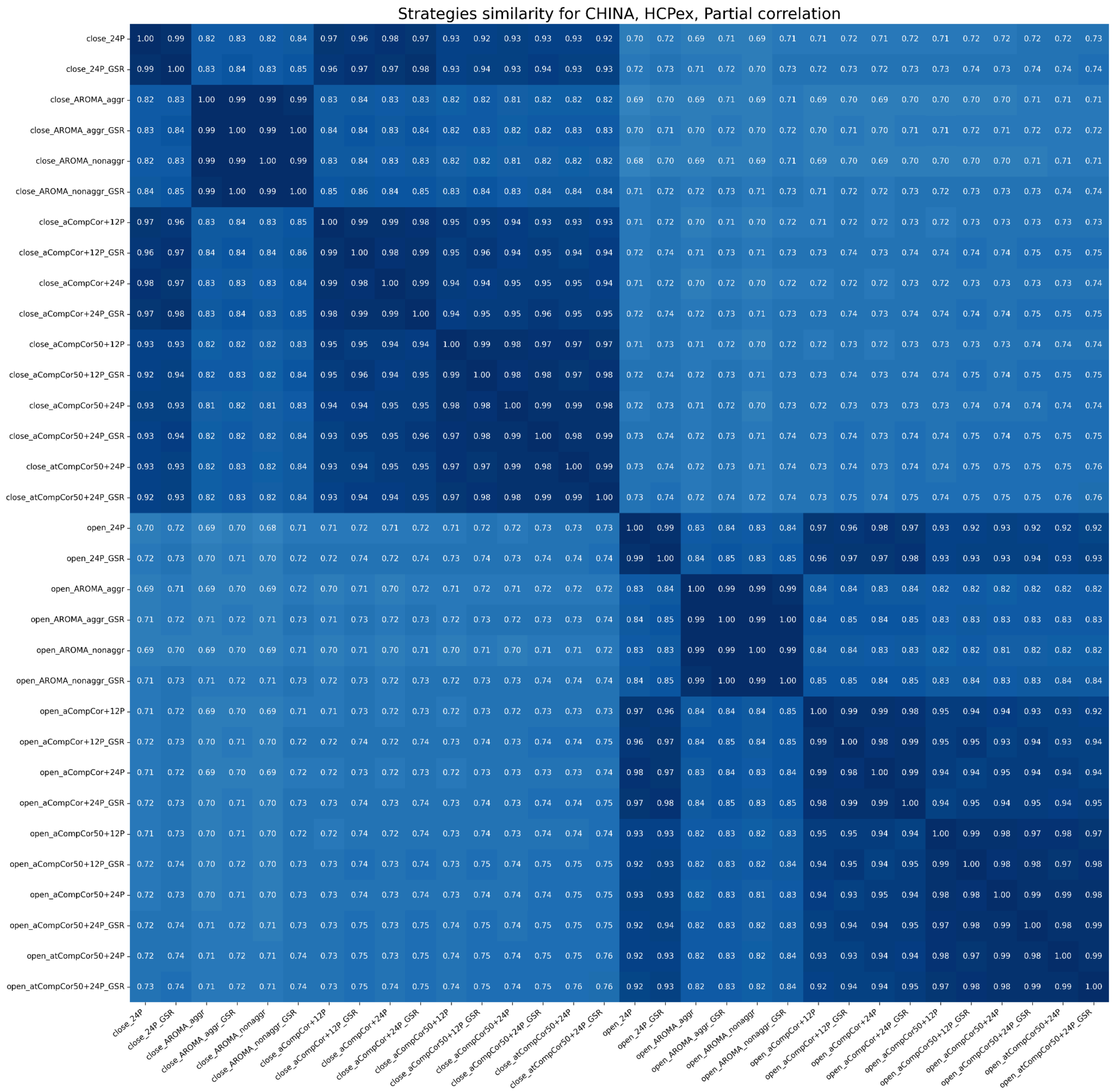


*Figure S15.*

*Similarity between signals of both EO and EC states denoised using different strategies for the China dataset, the HCPex atlas, partial correlation FC. The heatmap shows similarity for one FC measure. The color range is from white (0, low similarity) to dark blue (1, high similarity). Mean values across subjects are presented.*

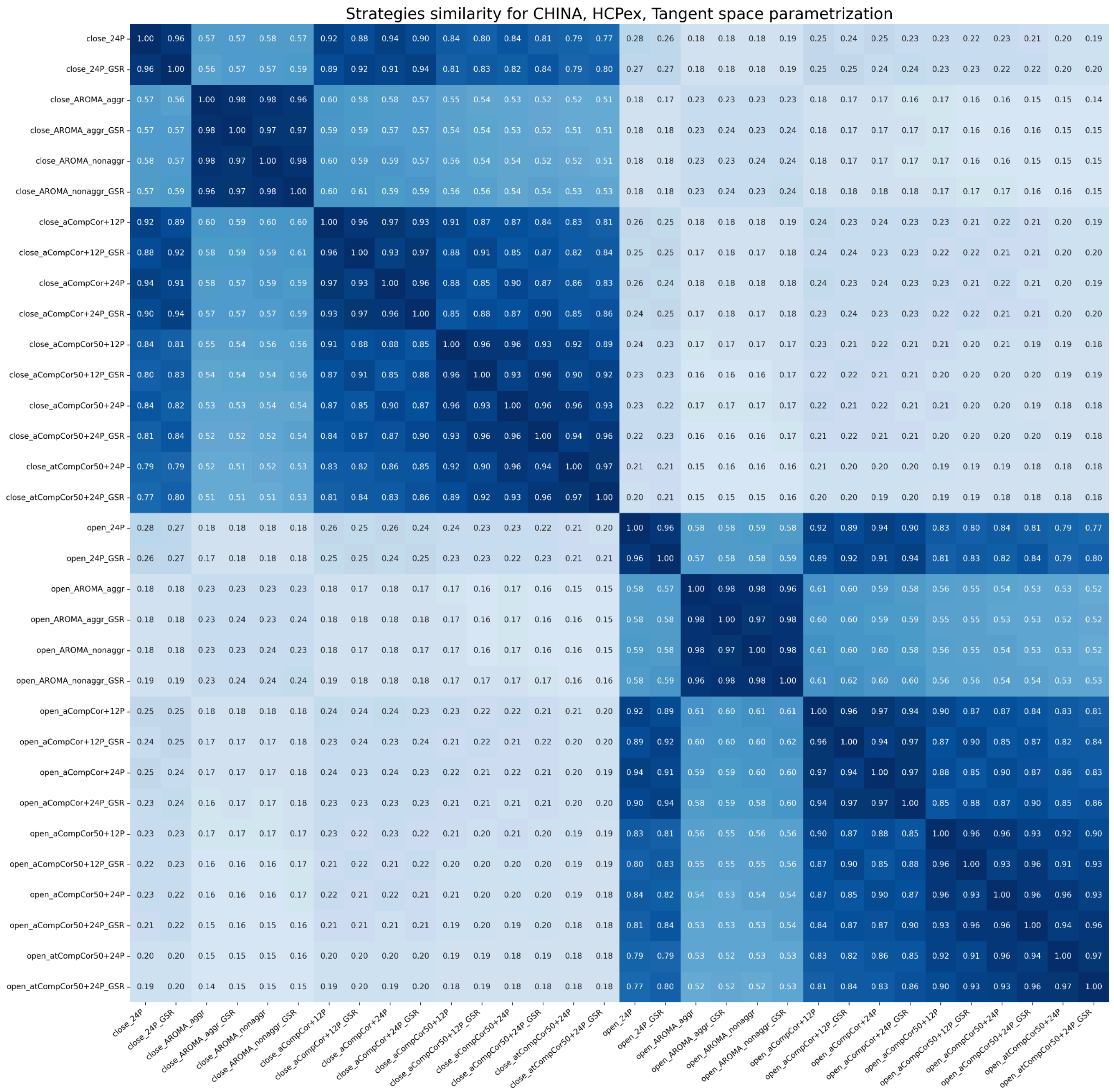


*Figure S16.*

*Similarity between signals of both EO and EC states denoised using different strategies for the China dataset, the HCPex atlas, tangent space FC. The heatmap shows similarity for one FC measure. The color range is from white (0, low similarity) to dark blue (1, high similarity). Mean values across subjects are presented.*

### Schaefer200

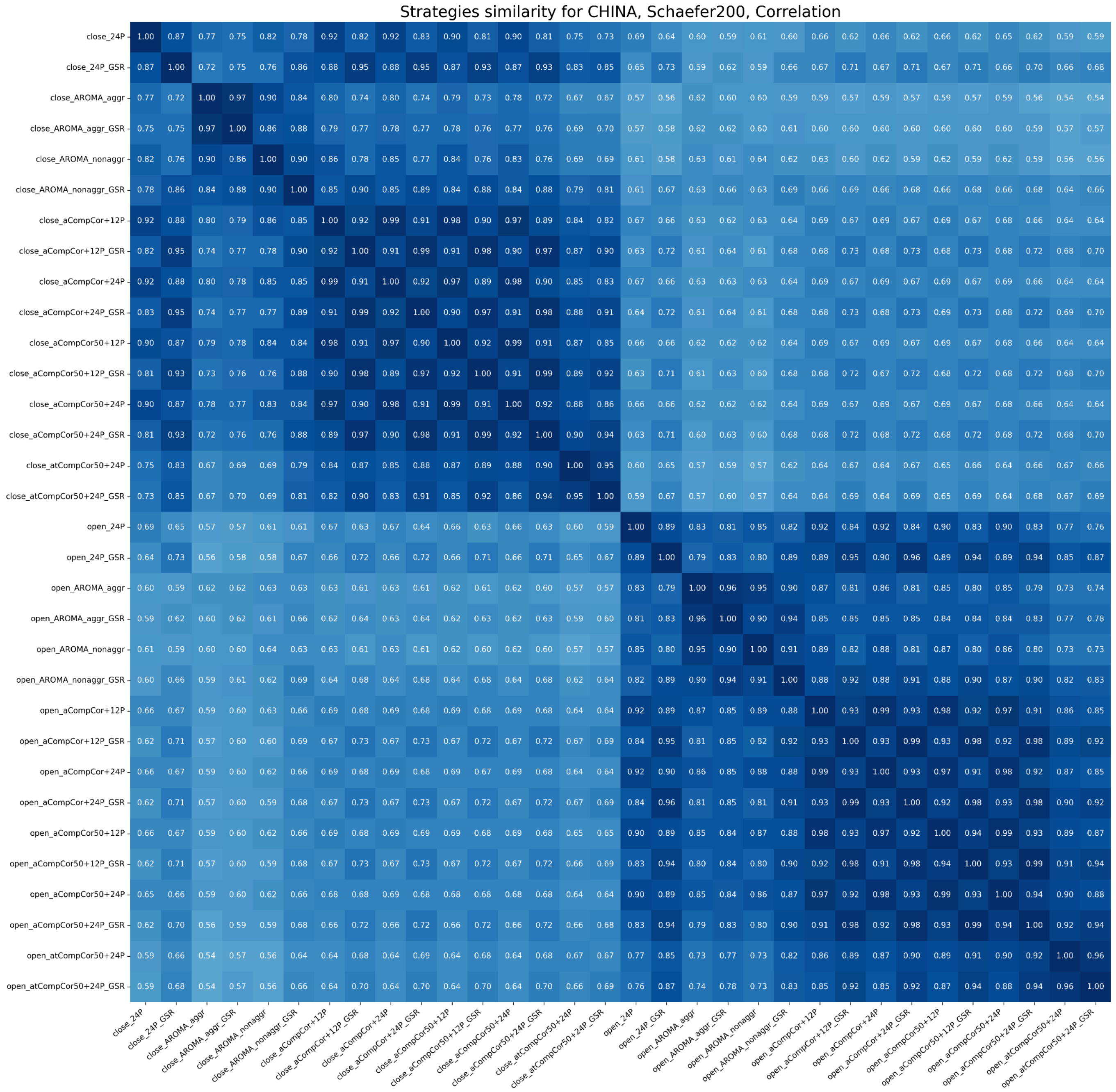


*Figure S17.*

*Similarity between signals of both EO and EC states denoised using different strategies for the China dataset, the Schaefer200 atlas, correlation FC. The heatmap shows similarity for one FC measure. The color range is from white (0, low similarity) to dark blue (1, high similarity). Mean values across subjects are presented.*

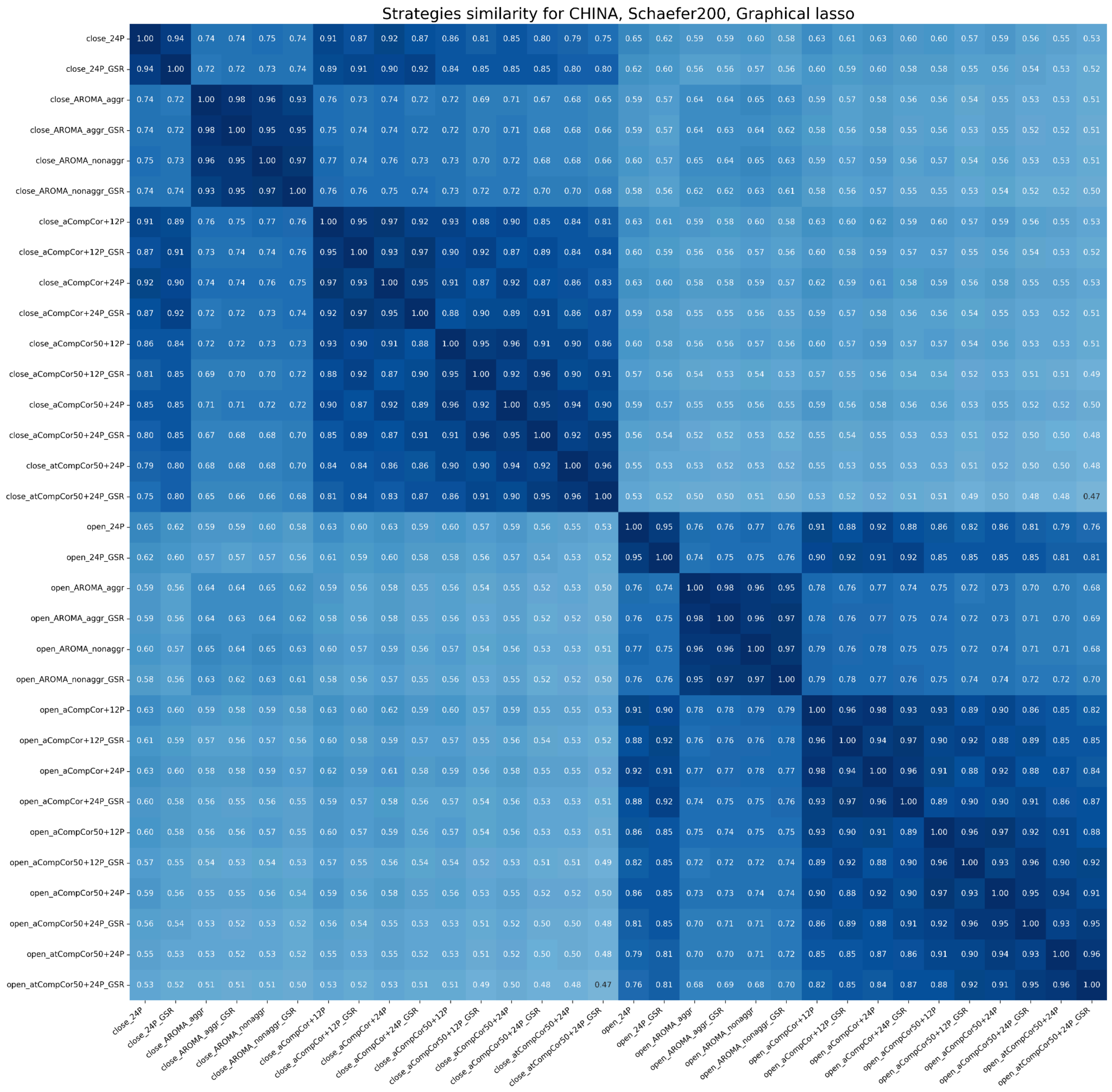


*Figure S18.*

*Similarity between signals of both EO and EC states denoised using different strategies for the China dataset, the Schaefer200 atlas, glasso FC. The heatmap shows similarity for one FC measure. The color range is from white (0, low similarity) to dark blue (1, high similarity). Mean values across subjects are presented.*

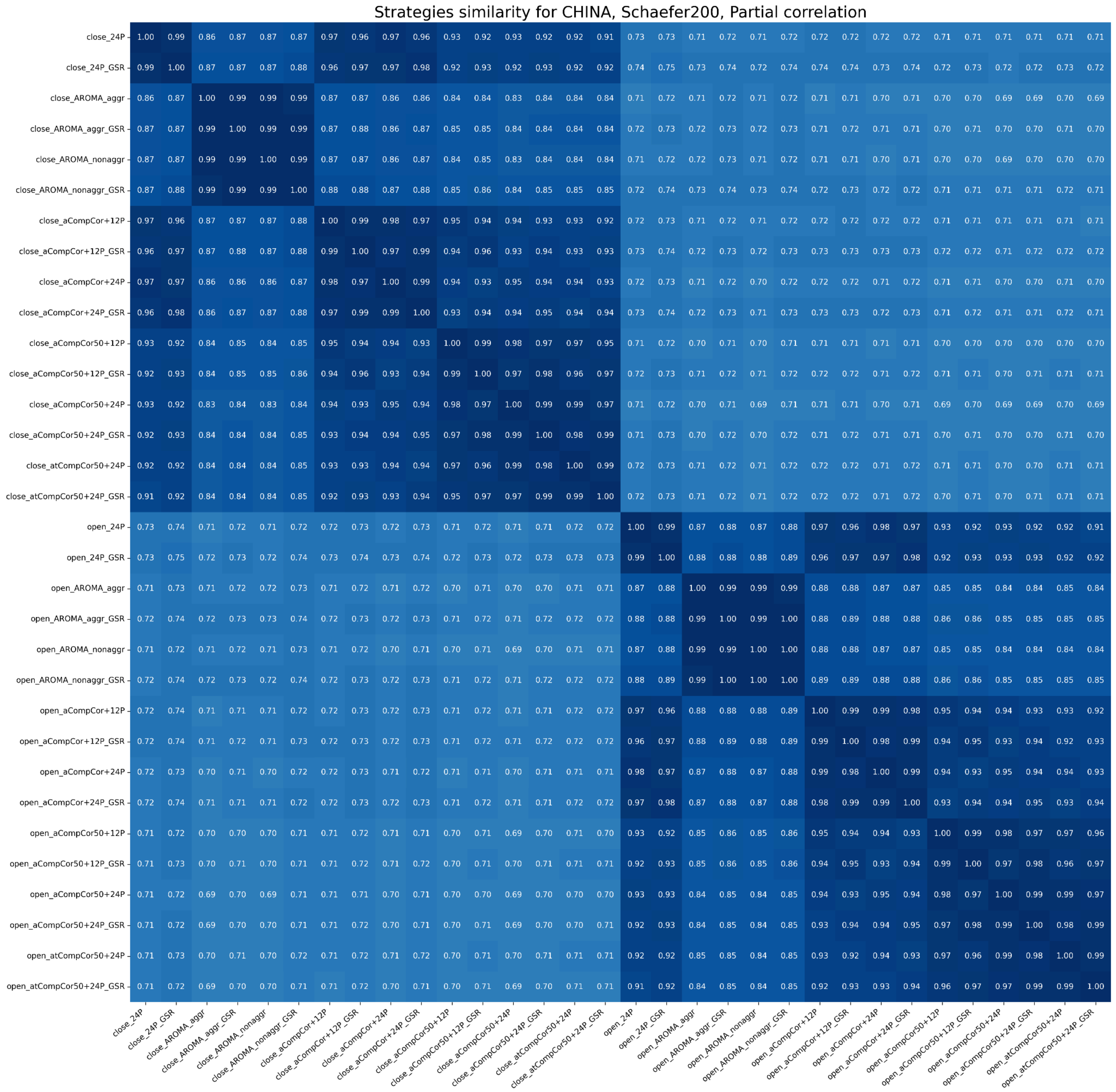


*Figure S19.*

*Similarity between signals of both EO and EC states denoised using different strategies for the China dataset, the Schaefer200 atlas, partial correlation FC. The heatmap shows similarity for one FC measure. The color range is from white (0, low similarity) to dark blue (1, high similarity). Mean values across subjects are presented.*

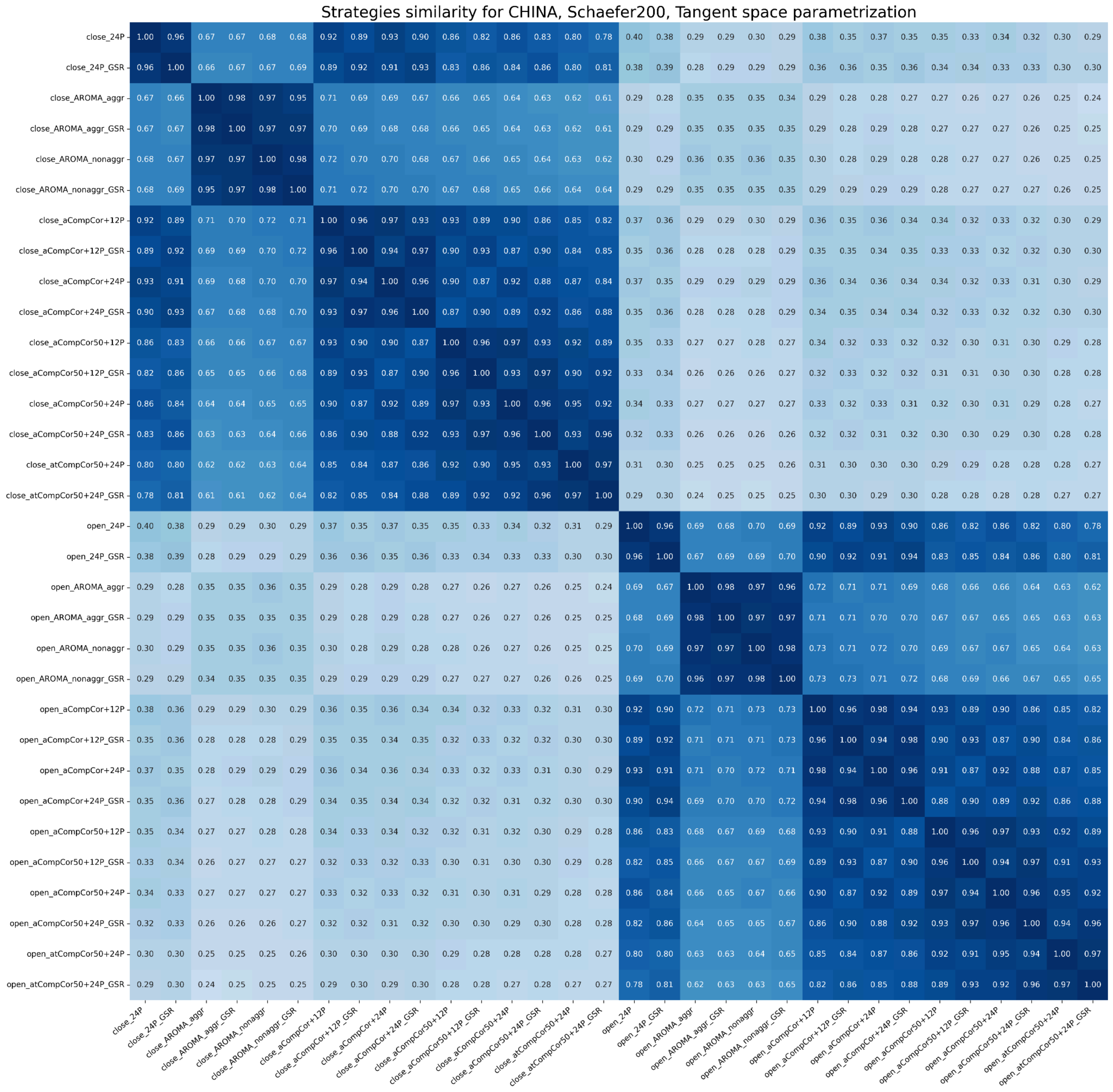


*Figure S20.*

*Similarity between signals of both EO and EC states denoised using different strategies for the China dataset, the Schaefer200 atlas, tangent space FC. The heatmap shows similarity for one FC measure. The color range is from white (0, low similarity) to dark blue (1, high similarity). Mean values across subjects are presented.*

### IHB dataset

#### AAL

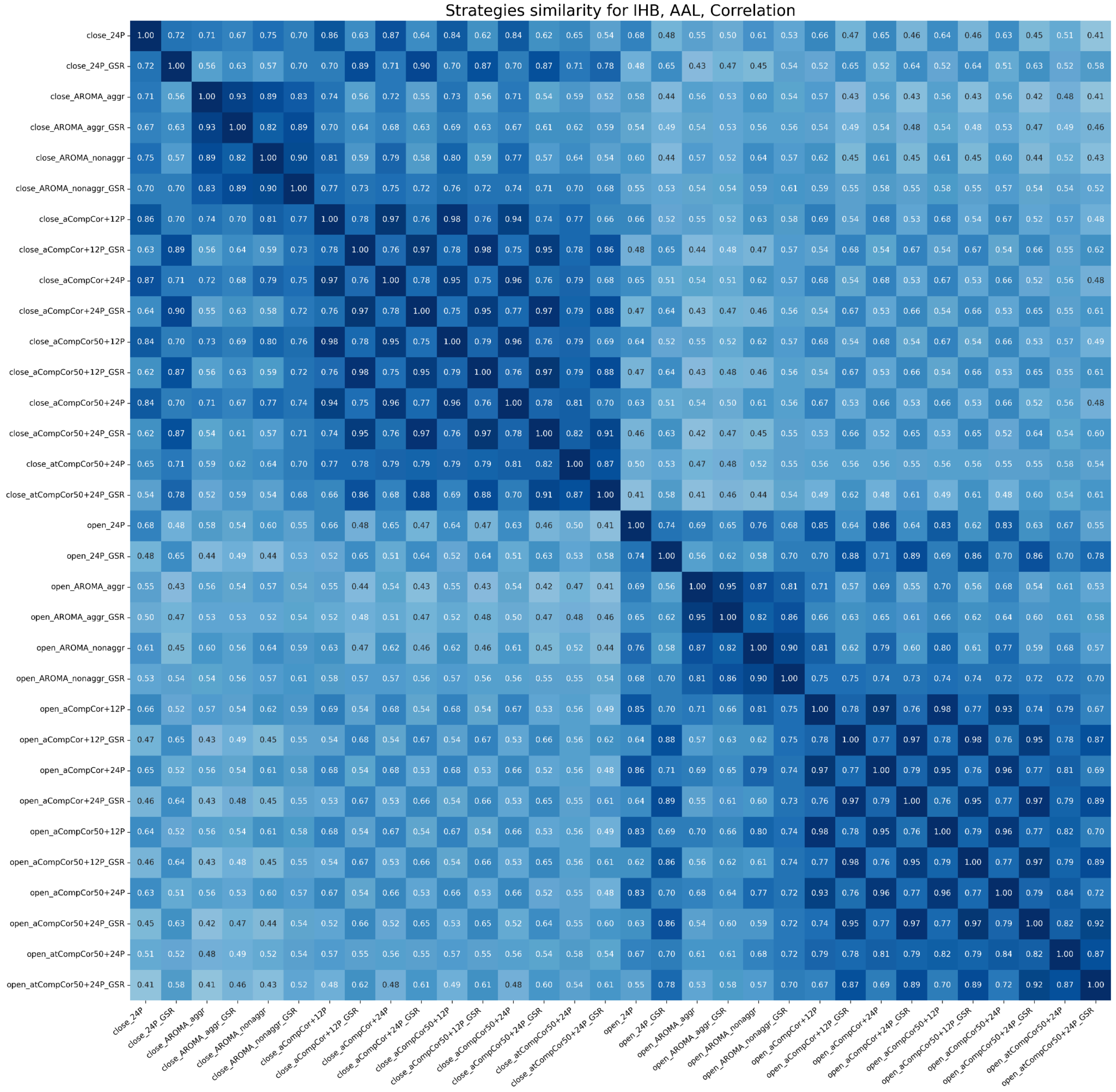


*Figure S21.*

*Similarity between signals of both EO and EC states denoised using different strategies for the IHB dataset, the AAL atlas, correlation FC. The heatmap shows similarity for one FC measure. The color range is from white (0, low similarity) to dark blue (1, high similarity). Mean values across subjects are presented.*

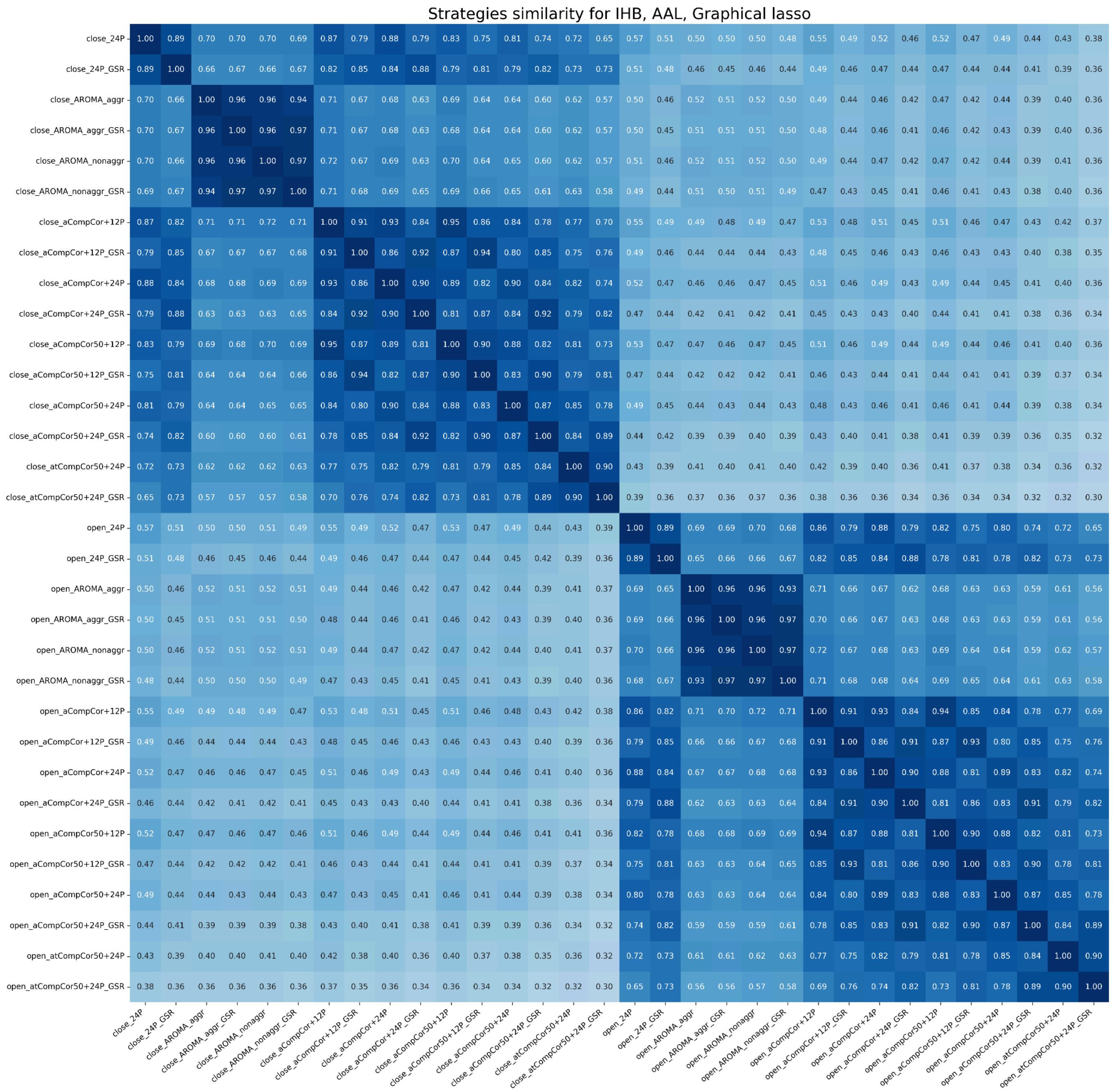


*Figure S22.*

*Similarity between signals of both EO and EC states denoised using different strategies for the IHB dataset, the AAL atlas, glasso FC. The heatmap shows similarity for one FC measure. The color range is from white (0, low similarity) to dark blue (1, high similarity). Mean values across subjects are presented.*

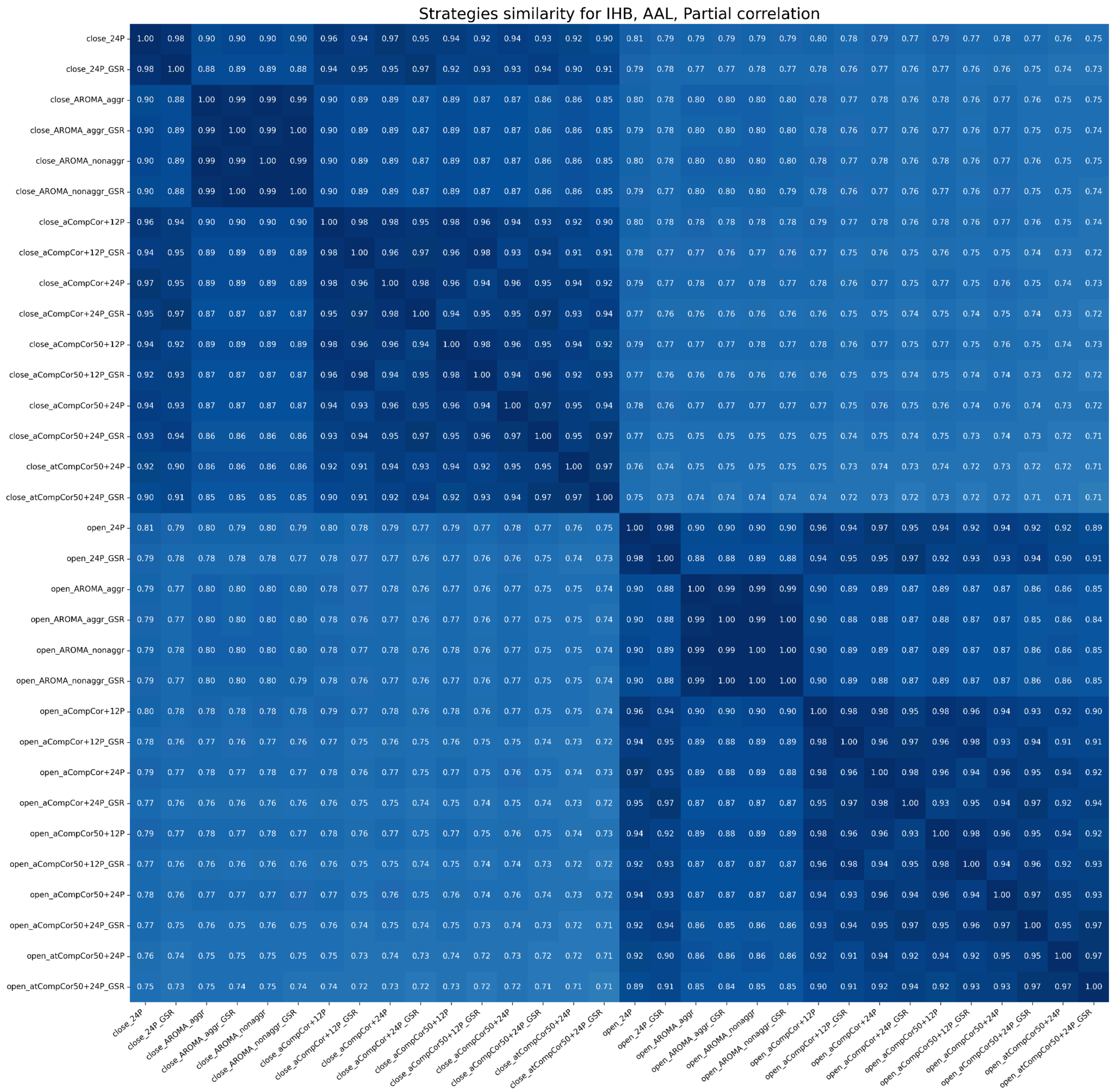


*Figure S23.*

*Similarity between signals of both EO and EC states denoised using different strategies for the IHB dataset, the AAL atlas, partial correlation FC. The heatmap shows similarity for one FC measure. The color range is from white (0, low similarity) to dark blue (1, high similarity). Mean values across subjects are presented.*

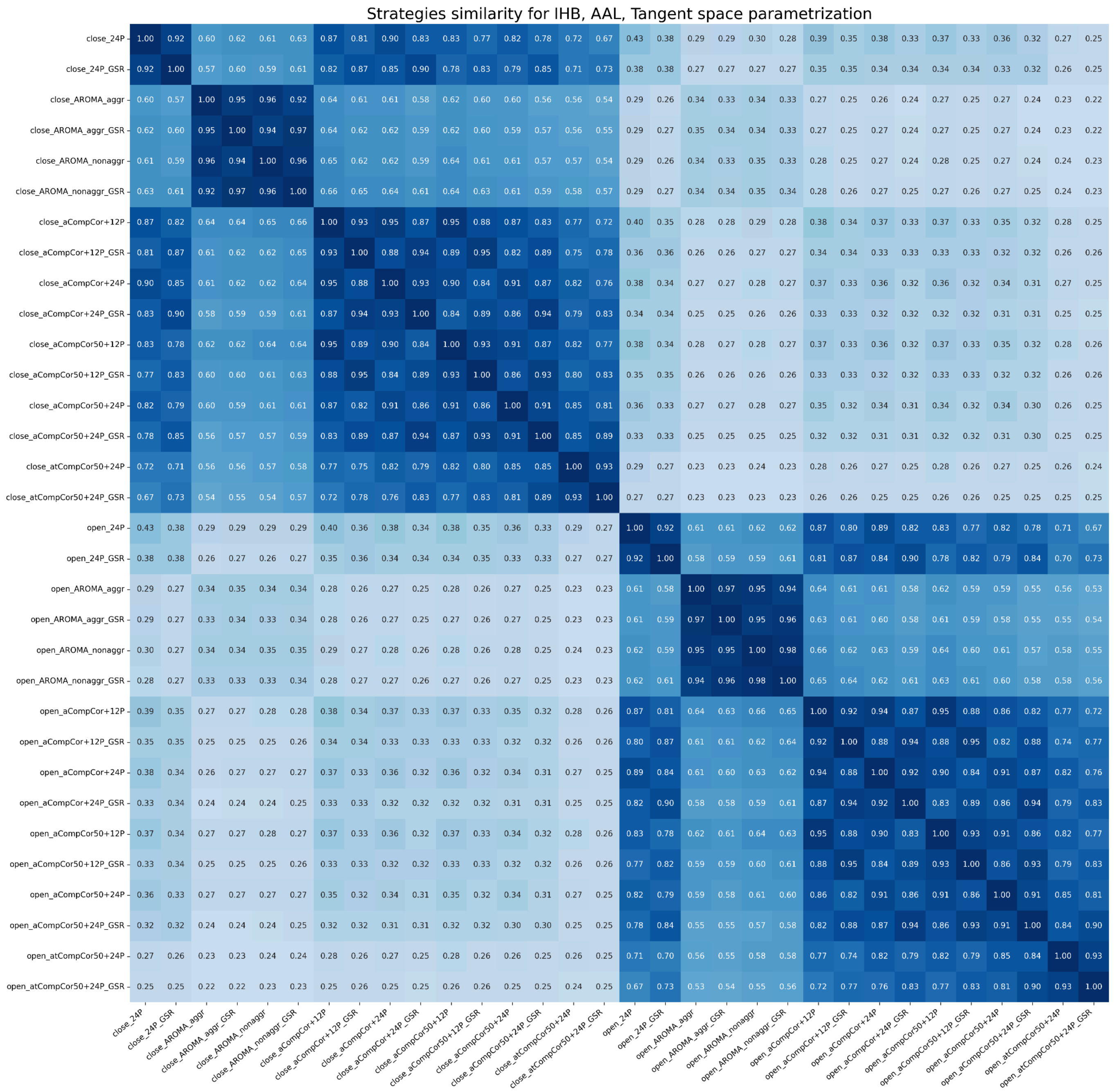


*Figure S24.*

*Similarity between signals of both EO and EC states denoised using different strategies for the IHB dataset, the AAL atlas, tangent space FC. The heatmap shows similarity for one FC measure. The color range is from white (0, low similarity) to dark blue (1, high similarity). Mean values across subjects are presented.*

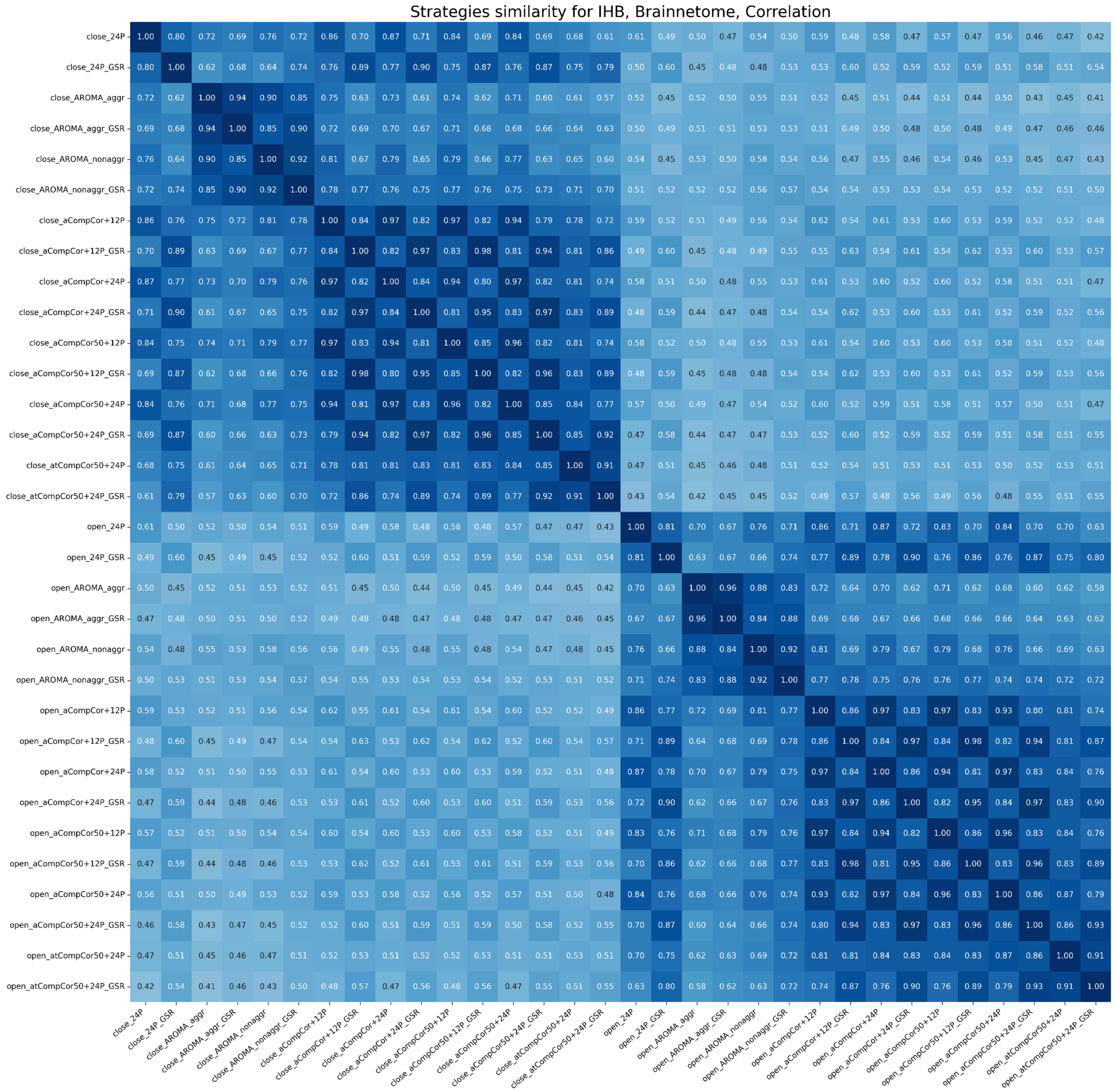


*Figure S25.*

*Similarity between signals of both EO and EC states denoised using different strategies for the IHB dataset, the Brainnetome atlas, correlation FC. The heatmap shows similarity for one FC measure. The color range is from white (0, low similarity) to dark blue (1, high similarity). Mean values across subjects are presented.*

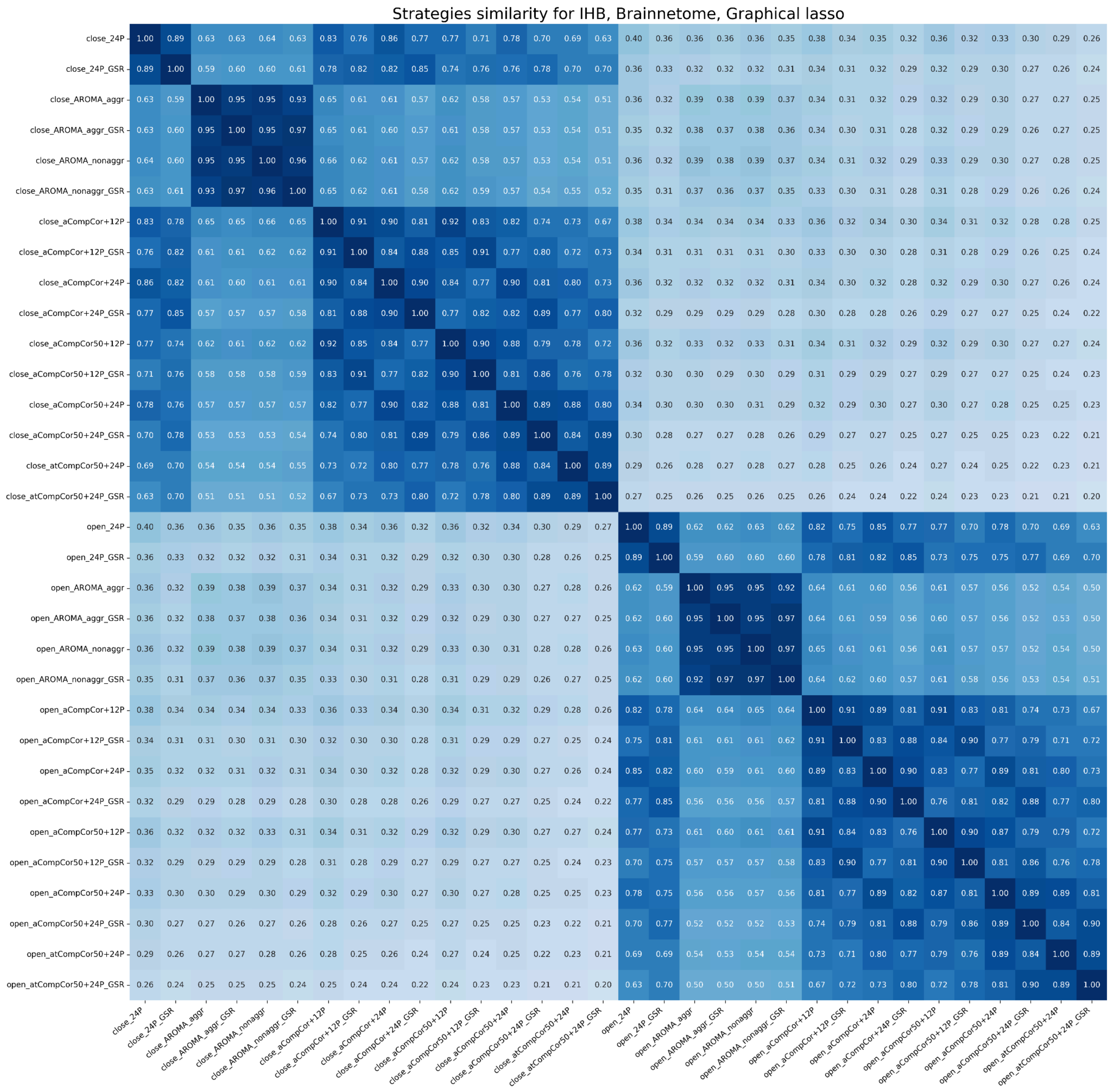


*Figure S26.*

*Similarity between signals of both EO and EC states denoised using different strategies for the IHB dataset, the Brainnetome atlas, glasso FC. The heatmap shows similarity for one FC measure. The color range is from white (0, low similarity) to dark blue (1, high similarity). Mean values across subjects are presented.*

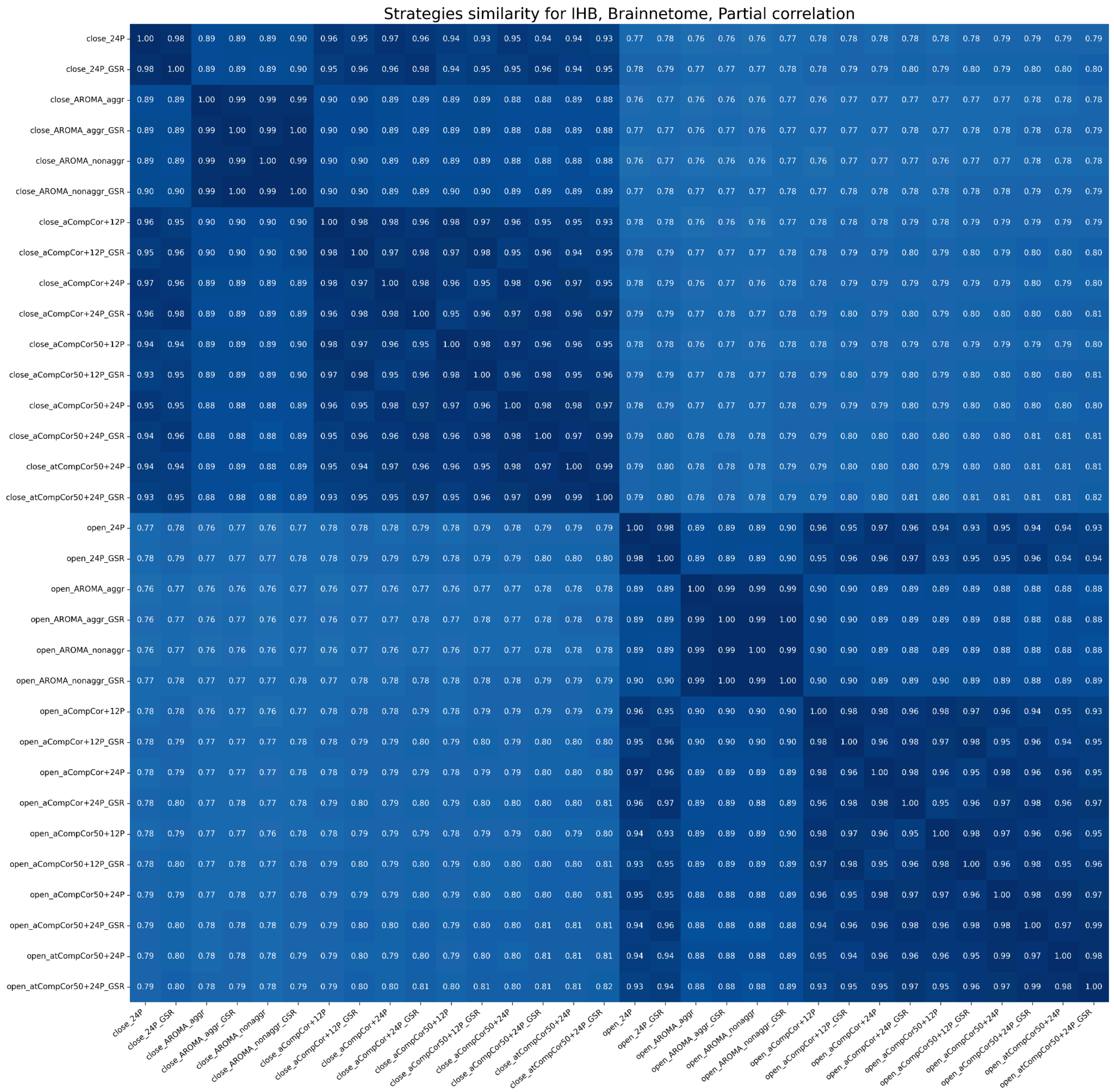


*Figure S27.*

*Similarity between signals of both EO and EC states denoised using different strategies for the IHB dataset, the Brainnetome atlas, partial correlation FC. The heatmap shows similarity for one FC measure. The color range is from white (0, low similarity) to dark blue (1, high similarity). Mean values across subjects are presented.*

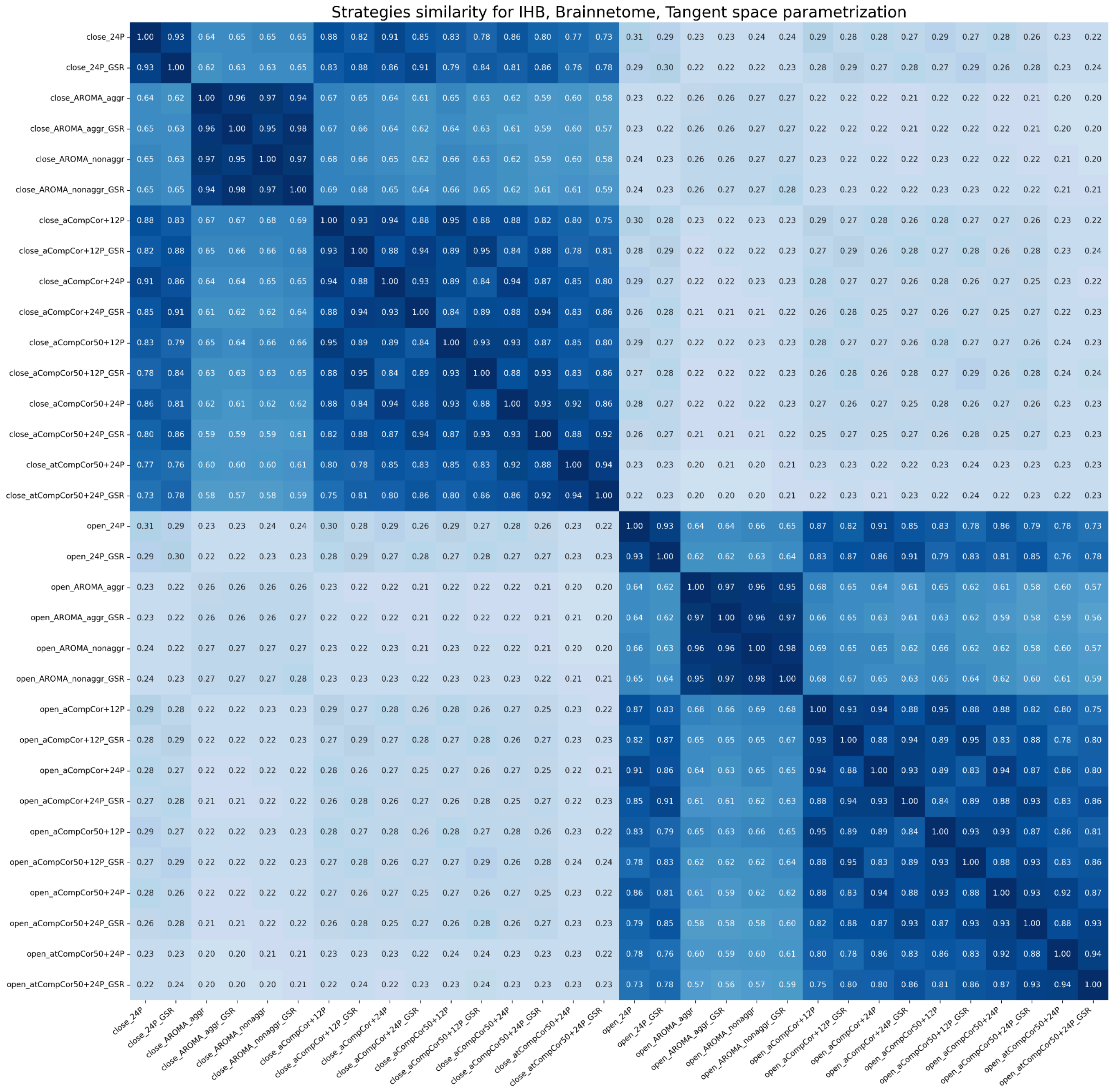


*Figure S28.*

*Similarity between signals of both EO and EC states denoised using different strategies for the IHB dataset, the Brainnetome atlas, tangent space FC. The heatmap shows similarity for one FC measure. The color range is from white (0, low similarity) to dark blue (1, high similarity). Mean values across subjects are presented.*

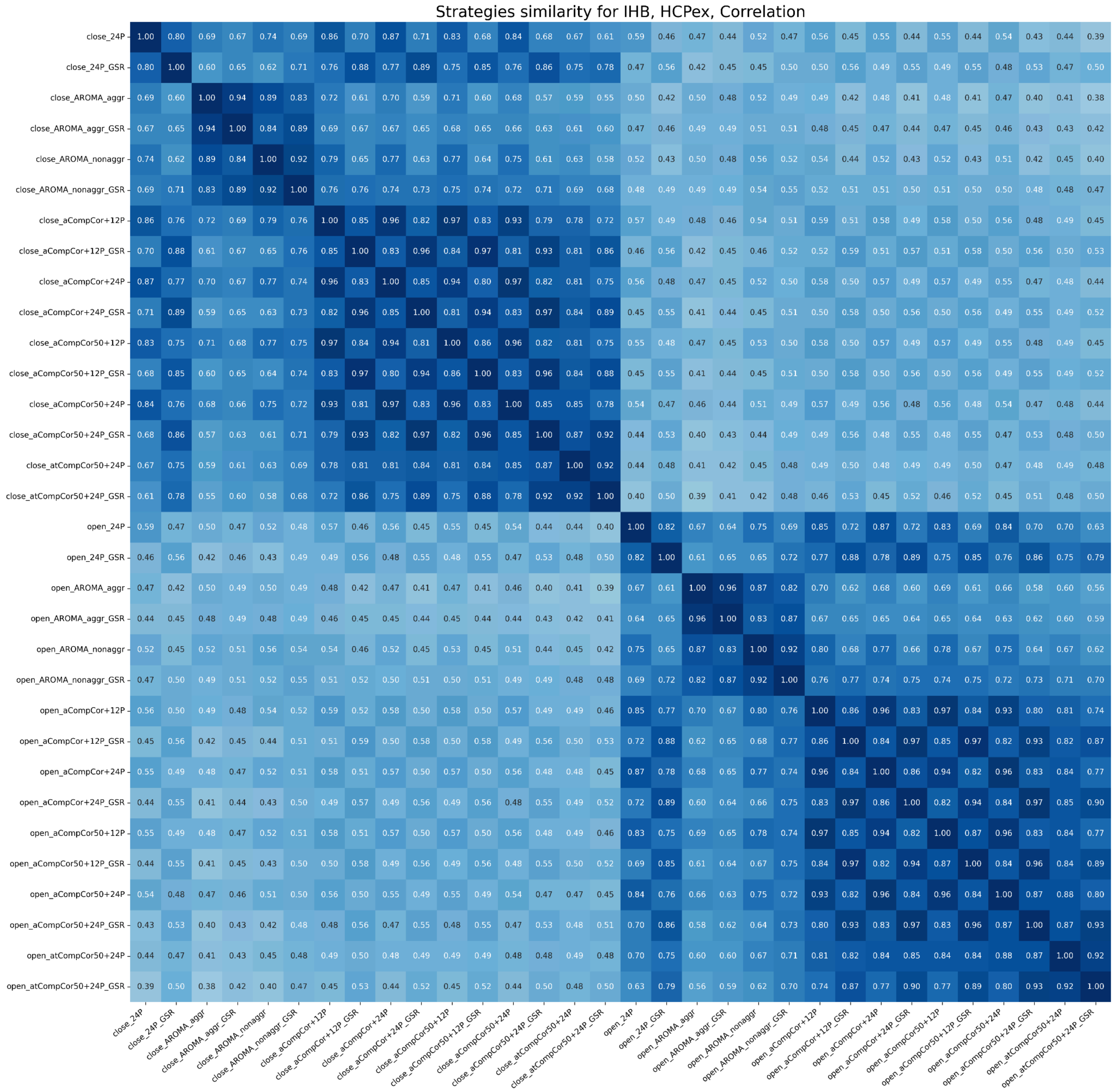


*Figure S29.*

*Similarity between signals of both EO and EC states denoised using different strategies for the IHB dataset, the HCPex atlas, correlation FC. The heatmap shows similarity for one FC measure. The color range is from white (0, low similarity) to dark blue (1, high similarity). Mean values across subjects are presented.*

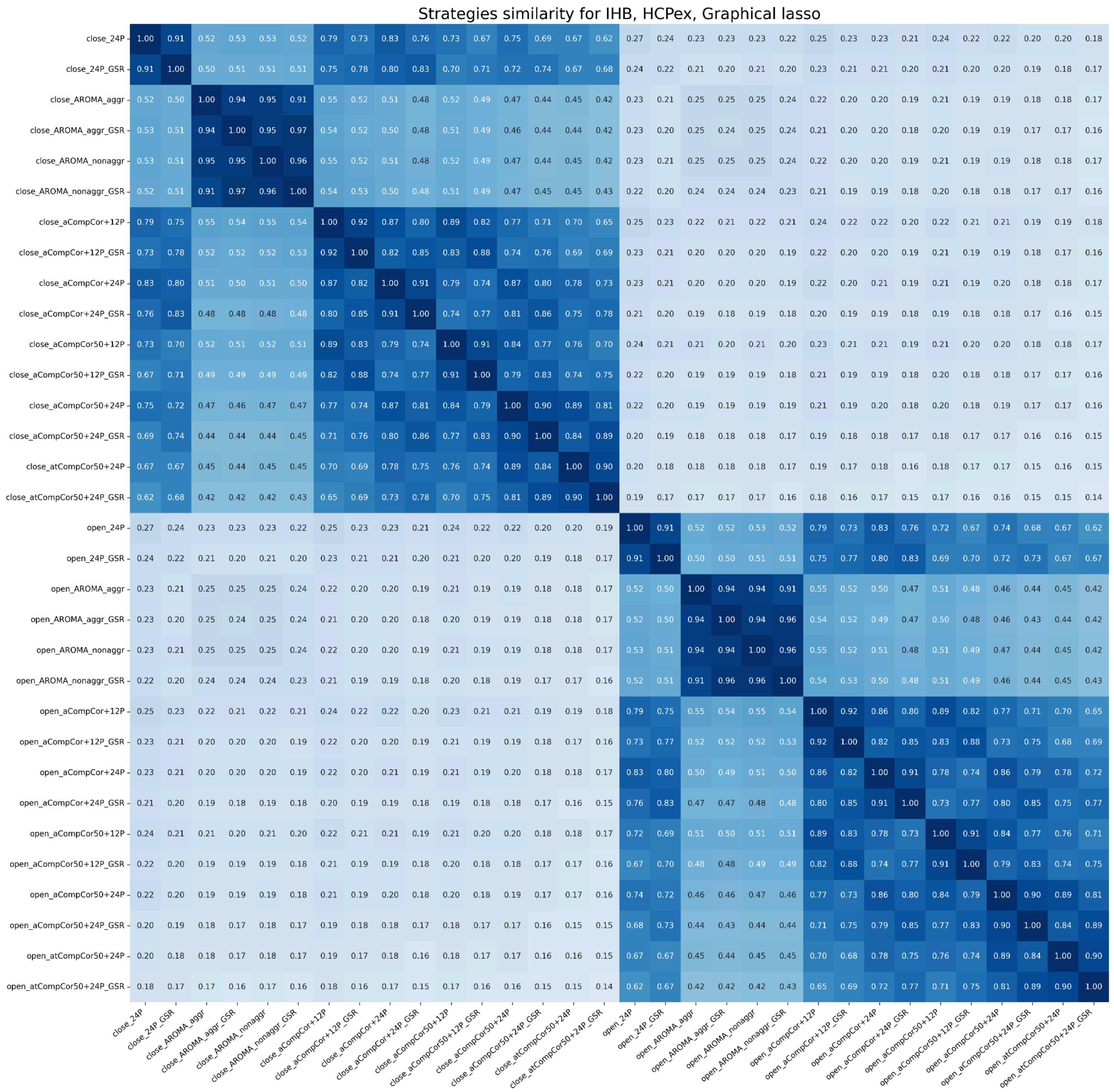


*Figure S30.*

*Similarity between signals of both EO and EC states denoised using different strategies for the IHB dataset, the HCPex atlas, glasso FC. The heatmap shows similarity for one FC measure. The color range is from white (0, low similarity) to dark blue (1, high similarity). Mean values across subjects are presented.*

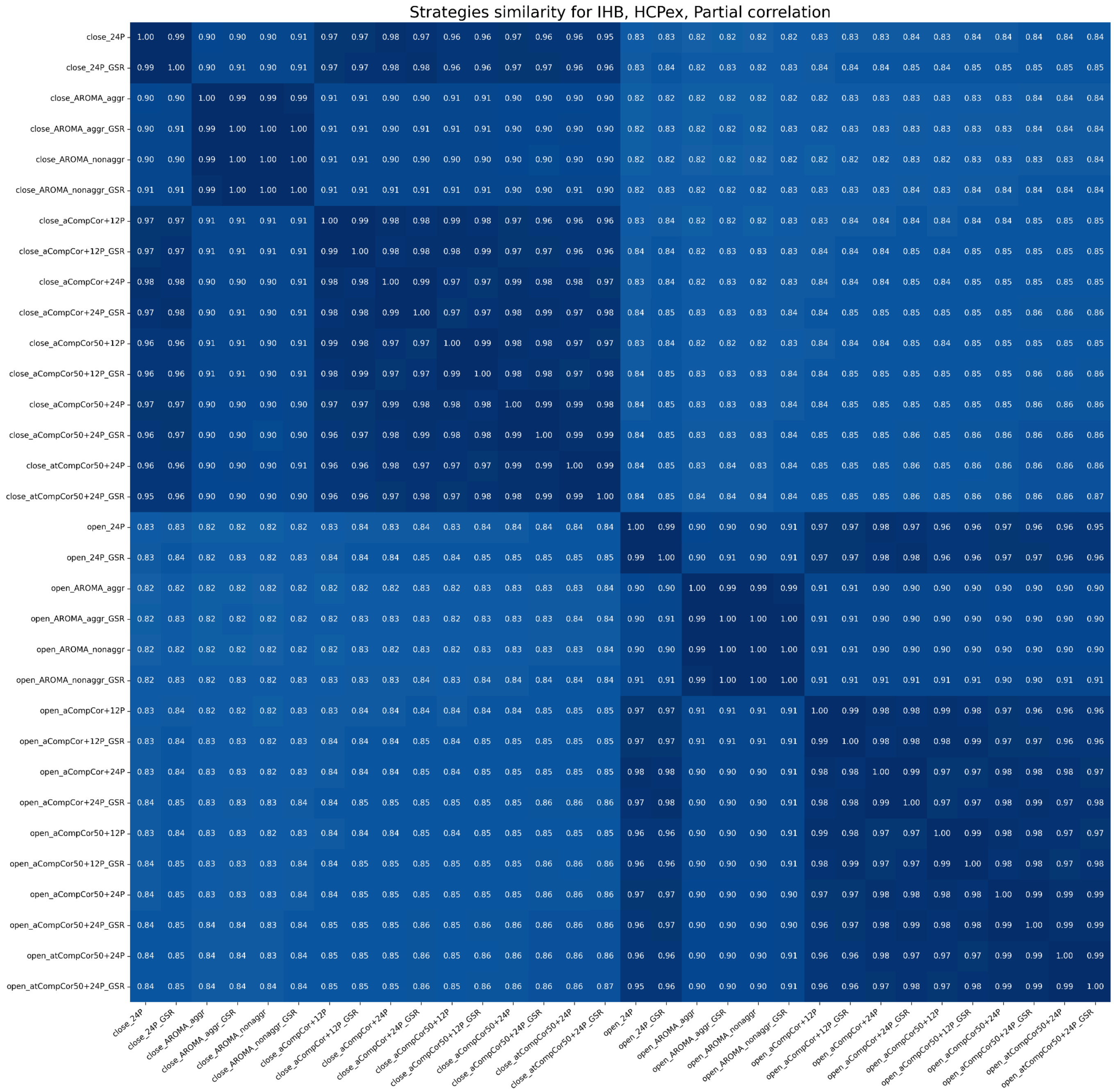


*Figure S31.*

*Similarity between signals of both EO and EC states denoised using different strategies for the IHB dataset, the HCPex atlas, partial correlation FC. The heatmap shows similarity for one FC measure. The color range is from white (0, low similarity) to dark blue (1, high similarity). Mean values across subjects are presented.*

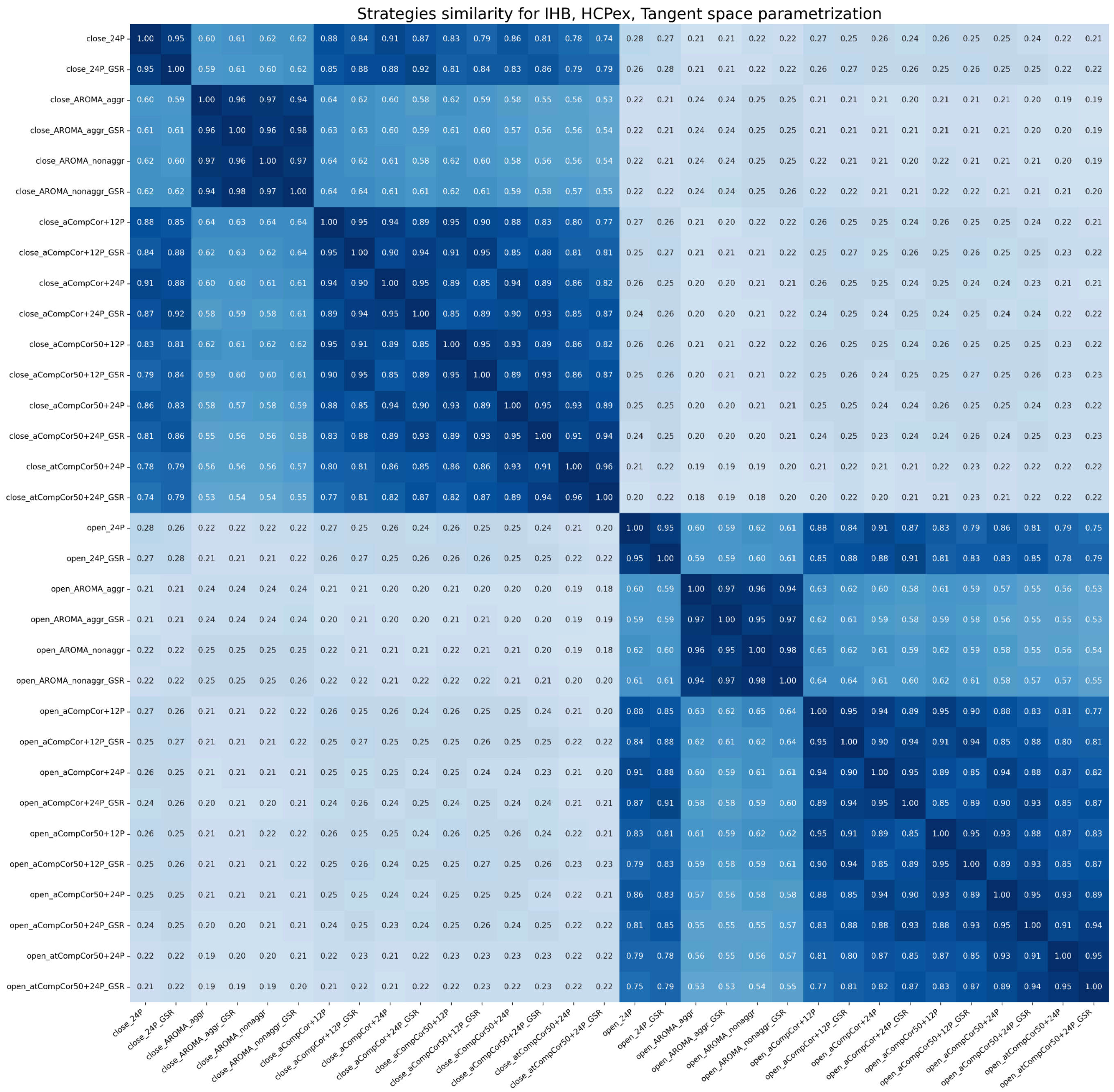


*Figure S32.*

*Similarity between signals of both EO and EC states denoised using different strategies for the IHB dataset, the HCPex atlas, tangent space FC. The heatmap shows similarity for one FC measure. The color range is from white (0, low similarity) to dark blue (1, high similarity). Mean values across subjects are presented.*

### Schaefer200

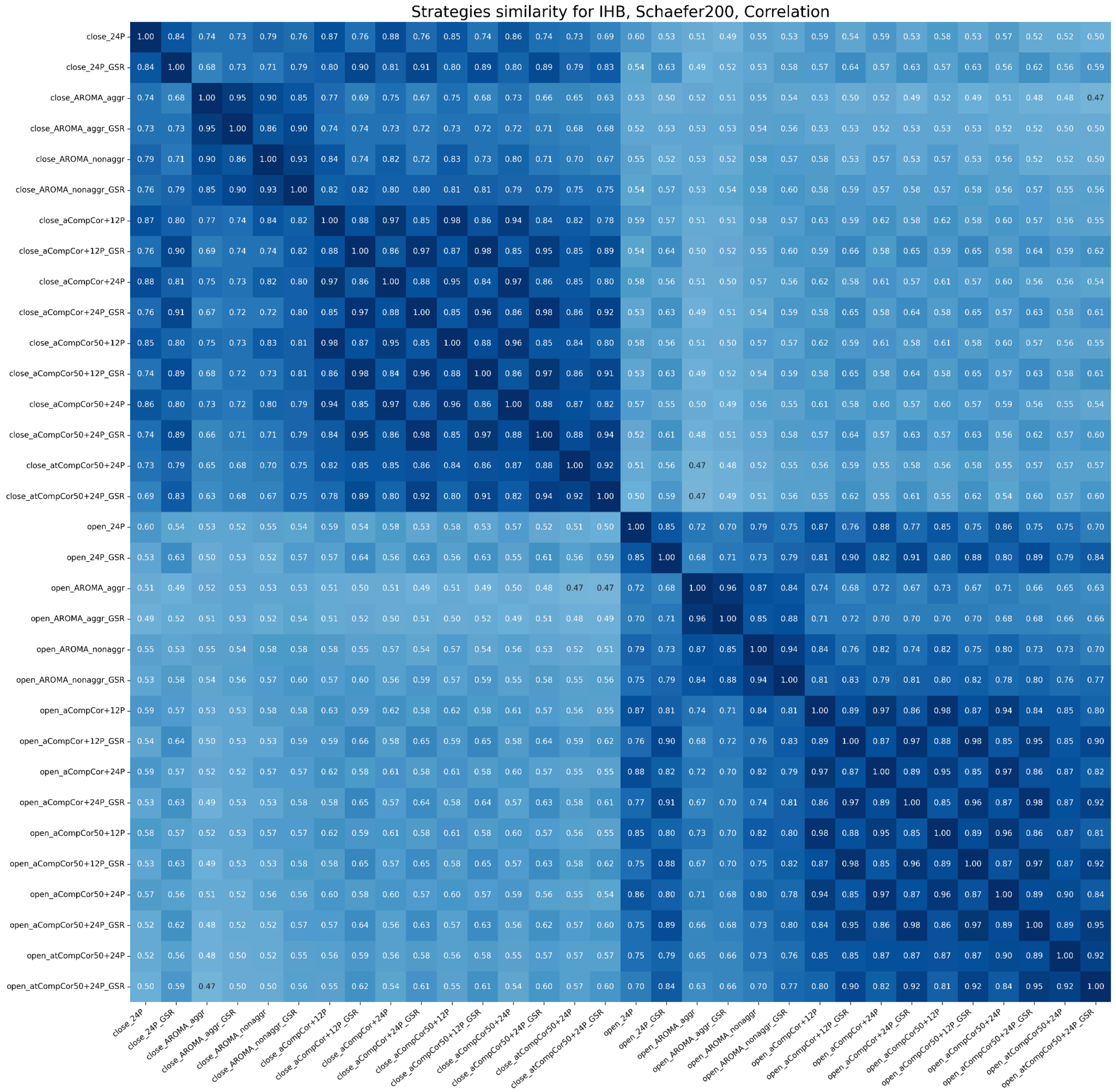


*Figure S33.*

*Similarity between signals of both EO and EC states denoised using different strategies for the IHB dataset, the Schaefer200 atlas, correlation FC. The heatmap shows similarity for one FC measure. The color range is from white (0, low similarity) to dark blue (1, high similarity). Mean values across subjects are presented.*

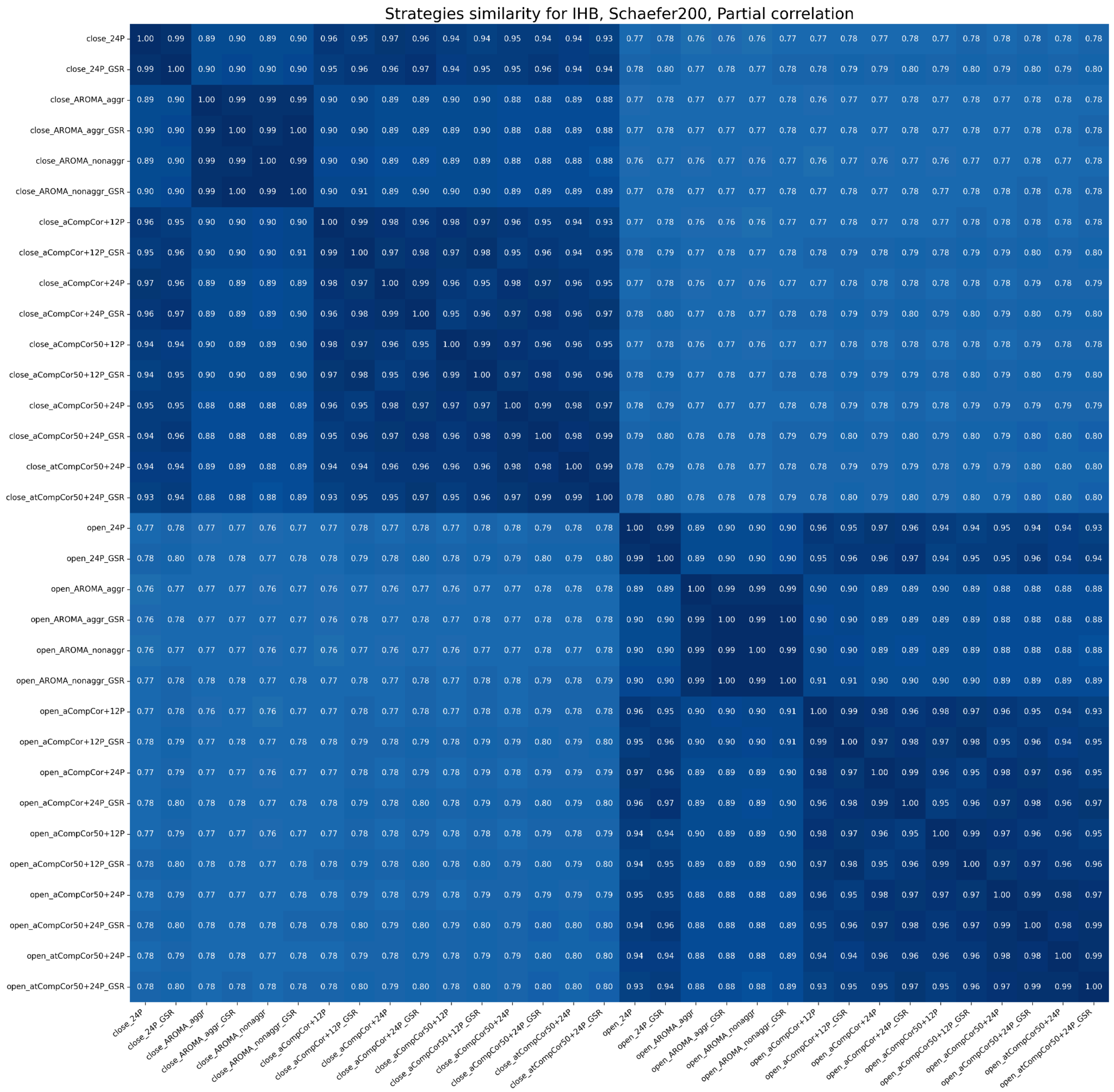


*Figure S34.*

*Similarity between signals of both EO and EC states denoised using different strategies for the IHB dataset, the Schaefer200 atlas, partial correlation FC. The heatmap shows similarity for one FC measure. The color range is from white (0, low similarity) to dark blue (1, high similarity). Mean values across subjects are presented.*

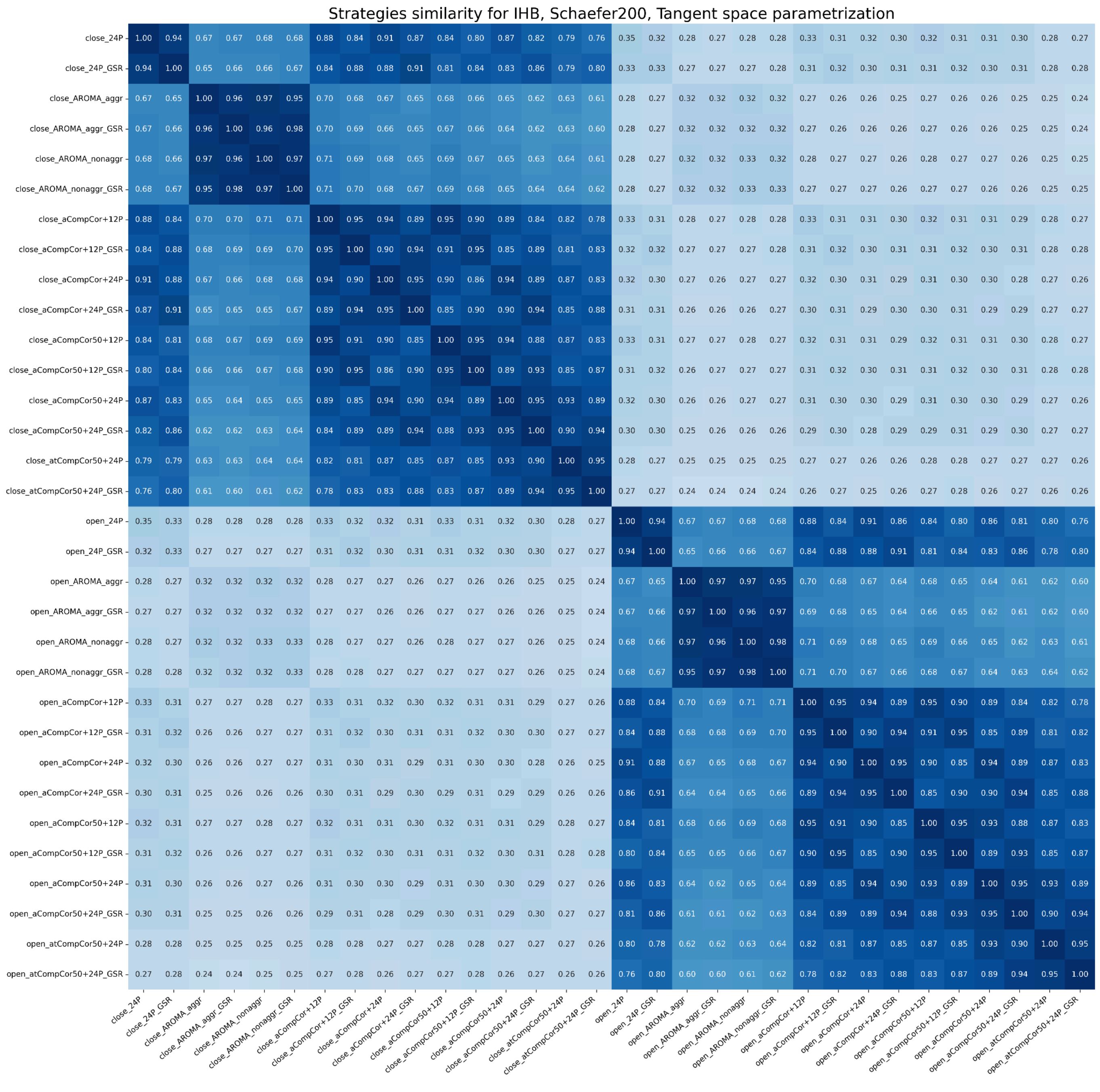


*Figure S35.*

*Similarity between signals of both EO and EC states denoised using different strategies for the IHB dataset, the Schaefer200 atlas, tangent space FC. The heatmap shows similarity for one FC measure. The color range is from white (0, low similarity) to dark blue (1, high similarity). Mean values across subjects are presented.*

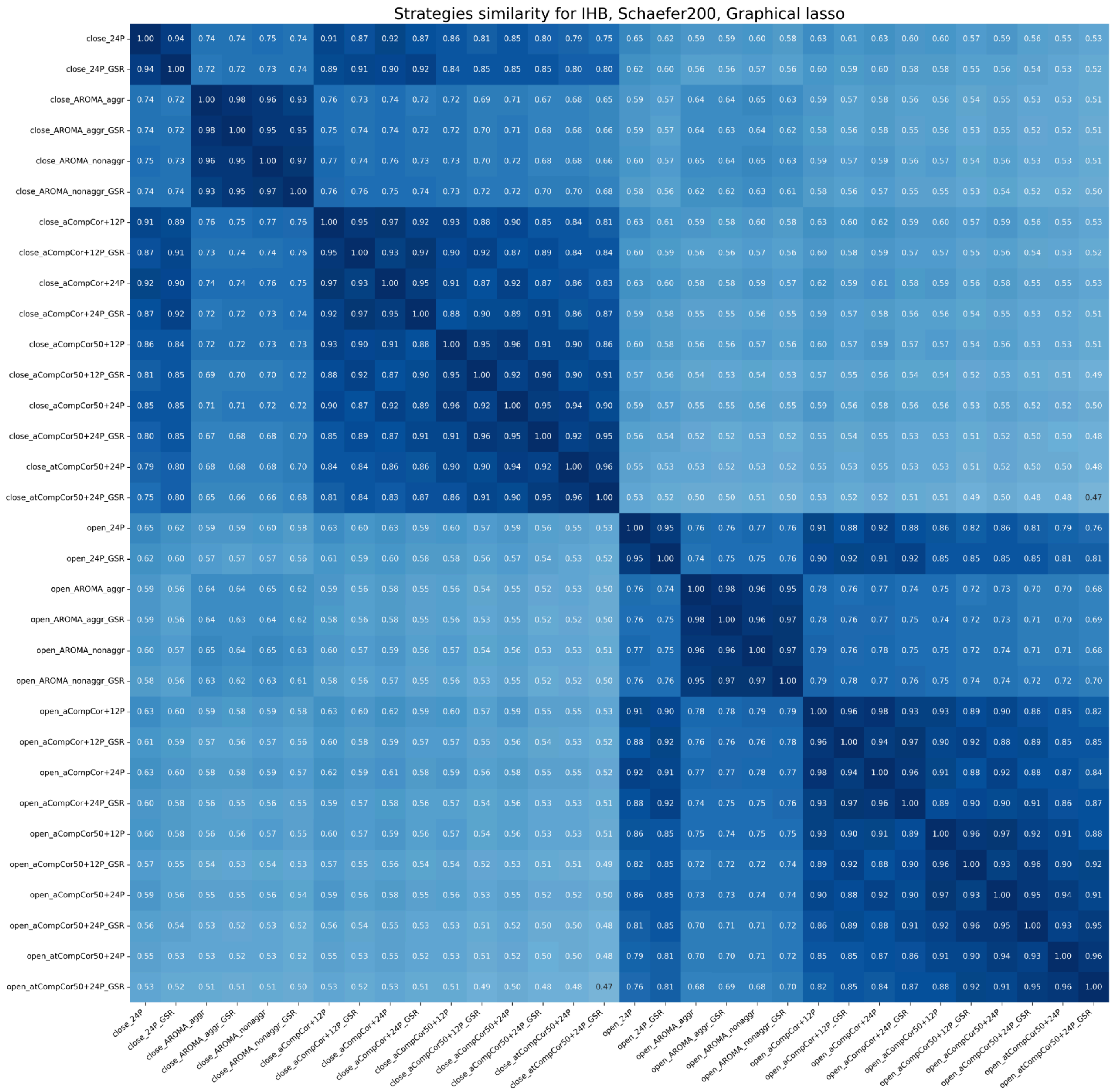


*Figure S36.*

*Similarity between signals of both EO and EC states denoised using different strategies for the IHB dataset, the Schaefer200 atlas, glasso FC. The heatmap shows similarity for one FC measure. The color range is from white (0, low similarity) to dark blue (1, high similarity). Mean values across subjects are presented.*
